# Transcription factor TGA2 is essential for UV-B stress tolerance controlling oxidative stress in Arabidopsis

**DOI:** 10.1101/2020.05.24.113589

**Authors:** Ariel Herrera-Vásquez, Alejandro Fonseca, José Manuel Ugalde, Liliana Lamig, Aldo Seguel, Tomás C. Moyano, Rodrigo A. Gutiérrez, Paula Salinas, Elena A. Vidal, Loreto Holuigue

## Abstract

Plants possess a diversity of Reactive Oxygen Species (ROS)-processing enzymes involved in sensing and controlling ROS levels under basal and stressful conditions. There is little information on the transcriptional regulators that control the expression of these ROS-processing enzymes, particularly at the onset of the defense response to abiotic stress. Filling this gap, this paper reports a critical role for Arabidopsis TGA class II factors (TGA2, TGA5, and TGA6) in the tolerance response to UV-B light and photooxidative stress, by activating the expression of genes with antioxidative roles. We identified two clusters of genes responsive to UV-B and activated by TGA2/5/6 were identified using RNAseq and clustering analysis. The *GSTU* gene family, which encodes glutathione transferase enzymes from the Tau subclass, was overrepresented in these clusters. We corroborated the TGA2-mediated activation in response to UV-B for three model genes (*GSTU7*, *GSTU8*, and *GSTU25*) using RT-qPCR and ChIP analyses. Interestingly, using *tga256* mutant and *TGA2*- and *GSTU7*-complemented mutant plants, we demonstrated that TGA2-mediated induction of *GSTU* genes is essential to control ROS levels and oxidative damage after UV-B and MeV treatments. This evidence positions TGA class II factors, particularly TGA2, as a key players in the redox signaling network of Arabidopsis plants.

**Highlight:** Arabidopsis TGA2 transcription factor is part of the redox-signaling network controlling ROS levels and oxidative damage in tolerance response to UV-B and photooxidative stress, via activation of antioxidant *GSTU* genes.

## Introduction

Plants are equipped with a diversity of genetic mechanisms to protect themselves, survive and adapt to stress caused by changes in environmental conditions and by pathogen attacks (Zhu, 2016; Diaz, 2018). Ultraviolet B light (UV-B), the electromagnetic radiation with the highest energy that reaches the earth surface from the sun, is one important source of environmental stress for plants, triggering developmental and defense genetic programs (Ulm and Jenkins, 2015; Yin and Ulm, 2017). Overexposure to UV-B radiation, like most stress conditions, induces an oxidative burst produced by increased cellular levels of Reactive Oxygen Species (ROS) (Brosché and Strid, 2003; Frohnmeyer and Staiger, 2003). Particularly under UV-B stress, ROS accumulation occurs in the apoplast, due to activation of the NADPH oxidase AtRBOHD and AtRBOHF subunits (Rao et al., 1996; Casati and Walbot, 2003; Kalbina and Strid, 2006); and in the chloroplasts, due to an effect of UV-B on photosystem (PS) II function (Kulandaivelu and Noorudeen, 1983; Larkum et al., 2001).

ROS have a dual effect in the defense response, at higher levels they produce oxidative damage of biomolecules such as lipids, proteins, and DNA (Das and Roychoudhury, 2014), while at lower levels they act as signals for the activation of defense and developmental genetic programs (Schopfer et al., 2001; Foyer and Noctor, 2013; Vaahtera et al., 2014; Considine et al., 2015; Mignolet-Spruyt et al., 2016; Noctor et al., 2018; Fichman and Mittler, 2020). Accordingly, one of the critical challenges for plants, as sessile organisms exposed to stressful environmental changes, is to sense and control ROS levels in order to properly respond to environmental signals, survive and adapt.

Plant cells possess a robust network of metabolic systems involved in sensing and controlling ROS levels (Foyer and Noctor, 2013; Farooq et al., 2019). These systems use non-enzymatic components, such as the redox buffers glutathione, ascorbate and NADPH; and enzymatic components represented by a diversity of multifunctional peroxidases, reductases, and dehydrogenases (Foyer and Noctor, 2011). These systems not only scavenge ROS keeping their levels low, but are also part of the cellular redox signaling network that transmits oxidative signals to trigger defense and developmental responses (Noctor et al., 2018). Considering this dual role, Noctor et al (2018) proposed to refer to these enzymatic systems as “ROS-processing systems”.

The main plant ROS-processing systems are the well-recognized antioxidative enzymes that directly detoxify ROS in different organelles (superoxide dismutase SOD, catalase CAT, ascorbate peroxidase APX) and that regenerate reduced forms of redox buffers (monodehydroascorbate reductase MDAR, dehydroascorbate reductase DHAR, glutathione reductase GR, and NADPH-thioredoxin reductase NTR) (Meyer et al., 2012). Supporting the robustness of ROS metabolism in plants, ROS-processing systems also include other enzymes encoded by large multigene families, such as peroxiredoxins (PRX), glutaredoxins (GRX), thioredoxins (TRX), glutathione peroxidases (GPX) and glutathione transferases (GST) (Dixon and Edwards, 2010; Meyer et al., 2012; Bela et al., 2015; Liebthal et al., 2018; Sylvestre-Gonon et al., 2019). Interestingly, these thiol-based enzymes (PRX, GRX, TRX) and peroxidases (GPX and GST) are encoded by genes highly responsive to biotic and abiotic stress, and to a wide variety of stress-associated compounds such as Salicylic Acid (SA), Jasmonic Acid (JA), Abscisic Acid and H_2_O_2_ (Meyer et al., 2012; Das and Roychoudhury, 2014; Choudhury et al., 2017). The functional roles, as well as the mechanisms and signaling pathways involved in the activation of most of these genes are still not fully known (Sewelam et al., 2016).

The TGA transcription factors are bZIP proteins that have been involved in plant responses to biotic stress and in developmental processes. The Arabidopsis genome codes for 10 *TGA* members divided in 5 classes according to their sequence similarities (Jakoby et al., 2002; Gatz, 2013). The TGA class II, that includes *TGA2*, *TGA5* and *TGA6*, show a redundant function in defense response against pathogens, being described as essential for the Systemic Acquired Response (SAR) against biotrophic pathogens (Zhang et al., 1999), and for the defense response against the necrotrophic pathogen *B. cinerea* (Zander et al., 2014) in Arabidopsis. This is due to their role as gene regulators in the SA-mediated (Zhang et al., 2003; Blanco et al., 2009) and the JA-Ethylene-mediated pathways (Zander et al., 2014), respectively. TGA class II factors have been also involved in controlling the expression of genes with detoxification functions in response to SA (Blanco et al., 2009; Herrera-Vasquez et al., 2015) and to oxylipins (Mueller et al., 2008; Stotz et al., 2013).

Current evidence supports a role for ROS as signaling molecules, and a functional role of diverse families of ROS-processing enzymes in the antioxidative defense response to stress. However, the transcriptional regulation mechanisms that link ROS signaling with the effector genes of the defense response, is poorly understood. Here, we show that TGA2/5/6 transcription factors are critical regulators of ROS-processing responses to stress. Accordingly, genetic and transcriptomic evidence supports a role for class II TGA factors, particularly TGA2, in the control of ROS levels and oxidative damage in the tolerance response to UV-B light and photooxidative stress, by regulating the transcription of a group of genes coding for ROS scavenging enzymes such as GSTU. These results indicate that TGA class II factors are key for cellular redox homeostasis, acting as essential regulators of the antioxidative response induced under stress conditions.

## Materials and methods

### Plant material and growth conditions

*Arabidopsis thaliana* wild type plants and all mutant lines were in Columbia (Col-0) background. The mutant lines used were *tga2-1/tga5-1/tga6-1* (*tga256*), *tga2-1/tga5-1* (*tga25*) and *tga6-1* (*tga6*) (Zhang et al., 2003); and *tga2-1/tga5-1/tga6-1* complemented with the *pUBQ:TGA2-V5* construct (*tga256*/TGA2 lines #1 and #2, this report), or the *pUBQ:GSTU7-V5* construct (*tga256/GSTU7* lines #1 and #2, this report). Seedlings were grown *in vitro* in 0.5X MS medium supplemented with 10 g/L sucrose and 0.3% Phytagel (Sigma) in a Percival growth chamber (16 h light, 100 µmoles m^−2^ s^−1^, 22 ± 2ºC).

### Genetic constructs and plant transformation

The *pUBQ:TGA2-V5* and *pUBQ:GSTU7-V5* constructs were generated using the Gateway technology following the manufacturer’s instructions (Invitrogen). The *TGA2* and *GSTU7* coding region was amplified from cDNA using the primers described in Table S2. The purified PCR products were cloned into the pENTR/SD/D-TOPO vector and then recombined into the pB7m34gw vector to express the proteins fused to a V5 tag controlled by the UBQ10 promoter (Grefen et al., 2010). *tga256* mutant plants were transformed by floral dip method using *Agrobacterium tumefaciens* C58 strain carring the corresponding vectors. Transgenic seeds were selected in 0.5X MS solid medium supplemented with 15 µg/ml glufosinate-ammonium. Two stable homozygous transgenic lines for each construct (indicated as #1 and #2) were used for further analyses.

### Plant stress treatments

The protocol for tolerance assays to germinate in SA is described on (Zhang et al., 2003) and we use 0.2mM SA. For SA treatments to evaluate *GRXC9* gene expression, 15-day-old seedlings were floated on 0.5 mM SA (treatment) or 0.5X MS medium as a control, and incubated for the indicated periods of time in a growth chamber.

To evaluate tolerance to UV-B irradiation, 15-day-old seedlings grown in plates with solid medium were exposed to UV-B light (0,210 mW/cm2) in a chamber equipped with two USHIO UVB F8T5.UB-V, UVP 3400401 fluorescent tubes (λ= 306 nm) during 24 h and then recovered for 72 h in the growth chamber under controlled conditions (16 h light, 100 µmoles m^−2^ s^−1^, 22 ± 2ºC). As control we used seedlings treated in the UV-B chamber and covered with a cellulose acetate polyester filter. Samples were taken at the indicated periods of time during the UV-B treatment or the recovery period to analyze fresh weight, ROS levels, and gene expression by RT-qPCR and by RNAseq. For ion leakage assays, 10-day-old seedlings floated in MilliQ water were exposed to UV-B as indicated above.

To evaluate Arabidopsis lines for tolerance to germinate in MeV, seeds were surface sterilized, spread on solid MS medium supplemented with 0.1 µM MeV, stratified at 4°C for 48 h in the dark, and then germinated and grown under standard controlled conditions. After 15 days, the number of green seedlings compared with the total number of germinated seeds (% survival) was recorded. To detect ROS production after MeV treatment, a 2 µl drop of MeV solutions (0, 15 and 30 µM) was placed on the surface of 20 leaves from different 15-day-old plants grown *in vitro*. Plants were then maintained under constant light (100 µmoles m^−2^ s^−1^) for 24 h and ROS staining was performed as described below.

### Ion leakage assays

Ion leakage was measured in seedlings subjected to UV-B irradiation for different periods of time. For each sample, three 10-day-old seedlings were floated abaxial side down onto a 12-well plate with 2 ml of MilliQ water, and incubated under UV-B as indicated above. The conductivity of the bathing solution was then measured at 22°C with a conductimeter. The seedlings with the bathing solution were introduced into sealed tubes, and sterilized by autoclaving. The bathing solution was measured again and this value was referred to as 100%. For each sample, ion leakage was expressed as percentage leakage referred to its corresponding 100%.

### ROS levels analysis

Accumulation of ROS *in situ* after UV-B was detected using DAB (3-3’ diaminobenzidine) on whole rosettes of 2-week-old seedlings treated with UV-B following the protocol described in (Daudi and O’Brien, 2012). In UV-B treatments, 16 wild type seedlings, 16 *tga256* seedlings, 14 *tga256*/TGA2 #1 seedlings and 10 *tga256*/TGA2 #2 seedlings were used. The experiment was repeated 3 times in different days. ImageJ software (Schneider et al., 2012) was used to define a threshold to distinguish the stained tissue and then to calculate the areas. Finally, the percentage of stained area with respect to the total was calculated for each seedling.

### Immunoblot assays

To evaluate the levels of TGA2-V5 and GSTU7-V5 proteins in Arabidopsis lines expressing the *pUBQ:TGA2-V5* and *pUBQ:GSTU7-V5* constructs, commercial monoclonal anti-V5 antibodies (Thermo Fisher Scientific) were used. Thirty µg of total protein of each sample were separated on 15% SDS–polyacrylamide gels and transferred to ImmobilonTM PDVF membranes (Millipore Co., Bedford, MA, USA). The membrane was blocked for 1 h in PBS-T with 5% non-fat dried milk at room temperature. To detect V5-tagged proteins a 1: 5,000 dilution of the primary anti-V5 antibody (Invitrogen, #R96025) and an anti-mouse horseradish peroxidase-conjugated secondary antibody (1:10,000 dilution, Goat anti-mouse, KPL, USA). The immunocomplexes were visualized using Thermo Scientific Pierce chemiluminiscent Western Blotting Substrate (#32109, Pierce) according to the manufacturer’s instructions.

### RT-qPCR analysis

Total RNA od whole seedlings was obtained from frozen samples using the TRIzol® Reagent (Invitrogen). cDNA was synthesized with the ImProm II Kit (Promega). qPCR was performed using the Brilliant III Ultra-Fast SYBR® Green QPCR Master mix reagents (Agilent Technologies) on a AriaMx realtime PCR system equipment. The expression levels of *GRXC9* (AT1G28480), *GSTU7* (AT2G29420), *GSTU8* (AT3G09270) and *GSTU22* (AT1G78340) were calculated relative to the *YLS8* (AT5G08290) housekeeping gene. Primers used for each gene are listed in Table S2.

### Library preparation and RNAseq analysis

Total RNA was obtained from 12 samples (UV-B-treated samples for 5 h and control UV-B-filtered samples) x (wild type and *tga2/5/6* mutant genotypes) x (three independent biological replicates), using the PureLink RNA Mini kit (Thermo Fischer Scientific, Cat 12183018A) as suggested by the manufacturer. Quality of total RNA isolated for library preparation was determined using capillary electrophoresis on a Fragment Analyzer (Advanced Analytical Technologies, Cat DNF-471). All samples used for library preparation had an RNA Quality Number (RQN) of 8 or superior. Library preparation for RNAseq was performed using the TruSeq® Stranded mRNA Library Prep Kit (Illumina, Cat. RS-122-2102), starting from 2 µg of total RNA and following the manufacturer’s instructions. Library quantitation was performed by qPCR using the Library Quant Kit Illumina GA (KAPA, Cat KK4824). The size range of the libraries was determined using the High Sensitivity NGS Kit on the Fragment Analyzer (Advanced Analytical Technologies, Cat DNF-474). The libraries were sequenced on a HiSeq2500, generating 125 bp paired end reads. Data quality was assessed with FastQC 0.11.6 (https://www.bioinformatics.babraham.ac.uk/projects/fastqc/) and Trimmomatic v. 0.36 (Bolger et al., 2014) was used to remove low quality reads, using the following settings: two mismatches allowed between adapter and seed sequence, minimum of Q=30 for palindromic alignment between adapter and sequence, minimum of Q=10 for simple alignment between adapter and sequence, removal of leading and trailing sequences if Q<3, sliding window of 4 bases and removal of sequence if Q<15. The sequences were mapped to the TAIR10 Arabidopsis genome using TopHat2. Read counts per genomic feature (TAIR10 annotation) were determined for each library using the featureCounts function of the Rsubread R library (McCarthy et al., 2012; Liao et al., 2013). Data was median normalized using the R library EBSeq (Leng and Kendziorski, 2019) and normalized and log2 transformed-data was subjected to a two-way ANOVA analysis, with a false discovery rate of 5%. For the ANOVA, we used a model considering the expression of a given gene Y as Yi = β0 + β1T + β2G + β3TG + ε, where β0 is the global mean; β1, β2, and β3 are the effects of T, G, and the TG interaction; and the variable ε is the unexplained variance.

### Clusters and gene ontology analysis

The data was normalized transforming the values by using the mean and the standard deviation of the row of the matrix to which the value belongs, using the z-score formula: Value = [(Value) – Mean(Row)]/[Standard deviation(Row)]. Normalized expression data of genes belonging to the TG regulated group was used to generate a hierarchical clustering of genes using Pearson correlation and average linkage method using the Multiple experiment Viewer analysis software (Howe et al., 2011). Gene Ontology analyses for overrepresentation of Biological processes and Molecular function terms were performed using the BioMaps tool at the VirtualPlant webpage (Katari et al., 2010), using as background the TAIR 10 genome, GO assignments by TAIR/TIGR, Fisher exact test and p<0.05.

### Peroxidase activity assays

The glutathione peroxidase activity of total protein extracts from wild type, *tga256* plants and *tga256*/TGA2 #1 complemented plants was measured as the velocity of decrease in Absorbance (=340) in 500 l of reaction mixture [50 mM phosphate buffer pH 7.0, 1 mM GSH, 10 mM H_2_O_2_, 0.15 mM NADPH and 5U glutathione reductase (Sigma, G3664)] using 50 g of protein extract. The activity was calculated as (Abs=340/t)/mg of protein on the first 180 sec of reaction while the Pearson correlation factor (R2) was higher than 0.98. For determination of NADPH concentration, NADPH(340)=6150mol^−1^cm^−1^ was used. For each reaction, the spontaneous degradation of NADPH was measured previous to protein extract incorporation into the reaction mixture and was subtracted from the measurements with the protein extract incorporated.

### Chromatin immunoprecipitation assay

ChIP assays were performed and analyzed as described (Saleh et al., 2008), using 3 g of fresh leaf tissue per sample. Five microliters of the following antibodies were used for immunoprecipitation assays: anti V5 antibody (Invitrogen, #R96025), and purified IgG (A2609, Santa Cruz Biotechnology) used as a nonspecific antibody control. The concentration of DNA in each sample (input chromatin and chromatin immunoprecipitated with either specific or nonspecific antibodies) was quantified by qPCR, using the Stratagene MX3000P® equipment and the Sensimix Plus SYBRGreen Reagents (Quantece). Primers used to amplify the promoter region containing the TGA elements are listed in Table S2.

### Accession numbers

*TGA2* (AT5G06950), *GRXC9* (AT1G28480), *CHS* (AT5G13930), *YLS8* (AT5G08290), *GSTU7* (AT2G29420), *GSTU8* (AT3G09270), *GST25* (AT1G17180), *ACT2* (AT3G18780)

### Data statement

The large-scale datasets can be found in the Sequence Read Archive (SRA) submission: SUB2037222 Transcriptomics analysis of the response of wild-type and tga2/5/6 Arabidopsis seedlings to UV-B, Oct 26 ’16 https://www.ncbi.nlm.nih.gov/sra/?term=PRJNA352413

## Results

### TGA class II factors are essential for tolerance to UV-B stress

To evaluate the role of class II TGAs in the defense response to stress in Arabidopsis, we used the previously characterized *tga2-1 tga5-1 tga6-1* (*tga256)* line that carries mutations in the three class II TGA genes: *TGA2* (AT5G06950), *TGA5* (AT5G06960) and *TGA6* (AT3G12250) (Zhang et al., 2003). A reduced tolerance to germinate in high levels of SA was previously reported for the *tga256* mutant plants, but not for the double *tga2-1 tga5-1* (*tga25*) or the single *tga6-1* (*tga6*) mutant plants, indicating a redundant role for the TGA2, TGA5 and TGA6 factors in response to SA (Zhang et al., 2003). As a genetic tool to evaluate the role of TGA2 under experimental conditions, we transformed *tga256* triple mutant with the *pUBQ:TGA2-V5* gene. Expression of the V5-tagged TGA2 factor, which was detected by immunoblot using an anti-V5 antibody in the two complemented lines selected (Fig. S1A), was enough to recover previously described phenotypes of the *tga256* mutant, such as the tolerance to germinate in MS medium supplemented with 0.2 mM SA (Fig. S1B) and the SA-controlled expression of the *GRXC9* gene, which is detected in wild type plants and abolished in the *tga256* mutant (Fig. S1C). The *GRXC9* gene (AT1G28480) encodes the glutaredoxin C9 enzyme and is a marker for the group of genes that are transiently induced by SA via an NPR1-independent and TGA class II-dependent pathway (Blanco et al., 2009; Herrera-Vasquez et al., 2015). These results indicate that the V5-tagged TGA2 factor is functional and its overexpression in the *tga256* mutant background is enough to complement SA-mediated responses that are abolished in the mutant.

To evaluate the role of class II TGAs in the defense response against UV-B stress, the tolerance to UV-B treatment was assayed in the *tga256* mutant and in the two *tga256*/TGA2 complemented lines. Seedlings were exposed to stress by UV-B radiation during 24 h followed by a recovery period of 72 h under standard growing conditions. As shown in Fig. 1, plants treated with UV-B show higher chlorosis and reduced plant fresh weight, compared to the control plants. The *tga256* mutant plants were more susceptible than wild type plants to UV-B treatment, a phenotype that was reverted in the two complemented lines (completely in line #1 and partially in line #2). To analyze the participation of TGA5 and TGA6 in the defense response to UV-B, we evaluated tolerance to UV-B in the *tga2-1 tga5-1* (*tga25*) double mutant as well as in the single *tga6-1* (*tga6*) mutant (Zhang et al., 2003). As shown in Fig. S2, plants from *tga25* and *tga6* mutant lines do not show higher susceptibility to UV-B than wild type plants. Therefore, the three TGA class II genes must be mutated in order for the plant to develop a susceptible phenotype, indicating that the three genes are redundant in this response, as was previously described for the responses to SA (Zhang et al., 2003). Interestingly, the overexpression of TGA2 is enough to complement the triple mutant and activate the stress tolerance.

**Fig. 1.**
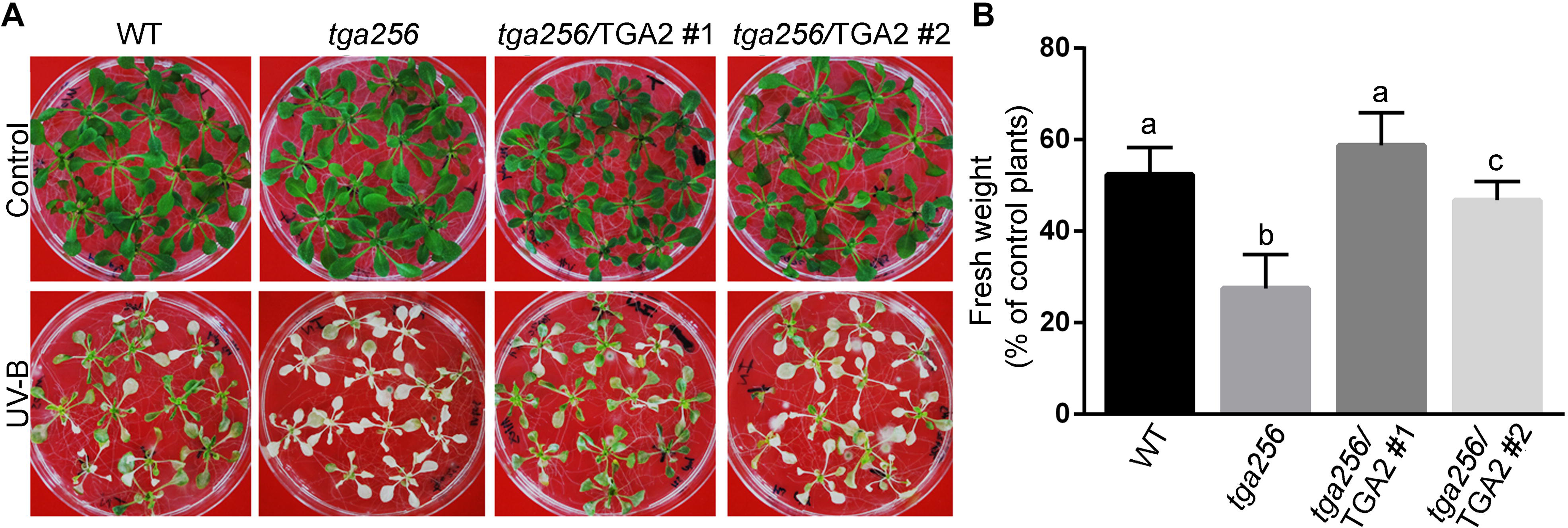
TGA class II factors are essential for tolerance to UV-B stress. Fifteen day-old seedlings of the indicated genotypes were treated with UV-B radiation for 24 h and then they recovered for 72 h in a growth chamber. Control treatments were performed under the same conditions with a UV-B filter. Pictures (**A**) and fresh weight measurements (**B**) were obtained at the end of the recovery period. Fresh weight of rosette tissue from UV-B treated plants was expressed as percentage of fresh weight of rosettes from control plants. Bars represent the mean ± standard deviation of at least 3 independent experiments (20 seedlings per genotype for each experiment). Statistical analysis was performed using ANOVA/Fisher’s LSD test. Different letters denote statistically significant differences at p<0.05.

### Identification of UV-B regulated genes controlled by TGA class II factors

In order to analyze the role of TGA class II in the genetic response to UV-B, we used RNAseq to analyze global expression profiles in wild type and *tga256* plants exposed to UV-B light. We exposed fifteen-day-old seedlings of wild type and *tga256* genotypes to UV-B for 5 hours and then total RNA was isolated from complete seedlings. In parallel, we performed control UV-B treatments in plants covered with a filter.

As a control for the response to UV-B treatment, we performed RT-qPCR to analyze the expression levels of the chalcone synthase gene (*CHS*, AT5G13930), a marker gene for this response (Fuglevand et al., 1996; Jenkins et al., 2001) (Fig. S3A). Furthermore, as a control for the *tga256* genotype, we analyzed expression data of the *PR-1* gene, which is not induced by UV-B at 5 hpt, but its expression is elevated in the *tga256* mutant due to basal repression mediated by the TGA2/5/6 factors (Zhang et al., 2003) (Fig. S3B).

Differential gene expression analysis was assessed using a two-way ANOVA analysis (p<0.01). The resulting number of genes that are differentially expressed after UV-B treatment (T), in the different genotypes (G), or as a result of the interaction between treatment and genotype factors (TG), are compared in Fig. S4.

We focused on the UV-B-responsive genes regulated by TGA class II factors, whose expression is affected by the treatment and genotype interaction (TG, 717 genes in total, see list in Table S1). These genes were clustered according to their patterns of gene expression using hierarchical clustering. We used a figure of merit analysis to determine the optimal number of different clusters that represent the major patterns of expression of TG genes. As shown in Fig. 2, we found five different clusters for which we analyzed the over-representation of Gene Ontology (GO) terms (Biological process and Molecular function). Two of the clusters include genes that are up regulated by UV-B in WT plants and whose UV-B induction is either completely abolished (cluster 1, 76 genes) or diminished (cluster 3, 124 genes) in the *tga256* mutant (Fig. 2). As expected, GO terms associated with defense to stress are over-represented in these two clusters. Another two clusters include genes down regulated by UV-B in WT plants, whose expression is affected in the *tga256* mutant only under control conditions, being negatively (cluster 2, 138 genes) or positively (cluster 5, 229 genes) regulated by TGA2/5/6 factors (Fig. 2). GO terms associated with photosynthesis (cluster 2), and growth and developmental processes (cluster 5), were over-represented in these clusters (Fig. 2), providing evidence for a negative influence of stress on plant fitness (Heidel et al., 2004; Kempel et al., 2011; Huot et al., 2014). Finally, one cluster (cluster 4, 150 genes) includes genes that are slightly up regulated by UV-B and more strongly up regulated by UV-B in *tga256* plants, suggesting a negative UV-B response control by TGA2/5/6. Significant GO terms were not found in this group of genes (Fig. 2).

**Fig. 2.**
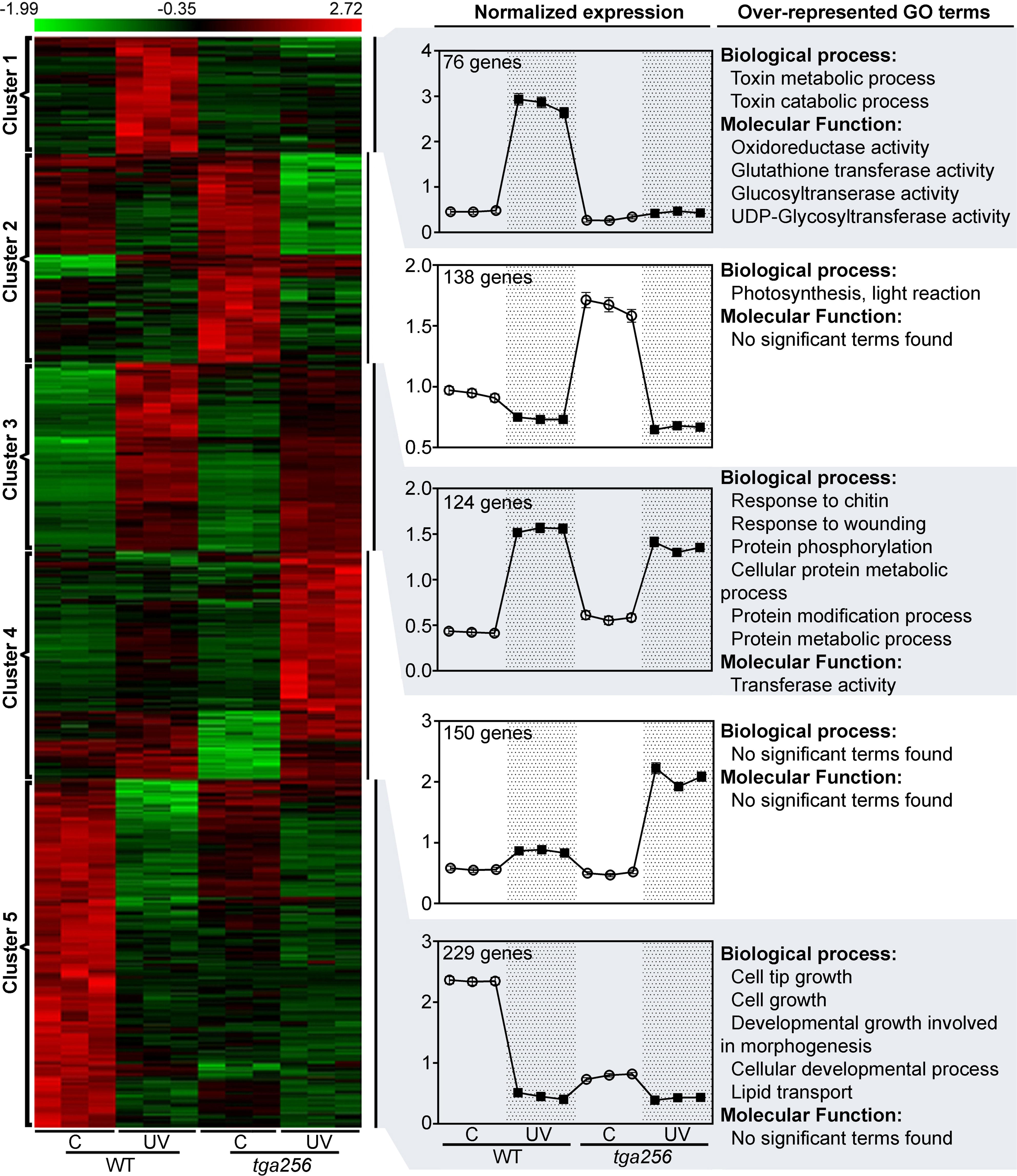
Expression patterns of UV-B responsive genes regulated by TGA2/5/6 factors. The 717 genes differentially expressed in response to treatment and genotype were grouped hierarchically. The normalized expression data of these genes in three biological replicates from wild type (WT) and *tga256* mutant plants treated under control (C) or UV-B (UV) conditions are shown. Genes were separated in 5 different clusters according to their expression profile patterns. The z-score of normalized expression for each group of genes under control (C, open circles) and UV-B conditions (UV, black squares), is shown in the graphs. Error bars represent standard error. The total number of genes is indicated at the upper left side of each graph. The over-represented GO terms were obtained according to the MIPS classification using BioMapstool from VirtualPlant (http://virtualplant.bio.nyu.edu), which compares the number of genes from each GO term in each cluster with the whole Arabidopsis genome. A Fisher Exact Test with FDR correction was used for the analysis (p<0.05).

Looking for genes that could be responsible for the phenotype of higher susceptibility to UV-B observed in the triple *tga256* mutant compared to wild type plants (Fig. 1), we focused on clusters 1 and 3 (Fig. 2). These clusters include stress defense genes up regulated by UV-B in a TGA2/5/6-dependent manner. Interestingly, over-representation of GO terms in these clusters, particularly cluster 1, suggests that class II TGAs are required for full activation of antioxidant and detoxifying capacity of the plant after stress. Accordingly, we found enrichment of genes with molecular functions of oxidoreductase and glutathione transferase activity in cluster 1, and also several *GST* genes in cluster 3 (Table S1). Moreover, we found that 41 out of the 48 *GST* genes detected in the RNAseq experiment were regulated by UV-B treatment, 33 of them being induced (underlined in Fig. 3A, dark gray circle in Fig. 3B). Of these UV-B induced *GST* genes, 20 had an altered UV-B response in the *tga256* genotype compared to wild type (light gray circle in Fig. 3B). Interestingly, 91.7% of the *GST* genes induced by UV-B treatment in a TGA class II dependent manner, belong to the *GST* subfamily Tau (*GSTU*) (dashed area in Fig. 3B). For these genes, there is a significant reduction in the UV-B-induced expression in *tga256* compared with wild type plants. This reduction is complete for *GSTU1*, *GSTU2*, *GSTU7*, *GSTU8*, *GSTU22, GSTU24* and *GSTU25*, and is partial for *GSTU13*, *GSTU17*, *GSTU19, GSTU28* and *GSTF8* genes (Fig. 3C). In the cases of *GSTU7*, *GSTU8* and *GSTU28*, basal expression was also decreased in the *tga256* mutant. We selected three *GSTU* genes that showed complete dependency of TGA class II factors (*GSTU7*, *GSTU8* and *GSTU25*) and we confirmed by RT-qPCR that the UV-B-induced expression of these genes, and the basal expression of *GSTU7* and *GSTU8*, had been restored in the *tga256*/TGA2 plants, indicating that TGA2 is required for basal and UV-B induced expression of these *GSTU* genes (Fig. 4A).

**Fig. 3.**
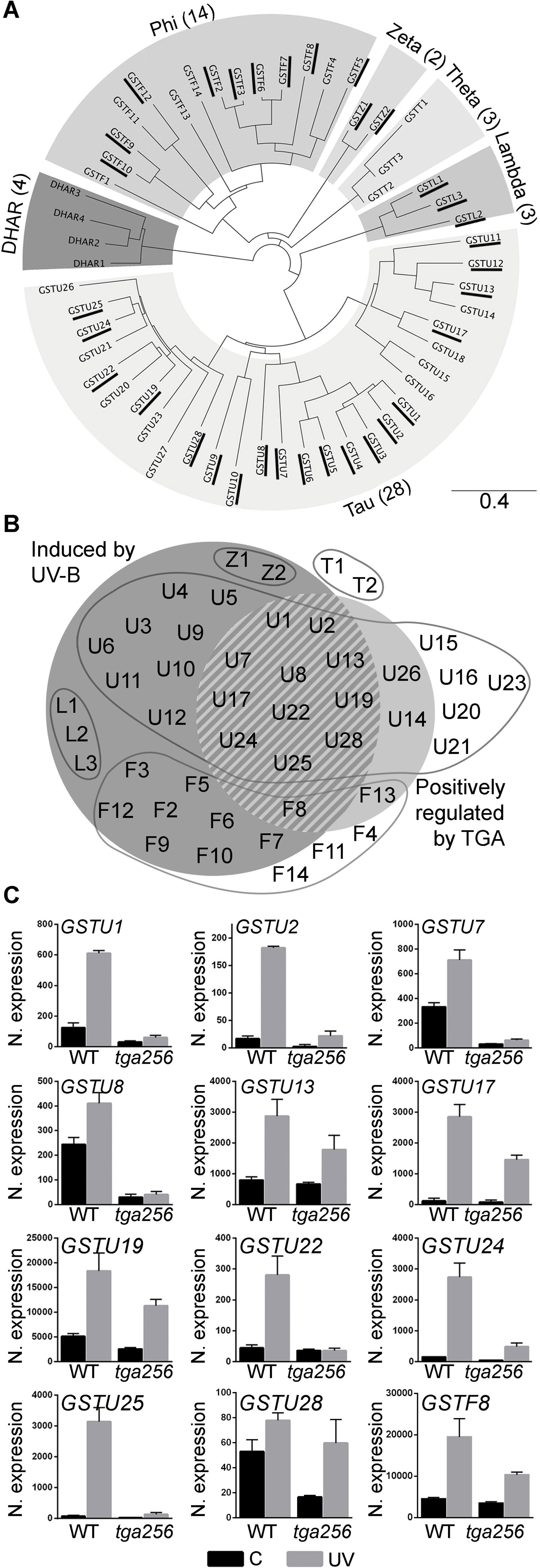
The induction of *GSTU* genes by UV-B stress is impaired in the *tga256* mutant plants. **(A)** Phylogenetic tree of the GST protein family from Arabidopsis. The tree is based on the alignment of GST protein sequences and calculated using the neighbor joining method. The six GST subfamilies described (Moons, 2005) are indicated. The underlined *GSTs* genes are positively regulated by UV-B treatment (ANOVA, p<0.05). **(B)** Venn’s diagram of the *GSTs* genes detected in the RNAseq experiments. *GSTs* genes, named according to (Moons, 2005), were grouped considering their responsiveness to UV-B treatment (dark gray circle) and their positive regulation by TGA class II factors (light gray circle) (ANOVA/Fisher’s LSD test). Continuous lines circle different GST families. **(C)** Expression levels of *GST* genes that are significantly induced under UV-B treatment and decreased in *tga256* mutant plants (dashed area in B). Results are presented as normalized expression of each *GSTU* gene from RNAseq data in wild type plants (WT) and in *tga256* triple mutant plants (*tga256*), subjected to UV-B treatment (UV, gray bars) or control conditions (C, black bars). Bars represent the mean ± standard deviation from three biological replicates.

**Fig. 4.**
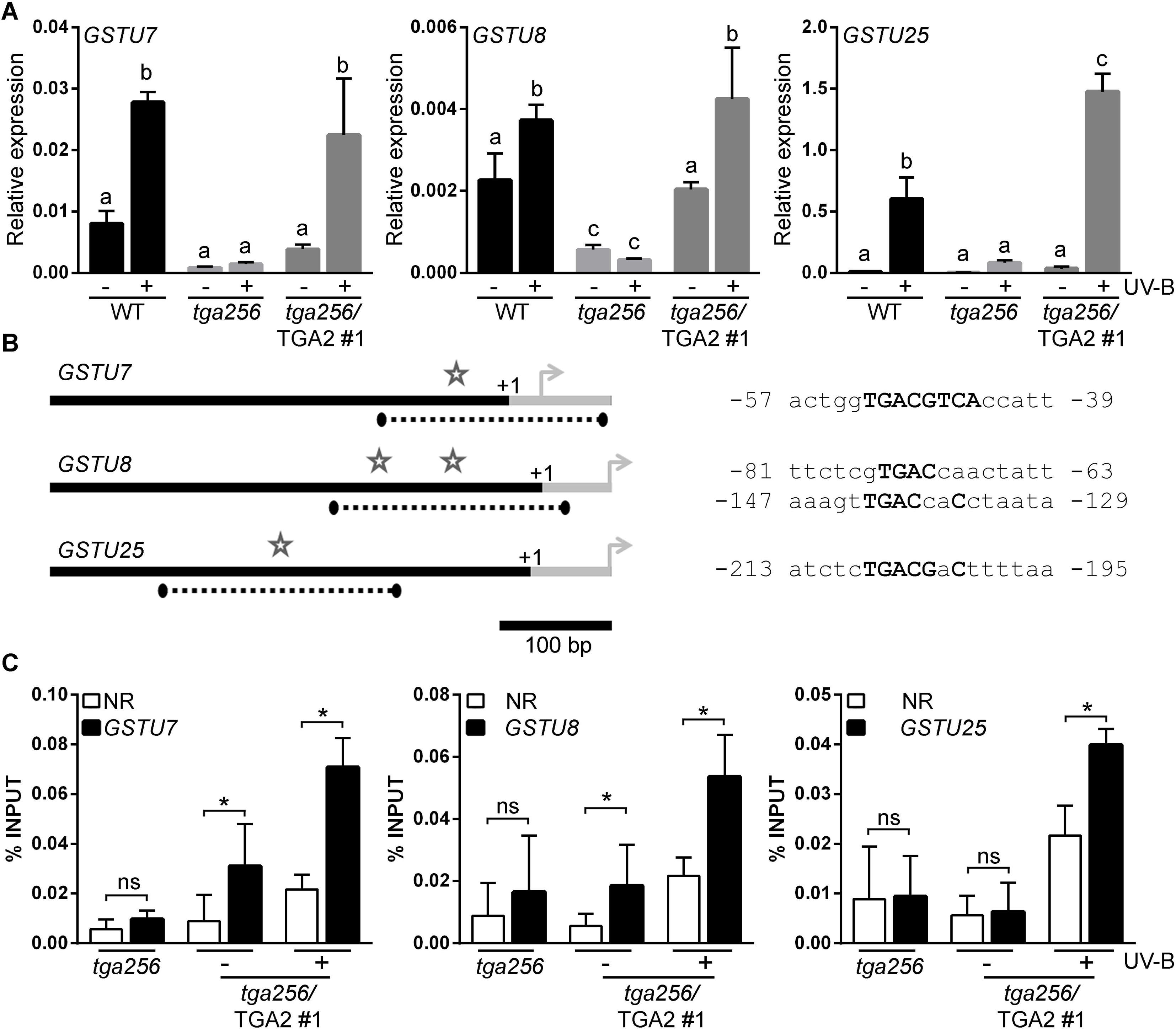
TGA2 controls the UV-B-induced expression of *GSTU7, GSTU8 and GSTU25* genes, via recruitment to their promoters. Wild type (WT), *tga256* triple mutant (*tga256*) and complemented plants (*tga256*/TGA2 line #1) were irradiated with UV-B for 5 hours (+). Untreated plants were used as control (-). **(A)** The expression of *GSTU7*, *GSTU8*, and *GSTU25* genes was evaluated by RT-qPCR. Data is presented as mean values of *GSTU* gene expression relative to the expression of the housekeeping *YLS8* gene. Error bars represent standard deviation from three biological replicates (4-5 seedlings each). Different letters above bars indicate significant differences (ANOVA/Fisher’s LSD test, p<0.05). **(B–C)** The *in vivo* binding of TGA2-V5 to the *GSTU7, GSTU8* and *GSTU25* gene promoters was evaluated by ChIP assays. **(B)** Structure of the proximal promoter region of the genes evaluated. Arrows indicate the position of the translation start codon; +1 indicates the transcription start site and the gray bars indicate the transcribed regions. Dotted lines indicate the amplified regions in the qPCR reaction. The stars indicate the position of the putative TGA binding sites. Bar indicates the length of 100 bp. The locations of the putative TGA binding sequences are shown (**B**, right panel). The numbers indicate distance in bp from the transcription start site. **(C)** The *in vivo* binding of TGA2-V5 to the *GST* promoters was evaluated in *tga256*/TGA2 #1 complemented plants, using the anti-V5 antibody. *tga256* mutant plants were used as negative controls for immunoprecipitation. The proximal promoter region of *GSTU7*, *GSTU8* and *GSTU25* were evaluated by qPCR. The coding region of the *ACTIN2* gene, which does not contain TGA binding elements, was used as a non-related region (NR). The values of immunoprecipitated DNA samples were expressed as the percentage of a non-immunoprecipitated sample (%INPUT). Error bars represent the mean ± standard deviation of three biological replicates. Asterisks indicate significant differences between binding of TGA2 to the proximal region of the *GSTU* gene evaluated and the non-related zone in each condition (ANOVA/Fisher’s LSD test, p<0.05, ns: non-significant).

The *GSTU7* and *GSTU25* genes contain a TGACG motif in the proximal promoter region (Mueller et al., 2008) and the *GSTU8* gene contains two TGA-like motifs in its proximal promoter (Fig. 4B). To evaluate whether TGA2 binds *in vivo* to these TGACG-containing regions, we performed ChIP assays in UV-B-irradiated and non-irradiated *tga256*/TGA2 plants using an anti-V5 antibody. Enrichment in TGA2 recruitment to the *GSTs* promoters was evaluated by comparison to a non-related region. Recruitment of TGA2-V5 to the promoter of these three *GSTs* genes was increased when the plants were irradiated with UV-B light (Fig. 4C). We also detected recruitment of TGA2 to the *GSTU7* and *GSTU8* promoters under basal conditions, consistent with the expression profile of these genes (Fig. 4A). This result indicates that TGA2 not only controls UV-B-induced expression of *GSTU7*, *GSTU8* and *GSTU25*, but also that it controls basal expression of *GSTU7* and *GSTU8*.

Some GST enzymes possess glutathione peroxidase (GPX) activity, being therefore part of the ROS-scavenging activity in plants (Roxas et al., 1997; Cummins et al., 1999; Dixon et al., 2009). In this context, we evaluated whether differences in *GSTs* expression - due to the lack or gain of TGA2 function - correlate with differences at the level of total GPX activity. Therefore, we assayed GPX activity using H_2_O_2_ as a substrate in total protein extracts from wild type, *tga256* and *tga256*/TGA2 plants, both under control and UV-B treatments. In wild type plants, UV-B irradiation induced an increase in GPX activity, an effect that was lost in the *tga256* mutant plants, and recovered in the TGA2 complemented plants (Fig. 5).

**Fig. 5:**
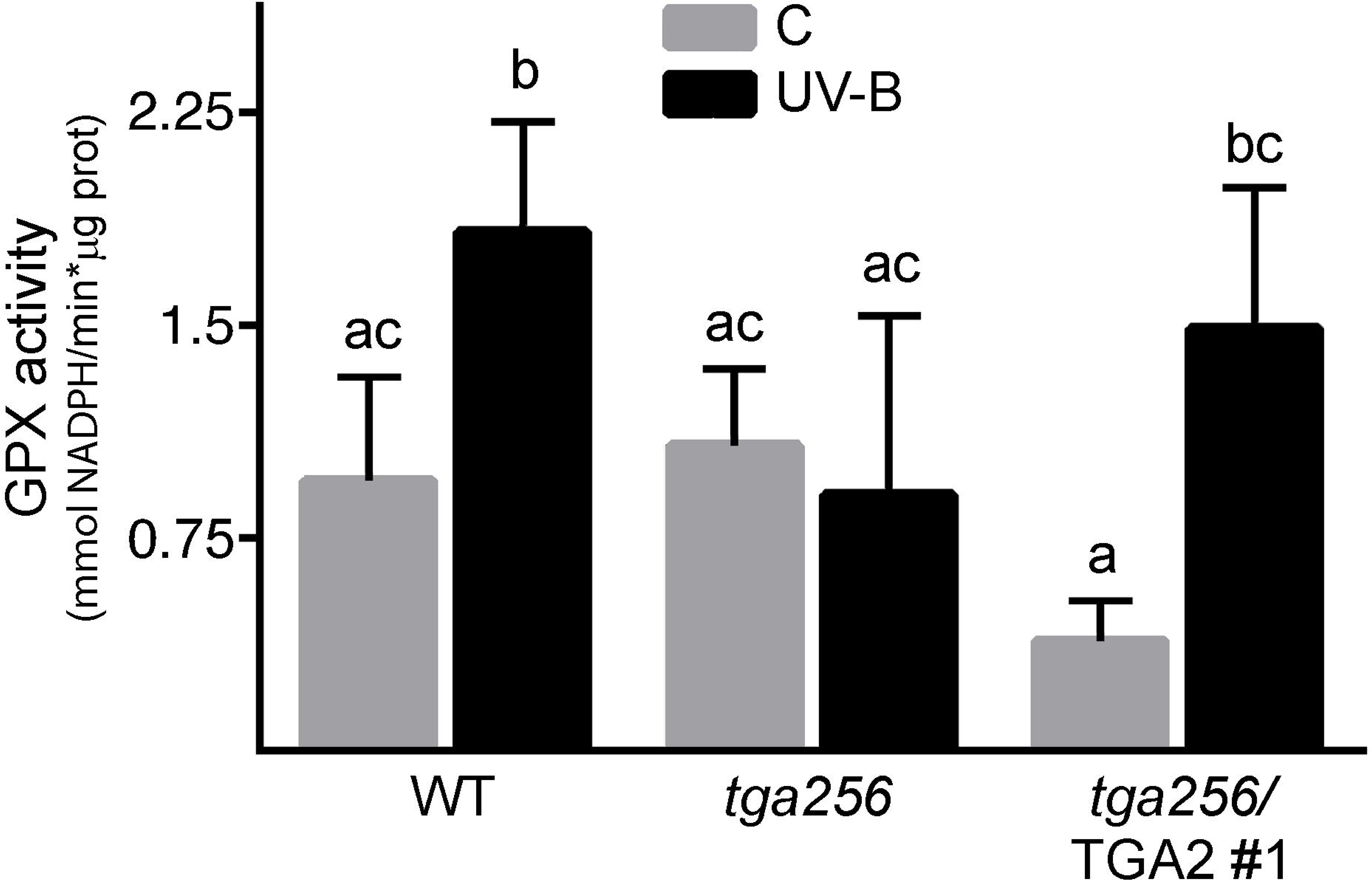
Increase in glutathione peroxidase (GPX) activity upon UV-B treatment in wild type plants is abolished in *tga256* mutant plants and recovered in *tga256*/TGA2 plants. Wild type plants (WT), the *tga256* triple mutant (*tga256*) and the complemented line (*tga256*/TGA2 #1) were irradiated with UV-B for 24 hours (gray bars) or maintained under UV-B-filtered control conditions (black bars). Total protein extracts were obtained from treated and untreated plants and the *in vitro* GPX activity was measured. GPX activity is expressed as NADPH consumption considering reaction time and protein concentration (mmol NADPH*min^−1^*µg of protein^−1^). Values from base lines, given by spontaneous NADPH degradation, were subtracted from enzyme activity values. Data is presented as mean values of GPX activity. Error bars represent the standard deviation from three biological replicates. Each extract sample was prepared using 1g of seedlings. Different letters above bars indicate significant differences (ANOVA/Fisher’s LSD test, p<0.05).

Together, these data indicate that TGA class II factors are essential for UV-B stress-induced expression of a group of *GST* genes and, accordingly, for increased peroxide-scavenging activity under this stress condition.

### TGA class II factors are essential for ROS containment in plants exposed to UV-B light and photooxidative stress

Considering that *tga256* mutant plants show a substantial reduction in the UV-B-induced expression of *GSTs* genes and a concomitant reduction in the peroxide-scavenging activity, we evaluated whether this mutant has an altered redox response to UV-B stress. For this, we measured oxidative damage and ROS levels after UV-B-stress treatment in seedlings from wild type, *tga256* mutant and *tga256*/TGA2 complemented lines. Oxidative membrane damage was measured by ion leakage assays at different periods of time during and after UV-B treatment. Increased ion leakage in the *tga256* mutant compared with wild type plants was detected during the recovery period, effect that was complemented in the TGA2-V5-expressing lines (Fig. 6A). ROS levels accumulated after UV-B treatment were also quantified in plants of all genotypes by *in situ* staining using 3-3’ diaminobenzidine (DAB) (Daudi and O’Brien, 2012) (Fig. 6B-C). Increased ROS levels were detected after UV-B treatment in *tga256* mutants compared to wild type plants, while expression of TGA2-V5 complemented this phenotype reducing ROS levels up to wild type levels (Fig. 6B-C).

**Fig. 6.**
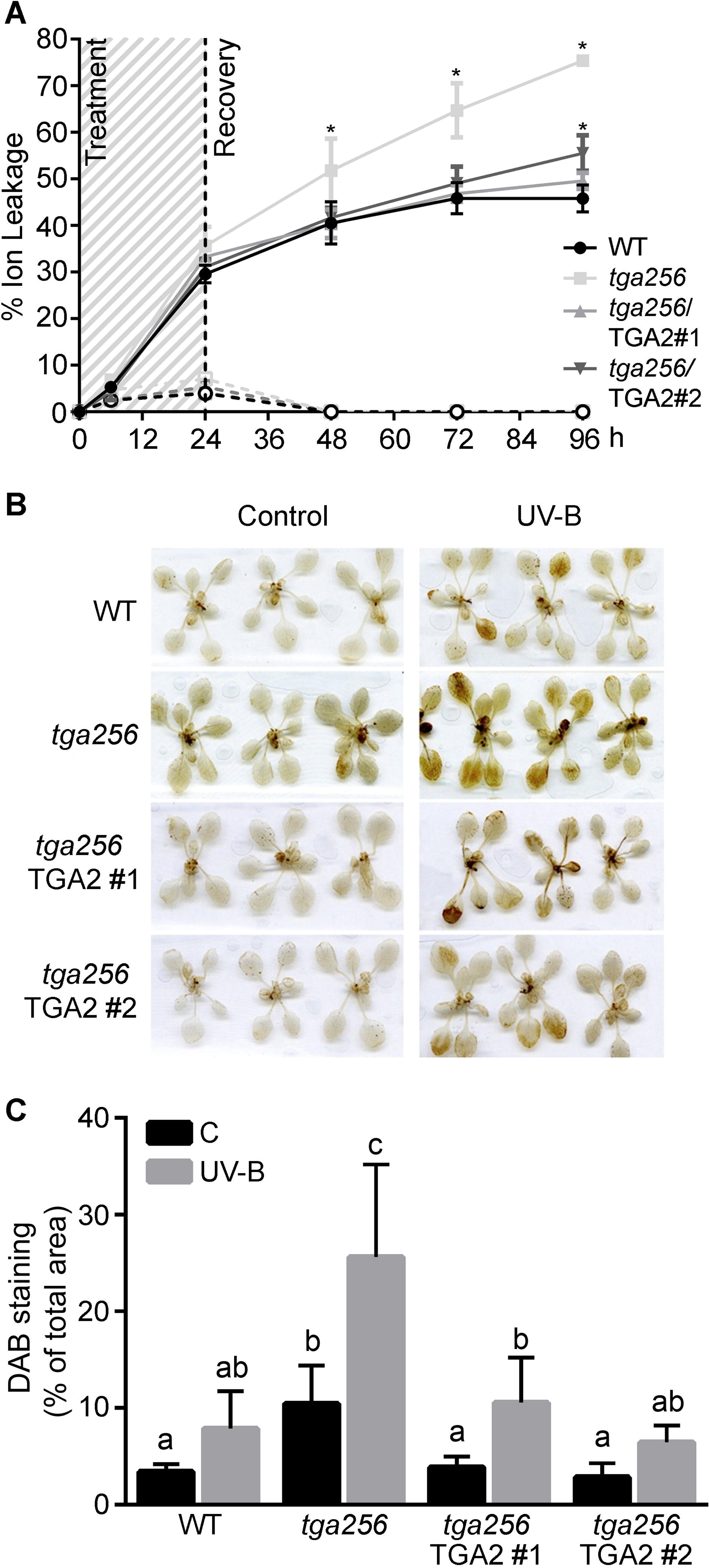
TGA2 plays a critical role in containment of oxidative damage and ROS accumulation in plants treated with UV-B. **(A)** Oxidative damage was evaluated by measuring ion leakage in 10 day-old seedlings irradiated with UV-B (continued lines) or filtered UV-B as a control condition (segmented lines) at different time points during treatment and recovery periods. Data show mean values ± standard deviation from 3 replicates. Asterisks represent statistically significant differences respect to wild type plants (WT) (two-way ANOVA/Fisher’s LSD test, p<0.01). The experiment was repeated three times with similar results. **(B)** ROS levels were detected using DAB staining in fifteen-day-old seedlings treated with UV-B radiation for 5 h. **(C)** The quantification of DAB staining was performed using the ImageJ software, calculating the percentage of stained area respect to the total rosette area for each seedling. Graph shows the mean value ± standard error from 10-16 wild type (WT), *tga256*, *tga256*/TGA2 #1 and *tga256*/TGA2 #2 seedlings irradiated with UV-B light (gray bars) or under control conditions (C, black bars). The experiment was repeated three times with similar results. Different letters above bars indicate significant differences (two-way ANOVA/Fisher’s LSD test p<0.01). **(D)** ROS produced were quantified using 2’,7’-dichlorodihydrofluorescein diacetate (H2DCF-DA) in plants irradiated with UV-B (gray bars) or under untreated control conditions (black bars) for 24 hours. Box plots show the distribution of eight to nine measurements, with the central line indicating the mean value and boxes the 5-95 percentile. Different letters above bars indicate significant differences between the samples (two-way ANOVA/Fisher’s LSD test p<0.05).

Taken together, these results indicate that expression of TGA class II genes, particularly TGA2, is essential to restrict ROS accumulation and protect tissues from oxidative damage after stress by UV-B treatment.

To further analyze the role of TGA2 in ROS containment after photooxidative stress, we performed treatments with Methyl Viologen (MeV, 1,1’-dimethyl-4,4’-bipyridylium chloride) in the presence of light. Under these conditions, MeV triggers photooxidative stress characterized by increased production of superoxide in the photosystem I (PSI) complex (Babbs et al., 1989; Fujii et al., 1990). We germinated seeds from wild type, *tga256* and *tga256*/TGA2 lines in 0.5X MS medium supplemented with 0.1 µM MeV and after 15 days we recorded the % of survival (% of germinated seeds that produce green seedlings). In this assay, MeV produced a strong reduction in seedling size in all lines, compared to control seedlings germinated in 0.5X MS medium (Fig. S5A). A comparison among lines indicates that mutation of class II TGAs (*tga256* mutant line) produced a significant reduction in the % of survival compared to wild type, an effect that was reversed in the *tga256*/TGA2 complemented lines (Fig. S5A-B). The higher susceptibility of the *tga256* mutant to germinate in MeV correlates with a lower capacity to restrict ROS production in MeV-treated seedlings (Fig. S5C). In fact, treatment of leaves from 15 days old seedlings with a drop of a MeV solution (15 and 30 µM) produced the localized accumulation of ROS in the treated tissue, which is less contained in the *tga256* mutant than in the wild type and complemented lines (Fig. S5C).

Together, these results support a key role of TGA class II factors in the antioxidative response triggered both by UV-B and by photooxidative stress conditions, particularly in activating a genetic response able to restrict ROS bursts and oxidative damage in stressed tissues.

### The expression of the GSTU7 gene complements the UV-B-sensitive phenotype of tga256 mutant plants

*GST* gene function, particularly the *GSTU* subfamily, is overrepresented in the group of genes positively controlled by TGA class II transcription factors in response to UV-B stress (Fig. 2). In order to evaluate the involvement of these *GSTU* genes in the tolerance to UV-B stress controlled by TGA class II genes (Fig. 4 and 5), we selected *GSTU7* as a representative UV-B-induced and TGA2-controlled gene (Fig. 3 and 4). We generated transgenic plants that constitutively express *GSTU7* in the *tga256* mutant background. We selected two *tga256*/GSTU7 lines (#1 and #2), in which we detected the transgenic GSTU7 protein by immunoblot using anti-V5 antibodies (Fig. 7A), for further analyses.

**Fig. 7.**
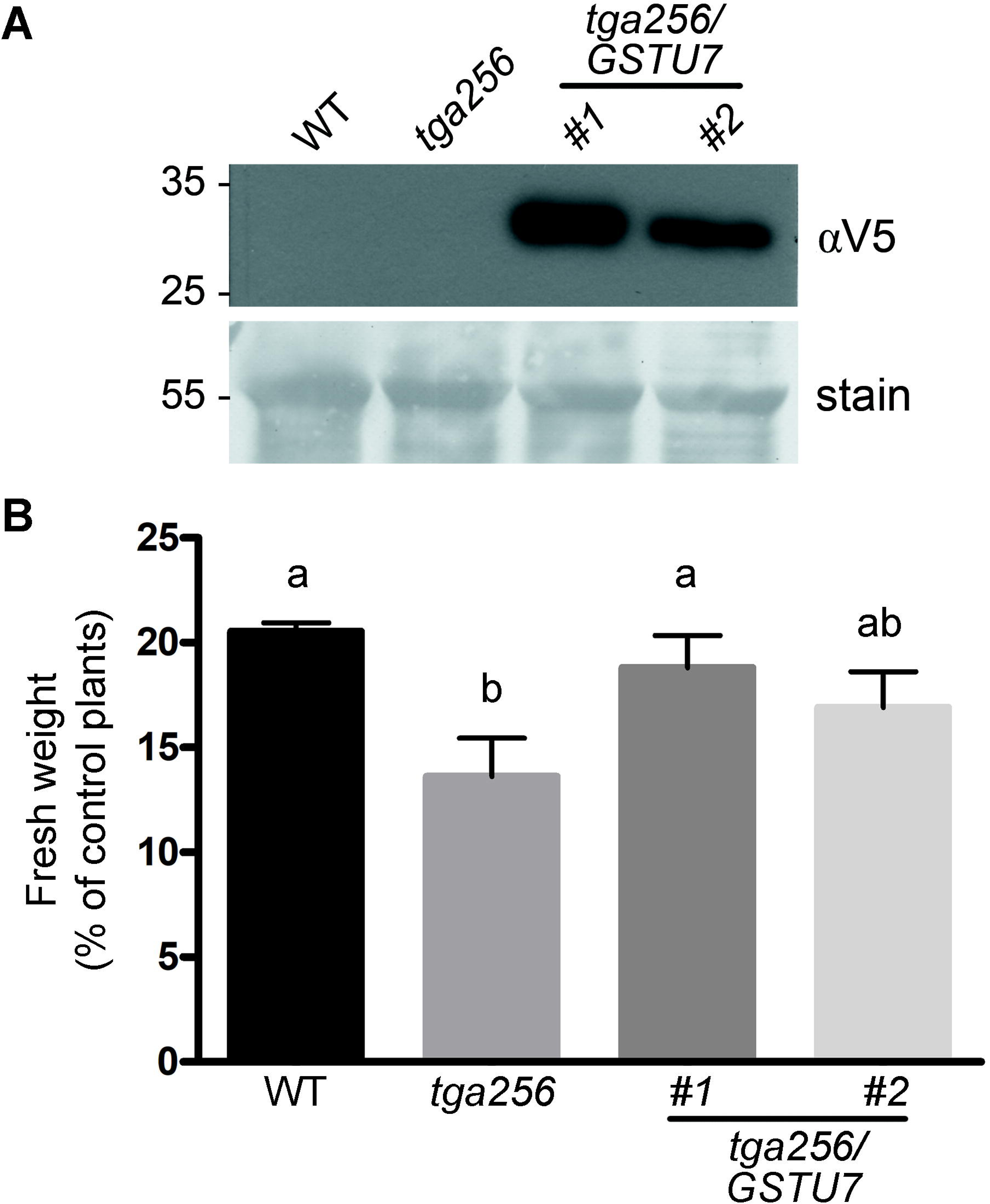
The expression of the *GSTU7* gene complements the UV-B-sensitive phenotype of *tga256* mutant plants. **(A)** Immunoblot for detection of the GSTU7-V5 protein in two *tga256/pUBQ:GSTU7-V5* complemented lines (*tga256/GSTU7* lines #1 and #2), and in the wild type (WT) and *tga256* plants as negative controls, using anti-V5 antibody (αV5). Coomassie staining (stain) indicates equivalent protein loading. **(B)** Fifteen day-old seedlings of the indicated genotypes were treated with UV-B radiation for 24 h and then recovered for 72 h under normal growth conditions. Fresh weight measurements were obtained at the end of the recovery period. Fresh weight of rosette tissue from UV-B treated plants was expressed as percentage of fresh weight of rosettes from control plants. Bars represent the mean ± standard deviation of 3 independent experiments (20 seedlings per genotype for each experiment). Statistical analysis was performed using ANOVA/Tukey’s test. Different letters denote statistically significant differences at p<0.05.

Consistent with the previous results (Fig. 1), we found a significant decrease in the fresh weight of *tga256* mutant plants compared to wild type plants after UV-B treatment, indicating increased susceptibility to this type of stress (Fig. 7B). Nevertheless, this phenotype was rescued when the mutant plants constitutively expressed the GSTU7 protein (Fig. 7B). This result indicates that GSTU enzymes, particularly GSTU7, are involved in the tolerance response to UV-B mediated by TGA2.

## Discussion

The evidence shown in this work supports an important role of TGA class II transcription factors - particularly TGA2 - as critical regulators of genes controlling ROS levels in the tolerance response to UV-B stress in Arabidopsis. First, plants of the *tga256* mutant genotype show a reduced capacity to scavenge ROS and restrict oxidative damage in response to UV-B and MeV-induced photooxidative stress. These phenotypes were complemented with a TGA2 transgene (*tga256*/TGA2 plants). Second, RNAseq followed by clustering analyses indicate that TGA2/5/6 factors positively control the UV-B-induced expression of a group of genes related to the antioxidative function. This group shows over-representation of GO terms associated to molecular functions of oxidoreductase and glutathione transferase activities, and includes several members of the *GST* gene family coding for peroxide-scavenging enzymes, particularly of the Tau subfamily (*GSTU* genes). Accordingly, the abolished or reduced induction of *GSTU* genes in the *tga256* mutant correlates with an impaired increase in GPX activity triggered by UV-B treatment. Third, the role of TGA2 as transcriptional activator of *GSTU* genes in response to UV-B was confirmed for *GSTU7*, *GSTU8*, and *GSTU25* genes through RT-qPCR and ChIP analyses in *tga256*/TGA2 plants. Fourth, the role for *GSTU* genes, particularly *GSTU7*, in the tolerance response to UV-B stress mediated by TGA2, was confirmed by the expression of GSTU7 in the *tga256* mutant plants (*tga256*/*GSTU7* plants). Together, this evidence supports a key role for TGA2 as activator of the antioxidant response that restricts ROS accumulation and oxidative damage during UV-B stress, through a mechanism that involves transcriptional activation of a set of genes coding for ROS-scavenging proteins such as GSTUs.

### GST genes in ROS modulation

The RNAseq and clustering analysis shown here gave a genetic support to the susceptibility phenotype of *tga256* mutant plants. Interestingly, many genes coding for peroxide-scavenging enzymes such as GSTUs, were detected in the clusters of UV-B-induced genes that are positively regulated by TGA2/5/6 (Fig. 2 and 3).

GSTUs represent the most numerous and one of the plant-specific subclasses of GSTs in Arabidopsis (Fig. 3A; (Moons, 2005). The idea that GSTs play a role in defense responses has been supported by evidence indicating: 1) their transcriptional induction in response to several biotic and abiotic stress conditions and 2) alterations in tolerance to stress conditions in plants with altered expression levels of some *GSTs*, using genetic tools (Nianiou-Obeidat et al., 2017; Kumar and Trivedi, 2018). Particularly, the overexpression of *GSTUs* has been proved to improve tolerance to oxidative stress produced by H_2_O_2_ (Sharma et al., 2014) and also by MeV (Yu et al., 2003; Xu et al., 2016). Previous reports showed that GST proteins possess glutathione-peroxidase (GPX) activity (Bartling et al., 1993; Cummins et al., 1999; Dixon et al., 2009), and that an increase in GPX activity by over-expression of *GSTs* genes produces a decrease in ROS accumulation (Cummins et al., 1999; Roxas et al., 2000). Accordingly, we observed that *tga256* mutant plants irradiated with UV-B show a reduction in levels of GPX activity and H_2_O_2_ accumulation, compared to irradiated wild type plants. The expression of TGA2 in the *tga256* background restores both phenotypes (Fig. 5 and 6).

### Involvement of TGA class II transcription factors in the stress defense response

TGA class II factors have been previously described as essential for development of the SAR against biotrophic pathogens (Zhang et al., 1999), due to their role as negative and positive transcriptional regulators of defense genes, such as *PR-1*, controlled by the SA- and NPR1-dependent pathway (Johnson et al., 2003; Zhang et al., 2003; Kesarwani et al., 2007; Pape et al., 2010). Also, our group and others have shown that TGA2/5/6 - and to a lesser extent TGA3 - are essential for the expression of *GRXC9*, a gene early induced by the SA-dependent and NPR1-independent pathway (Ndamukong et al., 2007; Herrera-Vasquez et al., 2015). The TGA2/5/6 factors positively regulates a high % of genes induced by oxylipins such as A1-phytoprostane (PPA1) and oxo-phytodienoic acid (OPDA) (Mueller et al., 2008). Furthermore, TGA2/5/6 factors have been proposed as a node of crosstalk between the SA- and the JA/ET-mediated pathways (Zander et al., 2012; Zander et al., 2014; Herrera-Vasquez et al., 2015), due to their role in controlling both the induction by JA/Et and the repression by SA of the *ORA59* gene, which codes for a master regulator of the JA/ET-mediated pathway (Zander et al., 2014). Accordingly, *tga256* mutant plants, in addition to a SAR-deficient phenotype, display increased susceptibility to the necrotrophic pathogen *B. cinerea* (Zander et al., 2014). Furthermore, the *tga256* mutant plants are more susceptible than wild type to germinate in the presence of 2,6-dichloroisonicotinic acid (INA), a SA-functional analog, and of 2,4,6-triiodobenzoic acid (TIBA), a chemical that blocks auxin transport (Fode et al., 2008).

In this context, the evidence shown here expands our knowledge about the role of TGA class II factors in the stress defense response, placing them as key positive regulators of the antioxidative defense response triggered by UV-B light exposure and by photooxidative stress induced by MeV. Accordingly, a clear phenotype of deficiency in the mechanisms that prevent ROS accumulation and oxidative damage was detected in *tga256* mutant plants subjected to both stress conditions.

In terms of mechanism, we show that TGA2 directly regulates the UV-B-induced expression of the *GSTU7*, *GSTU8* and *GSTU25* genes, as well as the basal expression of *GSTU7* and *GSTU8* (Fig. 4). This regulation seems to be mediated by the recruitment of TGA2 to *GSTU* promoters, as supported by the correlation between the basal and UV-B-induced transcript levels of these genes (Fig. 3C and 4A), and the TGA2 binding to their promoters (Fig. 4C). This was also the case for *PR-1* induction triggered by SA, where *PR-1* gene regulation is also mediated by the recruitment of TGA2 (and TGA3) to its promoter (Johnson et al. 2003). In contrast, in the case of the *GRXC9* and *ORA59* genes, TGA2 is constitutively bound to the respective promoters under basal or induced conditions (Zander et al., 2014; Herrera-Vasquez et al., 2015). These cases show the versatility of TGA2 as a transcriptional regulator. This versatility could be given by different co-regulators that act as TGA2-interacting partners at the promoter of different genes and during different stages of the defense response, such as NPR1, SCL14 and GRXC9 proteins (Johnson et al., 2003; Ndamukong et al., 2007; Fode et al., 2008; Herrera-Vasquez et al., 2015). An analysis of these interactions, in the context of the redox changes occurring during the defense response, lead us to propose TGA2/5/6 as a potential platform for redox regulation of defense gene expression (Ndamukong et al., 2007; Herrera-Vasquez et al., 2015).

Supporting the participation of TGA2 in redox control mediated by ROS, among the studies focused on describing cis-elements in gene promoters associated with oxidative stress

(Garretón et al., 2002; Petrov et al., 2012; Wang et al., 2013), our previous report shows that the TGA-binding motif *as-1* acts as an oxidative stress-responsive element (Garretón et al., 2002).

The evidence shown here provides further support to the idea that TGA class II transcription factors represent a redox regulatory node in stress responses. Then, TGA2/5/6 seem to be not only responsive to redox signals for controlling activation/repression of different groups of genes (Herrera-Vasquez et al., 2015), but they also impact on the cellular redox state by controlling the expression of genes responsible for restraining ROS increase, and therefore oxidative damage in response to stress.

## List of author contributions

L.H. and A.H-V. designed the research; A.H-V., A.F., J.M.U., L.L. and A.S. designed and performed the experiments and analyzed results; E.V., T.M. and R.G. performed and supervised the analysis of RNAseq data; P.S. supervised some experiments; L.H. and A.H-V. wrote the manuscript with input from other authors.

## Supplementary data

**Figure S1.** Expression of TGA2 complements the *tga256* mutant phenotype.

**Figure S2.** TGA class II are redundant in the response to UV-B

**Figure S3.** *CHS* and *PR-1* expression levels in wild type and *tga256* mutant plants in response to UV-B treatment.

**Figure S4.** Global expression analysis of wild type and *tga256* triple mutant plants in response to UV-B treatment.

**Figure S5:** The TGA2 factor is essential for tolerance and ROS control in response to photo oxidative stress.

**Table S1.** UV-B responsive genes regulated by TGA2/5/6 factors. List of genes differentially regulated by the interaction between treatment and genotype.

**Table S2.** Primers used for cloning, ChIP and RT-qPCR assays.

## Acknowledgements

The authors thank Xin Li (Department of Botany, University of British Columbia, Canada) for providing the *tga6-1*, *tga2-1 tga5-1* and *tga2-1 tga5-1 tga6-1* mutant lines.

This work was supported by the National Commission for Science and Technology CONICYT (FONDECYT grants N° 1141202 to L.H. and Nº 1141029 to P.S.) and the Millennium Science Initiative (Nucleus for Plant Synthetic and Systems Biology, grant NC130030 and Institute for Integrative Biology (iBio)). T.C.M., J.M.U. and A.S. were supported by a PhD fellowship from CONICYT.

## Notes

### Competing Interest Statement

The authors have declared no competing interest.

