## Supplementary material for "Transcription factor TGA2 is essential for UV-B stress tolerance controlling oxidative stress in Arabidopsis": Fig. S

### Supplementary data

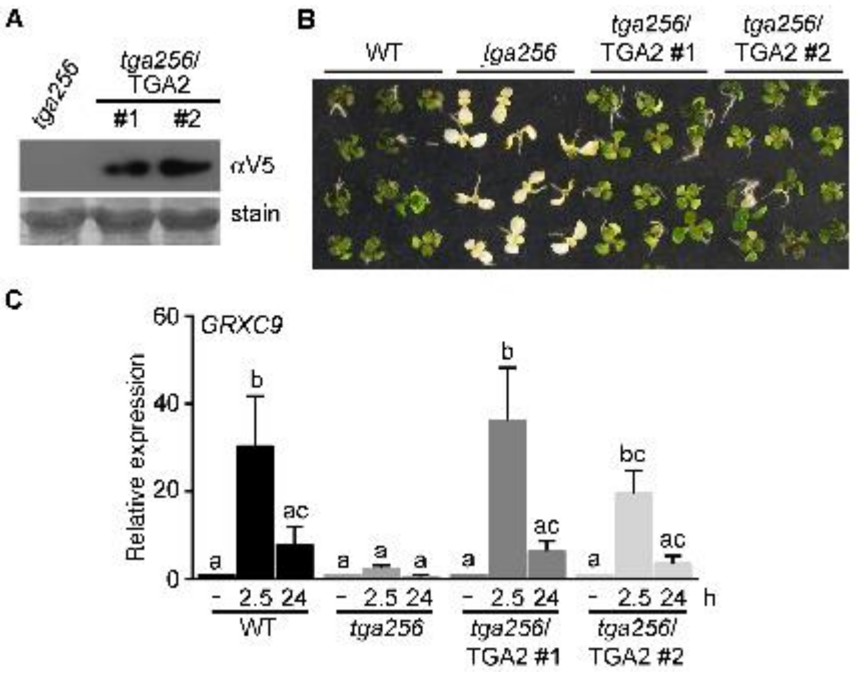


**Fig. S1. Expression of TGA2 complements the *tga256* mutant phenotype. (A)** Immunoblot for detection of the TGA2-V5 protein in two *tga256/pUBQ:TGA2-V5* complemented lines (*tga256/*TGA2 lines #1 and #2), and in the *tga2-1 tga5-1 tga6-1* mutant (*tga256*) as a negative control, using anti-V5 antibody (αV5). Coomassie staining (stain) indicates equivalent protein loading. **(B)** Assay of tolerance to germinate in SA. Seeds from wild type (WT), *tga256* mutant and the two *tga256*/TGA2-complemented lines were plated on 0.5X MS medium supplemented with 0.2 mM SA. The figure shows survival after 15 days. **(C)** Expression analysis of the *GRXC9* gene evaluated by RT-qPCR in 15-day-old seedlings (WT, *tga256*, and the two *tga256*/TGA2 complemented lines), under basal conditions (-) and after treatment with SA 0.5 mM for 2.5 and 24 h. *GRXC9* relative expression was calculated by normalizing *GRXC9* transcript levels to transcript levels of the housekeeping gene *YLS8* (AT5G08290), and to the WT basal condition. Bars represent the mean ± standard error values from at least three biological replicates. Statistical analysis was performed using ANOVA/Fisher's LSD test. Different letters denote statistically significant differences at $p<0.05$.

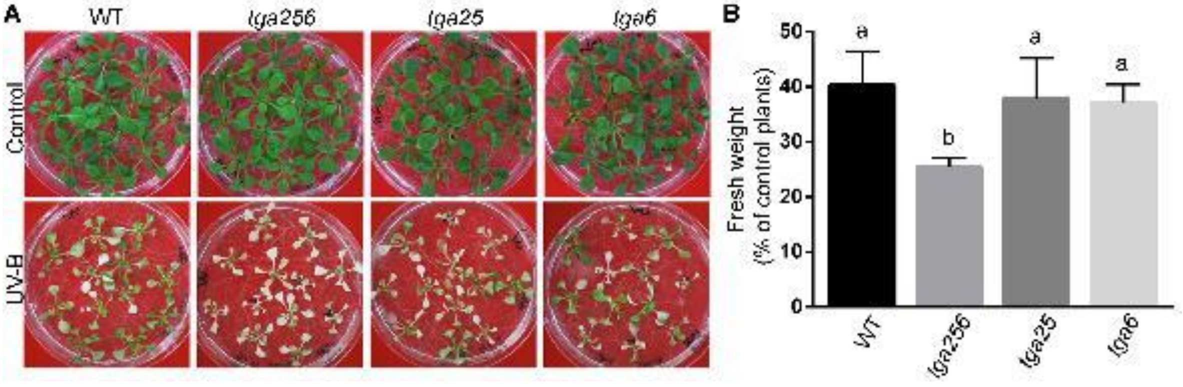


**Fig. S2. TGA class II are redundant in the response to UV-B.** Fifteen day-old seedlings of WT, *tga256*, *tga2-1 tga5-1* (*tga25*) and *tga6-1 (tga6)* mutant plants were treated with UV-B radiation for 24 h and then they recovered for 72 h in a growth chamber. Control treatments were performed under the same conditions with a UV-B filter. Pictures **(A)** and fresh weight measurements **(B)** were obtained at the end of the recovery period. Fresh weight of rosette tissue from UV-B treated plants was expressed as percentage of fresh weight of rosettes from control plants. Bars represent the mean ± standard deviation of at least 3 independent experiments (20 seedlings per genotype for each experiment). Statistical analysis was performed using ANOVA/Fisher's LSD test. Different letters denote statistically significant differences at $p<0.05$.

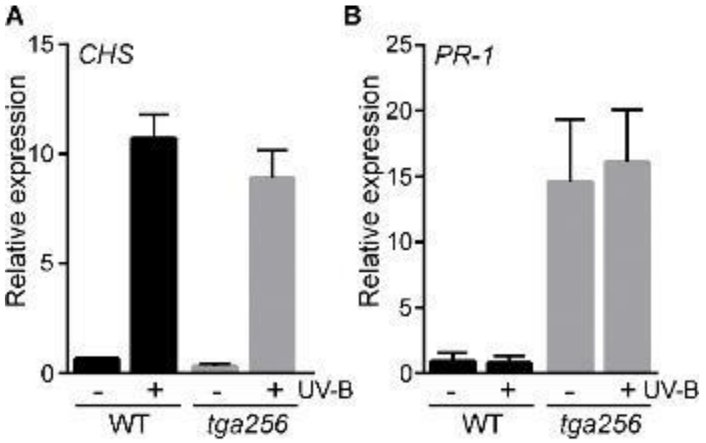


**Fig. S3. *CHS* and *PR-1* expression levels in wild type and *tga256* mutant plants in response to UV-B treatment.** Expression levels of CHS (A) and PR-1 (B) genes were measured by RT-qPCR in 15-day-old seedlings from wild type (WT, black bars) and tga256 mutant plants (gray bars) exposed to UV-B light during 5 hours (+). As a control, we used seedlings covered with a cellulose acetate polyester filter (-). Relative expression was calculated by normalizing gene transcript levels to transcript levels of the housekeeping gene YLS8 (AT5G08290). Error bars represent the mean ± SD from 3 replicates.

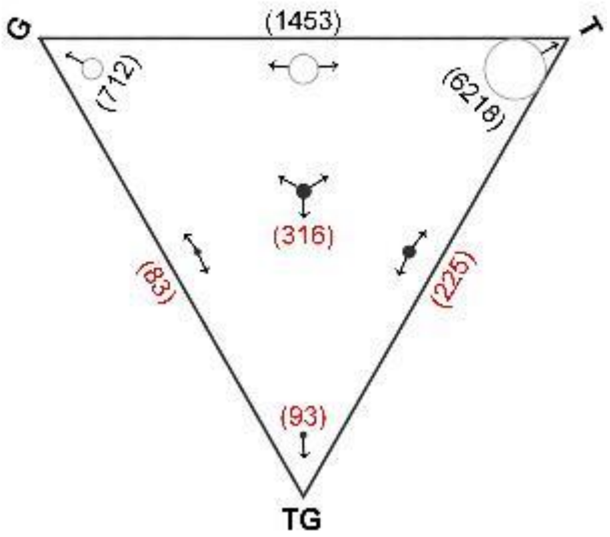


**Fig. S4. Global expression analysis of wild type and *tga256* triple mutant plants in response to UV-B treatment.** RNAseq data analyzed using two-way ANOVA ($p<0.01$) is represented using the Sungear tool and show the number of genes that are differentially expressed (Poultney et al., 2007). The triangle shows the factors at the vertices: UV-B treatment (T), genotype (G) and the interaction between UV-B treatment and genotype (TG). The circles inside the triangle represent the number of genes (in parentheses) controlled by the different factors, as indicated by the arrows around the circles. The size of each circle is proportional to the number of genes associated with that circle. White circles represent the genes differentially regulated by genotype or treatment alone; black circles and red numbers represent the 717 genes differentially regulated by the interaction between treatment and genotype (Table S1).

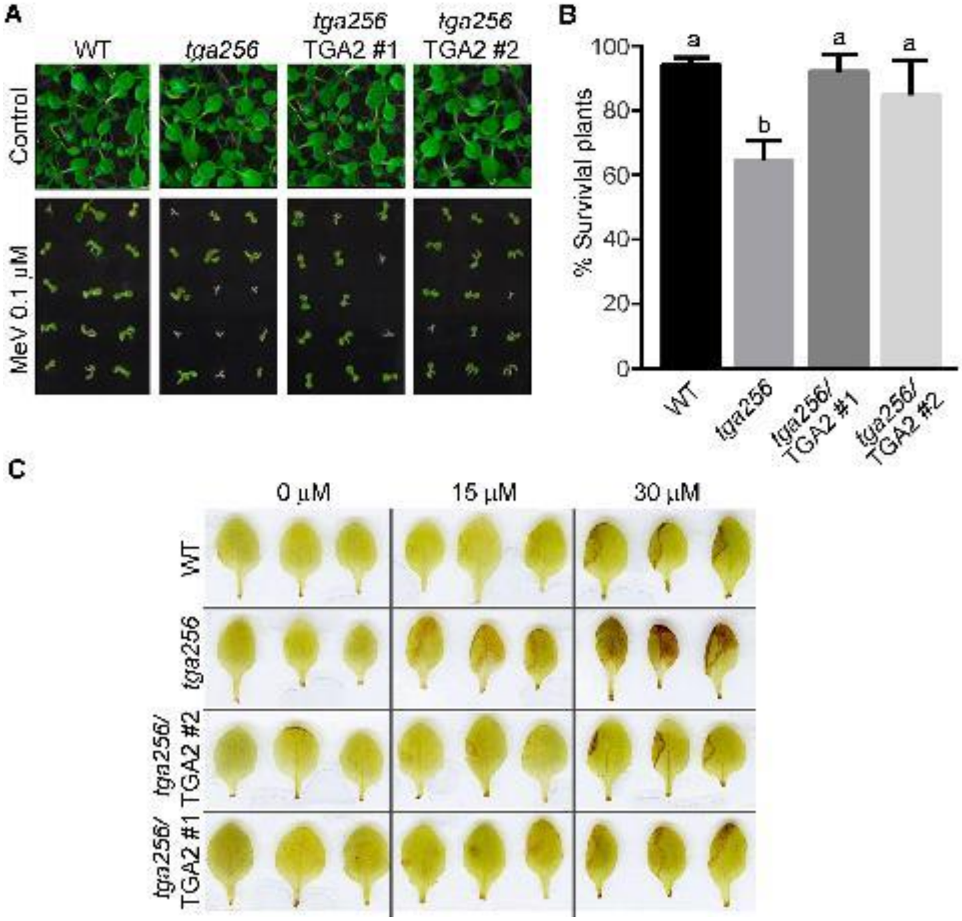


**Fig. S5. The TGA2 factor is essential for tolerance and ROS control in response to photo oxidative stress. (A-B)** Seeds from wild type plants (WT), the *tga256* mutant and the two *tga256*/TGA2 complemented lines were germinated in ½ MS medium alone (control) or in ½ MS medium supplemented with 0.1 µM MeV. After 15 days, pictures of seedlings were taken **(A)** and the % plant survival (% of green seedlings respect to the total germinated seeds) was recorded **(B).** Data show mean values ± standard deviation of 4 independent experiments (104 seeds each). Letters represent statistical differences between genotypes (one-way ANOVA/Fisher's LSD test, $p<0.01$). **(C)** Leaves from 15 day-old WT, *tga256* mutant and two *tga256*/TGA2 complemented lines seedlings were treated with a 2 µL drop of MeV (0, 15 and 30 µM) and incubated under constant light for 24 hours. ROS accumulation in the treated leaves was detected by DAB staining (1 mg/ml during 4 hours). The figure shows representative images of leaves from different treated plants (n=20). This experiment was repeated three times with similar results.

**Table S1.** UV-B responsive genes regulated by TGA2/5/6 factors. List of genes differentially regulated by the interaction between treatment and genotype.

**Table S2.** Primers used for cloning, ChIP and RT-qPCR assays.
