## Supplementary material for "Transcription factor TGA2 is essential for UV-B stress tolerance controlling oxidative stress in Arabidopsis": Table S1

**Supplementary Table S1. UV-B responsive genes regulated by TGA2/5/6 factors.** List of genes differentially regulated by the interaction between treatment and genotype.

| | | | Normalized expression | | | |
|---|---|---|---|---|---|---|
| | | | *Wild type* | | *tga256* | |
| Locus | Gene model Description | Gene Symbol | Control | UV-B | Control | UV-B |
| | ***Cluster 1*** | | | | | |
| At2g46950 | CYP709B2, cytochrome P450, family 709, subfamily B, polypeptide 2 | *CYP79B2* | 9,13 ± 0,93 | 18,89 ± 1,29 | 1,79 ± 1,29 | 0,44 ± 0,72 |
| At2g17830 | F-box and associated interaction domains-containing protein | | 41,53 ± 2,16 | 53,57 ± 2,09 | 38,96 ± 2,09 | 32,74 ± 5,21 |
| At1g33160 | Pseudogene, similar to actin | | 5,55E-17 ± 7,55E-33 | 6,48 ± 7,55E-33 | 5,55E-17 ± 7,55E-33 | 0,8 ± 1,29 |
| At4g16740 | ATTPS03, TPS03, terpene synthase 03 | *TPS3* | 1,99 ± 2,49 | 14,38 ± 1,25 | 0,73 ± 1,25 | 0,42 ± 0,68 |
| At1g22800 | S-adenosyl-L-methionine-dependent methyltransferases superfamily protein | | 236,22 ± 14,03 | 325,11 ± 5,92 | 222,27 ± 5,92 | 213,85 ± 35,64 |
| At4g33160 | F-box family protein | | 87,37 ± 14,05 | 98,6 ± 9,29 | 76,38 ± 9,29 | 49,01 ± 8,56 |
| At5g17030 | UGT78D3, UDP-glucosyl transferase 78D3 | *UGT78D3* | 5,55E-17 ± 7,55E-33 | 140,69 ± 0,63 | 0,36 ± 0,63 | 29,95 ± 15,67 |
| At1g64940 | CYP89A6, cytochrome P450, family 87, subfamily A, polypeptide 6 | *CYP89A6* | 4,05 ± 3,28 | 145,03 ± 0,76 | 1,8 ± 0,76 | 9,48 ± 3,08 |
| At1g69860 | Major facilitator superfamily protein | | 5,55E-17 ± 7,55E-33 | 2,36 ± 7,55E-33 | 5,55E-17 ± 7,55E-33 | 0,4 ± 0,65 |
| At1g79710 | Major facilitator superfamily protein | | 461,51 ± 56,02 | 473,68 ± 22,16 | 267,45 ± 22,16 | 171,25 ± 14,88 |
| At5g49130 | MATE efflux family protein | | 2,37 ± 2,01 | 35,91 ± 5,22 | 5,77 ± 5,22 | 5,7 ± 4,44 |
| At5g16980 | Zinc-binding dehydrogenase family protein | | 22,78 ± 7,22 | 72,68 ± 0,62 | 0,36 ± 0,62 | 2,64 ± 1,22 |
| At4g34135 | UGT73B2, UDP-glucosyltransferase 73B2 | *UGT73B2* | 375,44 ± 69,37 | 1533,89 ± 3 | 54,84 ± 3 | 156,93 ± 30,99 |
| At5g22140 | FAD/NAD(P)-binding oxidoreductase family protein | | 142,64 ± 1,14 | 461,87 ± 18,96 | 111,4 ± 18,96 | 73,78 ± 25,05 |
| At1g29195 | Unknown protein | | 24,95 ± 4,59 | 43,23 ± 0,54 | 10,97 ± 0,54 | 10,91 ± 1,58 |
| At3g08750 | F-box and associated interaction domains-containing protein | | 5,55E-17 ± 7,55E-33 | 4,43 ± 7,55E-33 | 5,55E-17 ± 7,55E-33 | 0,4 ± 0,65 |
| At1g33760 | Integrase-type DNA-binding superfamily protein | | 2,8 ± 1,64 | 9,09 ± 1,17 | 1,38 ± 1,17 | 5,55E-17 ± 7,55E-33 |
| At2g33200 | F-box family protein | | 5,55E-17 ± 7,55E-33 | 4,7 ± 1,24 | 0,72 ± 1,24 | 5,55E-17 ± 7,55E-33 |
| At1g11230 | Protein of unknown function (DUF761) | | 5,55E-17 ± 7,55E-33 | 17,27 ± 1,24 | 0,72 ± 1,24 | 2,13 ± 1,79 |
| At5g18890 | Inosine-uridine preferring nucleoside hydrolase family protein | *NUCLEOSIDE HYDROLASE 4 (NSH4)* | 5,55E-17 ± 7,55E-33 | 1,02 ± 7,55E-33 | 5,55E-17 ± 7,55E-33 | 5,55E-17 ± 7,55E-33 |
| At3g09640 | APX1B, APX2, ascorbate peroxidase 2 | *APX2* | 5,55E-17 ± 7,55E-33 | 12,05 ± 7,55E-33 | 5,55E-17 ± 7,55E-33 | 3,5 ± 0,72 |
| At2g46130 | ATWRKY43, WRKY43, WRKY DNA-binding protein 43 | *WRKY43* | 3,73 ± 0,6 | 16,27 ± 0,53 | 1,76 ± 0,53 | 4,41 ± 2,98 |
| At2g15490 | UGT73B4, UDP-glycosyltransferase 73B4 | *UGT73B4* | 70,83 ± 13,99 | 2412,29 ± 7,95 | 20,45 ± 7,95 | 415,79 ± 47,05 |
| At5g45870 | PYL12, RCAR6, PYR1-like 12 | *PYL12* | 1,19 ± 1,09 | 15,85 ± 7,55E-33 | 5,55E-17 ± 7,55E-33 | 1,76 ± 1,45 |

**Supplementary Table S1. UV-B responsive genes regulated by TGA2/5/6 factors.** List of genes differentially regulated by the interaction between treatment and genotype.

| Locus | Gene model Description | Gene Symbol | Normalized expression | | | |
|---|---|---|---|---|---|---|
| | | | *Wild type* | | *tga256* | |
| | | | Control | UV-B | Control | UV-B |
| | ***Cluster 1*** | | | | | |
| At3g13784 | AtcwINV5, CWINV5, cell wall invertase 5 | *CELL WALL INVERTASE 5 (CWINV5)* | 8,03 ± 2,87 | 95,96 ± 2,16 | 2,1 ± 2,16 | 9,57 ± 8,19 |
| At1g04380 | 2-oxoglutarate (2OG) and Fe(II)-dependent oxygenase superfamily protein | | 2,53 ± 1,51 | 12,27 ± 1,17 | 1,38 ± 1,17 | 3,53 ± 1,56 |
| At5g40610 | NAD-dependent glycerol-3-phosphate dehydrogenase family protein | *(GPDHP)* | 287,5 ± 28,52 | 442,31 ± 8,04 | 277,39 ± 8,04 | 310,96 ± 19,03 |
| At5g40348 | other RNA | | 5,55E-17 ± 7,55E-33 | 2,35 ± 7,55E-33 | 5,55E-17 ± 7,55E-33 | 5,55E-17 ± 7,55E-33 |
| At2g29490 | ATGSTU1, GST19, GSTU1, glutathione S-transferase TAU 1 | *GSTU1* | 124,51 ± 32,3 | 610,5 ± 7,11 | 31,57 ± 7,11 | 60,77 ± 13,89 |
| At2g33710 | Integrase-type DNA-binding superfamily protein | | 26,35 ± 7,43 | 289,12 ± 5,31 | 13,31 ± 5,31 | 63,68 ± 15,72 |
| At4g26280 | P-loop containing nucleoside triphosphate hydrolases superfamily protein | | 5,55E-17 ± 7,55E-33 | 1,68 ± 7,55E-33 | 5,55E-17 ± 7,55E-33 | 5,55E-17 ± 7,55E-33 |
| At4g02940 | oxidoreductase, 2OG-Fe(II) oxygenase family protein | | 809,97 ± 43,64 | 1413,2 ± 32,6 | 641,16 ± 32,6 | 588,77 ± 120,42 |
| At4g20840 | FAD-binding Berberine family protein | | 222,4 ± 30,57 | 263,35 ± 12,51 | 164,8 ± 12,51 | 127,5 ± 5,87 |
| At4g15755 | Calcium-dependent lipid-binding (CaLB domain) family protein | | 5,55E-17 ± 7,55E-33 | 2,69 ± 7,55E-33 | 5,55E-17 ± 7,55E-33 | 5,55E-17 ± 7,55E-33 |
| At2g29452 | | | 5,55E-17 ± 7,55E-33 | 3,04 ± 0,62 | 0,36 ± 0,62 | 0,42 ± 0,68 |
| At5g06960 | OBF5, TGA5, OCS-element binding factor 5 | *OBF5* | 177,86 ± 9,3 | 236,98 ± 0,94 | 1,02 ± 0,94 | 0,83 ± 1,35 |
| At1g79410 | 5-Oct, AtOCT5, organic cation/carnitine transporter5 | *5OCT* | 159,82 ± 35,2 | 496,91 ± 5,22 | 24,11 ± 5,22 | 42,72 ± 14,61 |
| At3g49520 | F-box and associated interaction domains-containing protein | | 5,55E-17 ± 7,55E-33 | 2,02 ± 7,55E-33 | 5,55E-17 ± 7,55E-33 | 5,55E-17 ± 7,55E-33 |
| At5g49138 | other RNA | | 2,37 ± 2,01 | 33,14 ± 4,71 | 4,72 ± 4,71 | 4,84 ± 4,75 |
| At4g34138 | UGT73B1, UDP-glucosyl transferase 73B1 | *UGT73B1* | 757,82 ± 55,62 | 1196,9 ± 20,1 | 59,69 ± 20,1 | 31,76 ± 11,33 |
| At2g38010 | Neutral/alkaline non-lysosomal ceramidase | | 180,17 ± 22,13 | 358,03 ± 9,42 | 199,47 ± 9,42 | 181,72 ± 34,08 |
| At1g26233 | SNOR95, SNOR95; snoRNA | *SMALL NUCLEOLAR RNA95 (SNOR95)* | 0,24 ± 0,41 | 5,03 ± 0,63 | 0,36 ± 0,63 | 1,33 ± 0,08 |
| At2g29480 | ATGSTU2, GST20, GSTU2, glutathione S-transferase tau 2 | *GSTU2* | 16,5 ± 5,13 | 182,09 ± 3,71 | 2,17 ± 3,71 | 21,9 ± 8,5 |
| At1g64340 | Unknown protein | | 5,55E-17 ± 7,55E-33 | 66,38 ± 7,55E-33 | 5,55E-17 ± 7,55E-33 | 6,9 ± 3,85 |
| At3g04000 | NAD(P)-binding Rossmann-fold superfamily protein | | 51,08 ± 16,77 | 253,55 ± 9,23 | 49,18 ± 9,23 | 98,46 ± 38,82 |

**Supplementary Table S1. UV-B responsive genes regulated by TGA2/5/6 factors.** List of genes differentially regulated by the interaction between treatment and genotype.

| | | | Normalized expression | | | |
|---|---|---|---|---|---|---|
| | | | *Wild type* | | *tga256* | |
| Locus | Gene model Description | Gene Symbol | Control | UV-B | Control | UV-B |
| ***Cluster 1*** | | | | | | |
| At5g53760 | ATMLO11, MLO11, Seven transmembrane MLO family protein | *(MLO11)* | 136,22 ± 19,61 | 285,45 ± 23,11 | 133,97 ± 23,11 | 145,35 ± 16,05 |
| At2g29380 | HAI3, highly ABA-induced PP2C gene 3 | *HAI3* | 1,25 ± 1,72 | 16,73 ± 7,55E-33 | 5,55E-17 ± 7,55E-33 | 1,2 ± 1,94 |
| At1g62690 | unknown protein | | 1,2 ± 1,55 | 7,15 ± 0,75 | 2,49 ± 0,75 | 1,3 ± 1,26 |
| At2g39240 | RNA polymerase I specific transcription initiation factor RRN3 protein | | 3,26 ± 1,26 | 15,35 ± 1,17 | 5,32 ± 1,17 | 5,73 ± 2,76 |
| At5g41250 | Exostosin family protein | | 5,55E-17 ± 7,55E-33 | 1,72 ± 7,55E-33 | 5,55E-17 ± 7,55E-33 | 5,55E-17 ± 7,55E-33 |
| At2g35090 | Protein of unknown function (DUF1640) | | 5,55E-17 ± 7,55E-33 | 1,37 ± 7,55E-33 | 5,55E-17 ± 7,55E-33 | 5,55E-17 ± 7,55E-33 |
| At1g62975 | basic helix-loop-helix (bHLH) DNA-binding superfamily protein | | 43,56 ± 10,01 | 75,46 ± 0,82 | 24,04 ± 0,82 | 13,52 ± 1,05 |
| At5g04400 | anac077, NAC077, NAC domain containing protein 77 | *NAC77* | 5,55E-17 ± 7,55E-33 | 4,41 ± 0,62 | 0,36 ± 0,62 | 0,42 ± 0,68 |
| At3g25240 | Protein of unknown function (DUF506) | | 3,7 ± 3,96 | 136,51 ± 1,28 | 1,48 ± 1,28 | 46,23 ± 15,43 |
| At1g19210 | Integrase-type DNA-binding superfamily protein | | 0,53 ± 0,92 | 9,83 ± 1,2 | 2,16 ± 1,2 | 1,68 ± 1,34 |
| At1g78340 | ATGSTU22, GSTU22, glutathione S-transferase TAU 22 | *GSTU22* | 44,13 ± 10,81 | 280,45 ± 4,77 | 36,22 ± 4,77 | 35,36 ± 8,2 |
| At1g76690 | ATOPR2, OPR2, 12-oxophytodienoate reductase 2 | *OPR2* | 250,49 ± 22,65 | 624,97 ± 10,14 | 57,49 ± 10,14 | 89,39 ± 18,56 |
| At3g59140 | ATMRP14, MRP14, multidrug resistance-associated protein 14 | *ATP-BINDING CASSETTE C1 (ABCC1)* | 133,06 ± 64,35 | 726,96 ± 29,29 | 104,9 ± 29,29 | 150,41 ± 31,83 |
| At4g18050 | PGP9, P-glycoprotein 9 | *ATP-BINDING CASSETTE B9 (ABCB9)* | 123,93 ± 11 | 355 ± 11,51 | 83,53 ± 11,51 | 144,03 ± 24,16 |
| At1g17180 | ATGSTU25, GSTU25, glutathione S-transferase TAU 25 | *GSTU25* | 76,53 ± 27,71 | 3139,32 ± 8,16 | 20,48 ± 8,16 | 134,99 ± 56,66 |
| At1g22810 | Integrase-type DNA-binding superfamily protein | | 3,94 ± 2,59 | 93,66 ± 2,11 | 7,77 ± 2,11 | 19,68 ± 9,1 |
| At1g17170 | ATGSTU24, GST, GSTU24, glutathione S-transferase TAU 24 | *GSTU24* | 151,35 ± 4,76 | 2739,9 ± 6,33 | 42,05 ± 6,33 | 483,69 ± 123,46 |
| At3g19920 | Unknown protein | | 5,55E-17 ± 7,55E-33 | 22,91 ± 0,59 | 0,69 ± 0,59 | 7,38 ± 1,55 |
| At2g32910 | DCD (Development and Cell Death) domain protein | | 512,81 ± 21,28 | 724,72 ± 54,02 | 545,41 ± 54,02 | 591,14 ± 34,14 |
| At1g60750 | NAD(P)-linked oxidoreductase superfamily protein | | 4,86 ± 3,15 | 141,18 ± 1,28 | 1,48 ± 1,28 | 14,31 ± 2,6 |
| At4g13180 | NAD(P)-binding Rossmann-fold superfamily protein | | 348,02 ± 49,5 | 1680,41 ± 5,86 | 40,93 ± 5,86 | 150 ± 1,16 |

**Supplementary Table S1. UV-B responsive genes regulated by TGA2/5/6 factors.** List of genes differentially regulated by the interaction between treatment and genotype.

| Locus | Gene model Description | Gene Symbol | Normalized expression | | | |
|---|---|---|---|---|---|---|
| | | | *Wild type* | | *tga256* | |
| | | | Control | UV-B | Control | UV-B |
| | ***Cluster 1*** | | | | | |
| At5g07190 | ATS3, seed gene 3 | *SEED GENE 3 (ATS3)* | 2,43 ± 2,33 | 62,63 ± 0,54 | 0,32 ± 0,54 | 8,36 ± 6,8 |
| At1g22240 | APUM8, PUM8, pumilio 8 | *PUMILIO 8 (PUM8)* | 0,24 ± 0,41 | 15,08 ± 1,25 | 0,73 ± 1,25 | 2,61 ± 2,4 |
| At5g40605 | transposable element gene | | 5,55E-17 ± 7,55E-33 | 1,68 ± 7,55E-33 | 5,55E-17 ± 7,55E-33 | 5,55E-17 ± 7,55E-33 |
| At4g15760 | MO1, monooxygenase 1 | *MONOOXYGENASE 1 (MO1)* | 542,21 ± 69,74 | 2921,59 ± 3,52 | 89,29 ± 3,52 | 851,37 ± 66,71 |
| At4g25930 | Protein of unknown function (DUF295) | | 5,55E-17 ± 7,55E-33 | 9,42 ± 1,08 | 1,05 ± 1,08 | 2,1 ± 2,31 |
| At5g16920 | Fasciclin-like arabinogalactan family protein | | 2,31 ± 0,31 | 3,74 ± 7,55E-33 | 5,55E-17 ± 7,55E-33 | 5,55E-17 ± 7,55E-33 |
| At1g05562 | Potential natural antisense gene, locus overlaps with AT1G05560 | | 561,21 ± 38,07 | 1721,63 ± 73,15 | 342,83 ± 73,15 | 200,08 ± 27,02 |
| At3g50100 | SDN1, small RNA degrading nuclease 1 | *SMALL RNA DEGRADING NUCLEASE 1 (SDN1)* | 62,23 ± 5,05 | 69,55 ± 4,88 | 46,33 ± 4,88 | 31,49 ± 5,12 |
| At1g05560 | UGT1, UGT75B1, UDP-glucosyltransferase 75B1 | *UGT75B1* | 624,64 ± 41,47 | 1971,01 ± 79,08 | 365,84 ± 79,08 | 212,61 ± 30,1 |
| At2g40170 | ATEM6, EM6, GEA6, Stress induced protein | *LATE EMBRYOGENESIS ABUNDANT 6 (GEA6)* | 0,46 ± 0,4 | 8,75 ± 7,55E-33 | 5,55E-17 ± 7,55E-33 | 2,09 ± 1,74 |
| | ***Cluster 2*** | | | | | |
| At5g52100 | crr1, Dihydrodipicolinate reductase, bacterial/plant | *CHLORORESPIRATIO N REDUCTION 1 (crr1)* | 458,37 ± 15,74 | 303,9 ± 22,65 | 544,73 ± 22,65 | 254,78 ± 42,24 |
| At2g13360 | AGT, AGT1, SGAT, alanine:glyoxylate aminotransferase | *ALANINE:GLYOXY-LATE AMINO-TRANSFERASE* | 10500,75 ± 369,61 | 9152,6 ± 3560,91 | 19150,31 ± 3560,91 | 7408,92 ± 1016,83 |
| At3g14650 | CYP72A11, cytochrome P450, family 72, subfamily A, polypeptide 11 | *CYP72A11* | 1116,34 ± 62,01 | 1141,95 ± 89,74 | 1302,47 ± 89,74 | 803,32 ± 87,97 |
| At1g32060 | PRK, phosphoribulokinase | *PHOSPHORIBULOKIN ASE (PRK)* | 9959,61 ± 820,2 | 8744,09 ± 2835,56 | 15133,72 ± 2835,56 | 7140,73 ± 834,27 |
| At2g33750 | ATPUP2, PUP2, purine permease 2 | *PURINE PERMEASE 2 (PUP2)* | 11,66 ± 1,49 | 1,43 ± 0,86 | 23,62 ± 0,86 | 5,55E-17 ± 7,55E-33 |
| At3g02540 | RAD23-3, RAD23C, Rad23 UV excision repair protein family | *RADIATION SENSITIVE23C (RAD23C)* | 1110,82 ± 159,8 | 853,75 ± 64,29 | 1525,38 ± 64,29 | 718,56 ± 82,9 |
| At3g44020 | thylakoid lumenal P17.1 protein | | 120,21 ± 12,24 | 101,69 ± 30,26 | 164,17 ± 30,26 | 65,69 ± 14,29 |
| At1g80440 | Galactose oxidase/kelch repeat superfamily protein | | 1000,41 ± 173,91 | 263,56 ± 15,34 | 2008,76 ± 15,34 | 316,32 ± 4,09 |

**Supplementary Table S1. UV-B responsive genes regulated by TGA2/5/6 factors.** List of genes differentially regulated by the interaction between treatment and genotype.

| | | | Normalized expression | | | |
|---|---|---|---|---|---|---|
| | | | *Wild type* | | *tga256* | |
| Locus | Gene model Description | Gene Symbol | Control | UV-B | Control | UV-B |
| ***Cluster 2*** | | | | | | |
| At4g32570 | TIFY8, TIFY domain protein 8 | *TIFY DOMAIN PROTEIN 8 (TIFY8)* | 671,13 ± 53,98 | 336,56 ± 70,42 | 903,49 ± 70,42 | 340,43 ± 20,99 |
| At1g15260 | Unknown protein | | 303,45 ± 6,37 | 33,74 ± 24,8 | 309,19 ± 24,8 | 48,68 ± 2,8 |
| At1g80060 | Ubiquitin-like superfamily protein | | 34,63 ± 5,81 | 28,9 ± 3,7 | 54,3 ± 3,7 | 26,55 ± 2,78 |
| At3g14415 | Aldolase-type TIM barrel family protein | | 4220,67 ± 563,71 | 3774,71 ± 752,93 | 6492,02 ± 752,93 | 3253,01 ± 328,97 |
| At1g07970 | CONTAINS InterPro DOMAIN/s: Cytochrome B561-related | | 300,19 ± 22,25 | 241,97 ± 36,78 | 340,36 ± 36,78 | 189,13 ± 11,01 |
| At5g41280 | Receptor-like protein kinase-related family protein | | 3,3 ± 1,87 | 2,62 ± 15,88 | 105,72 ± 15,88 | 46,83 ± 3,97 |
| At1g55000 | peptidoglycan-binding LysM domain-containing protein | | 442,24 ± 27,99 | 296,37 ± 17,7 | 790,02 ± 17,7 | 337,89 ± 50,62 |
| At1g12580 | PEPKR1, phosphoenolpyruvate carboxylase-related kinase 1 | *PEPKR1* | 493,26 ± 13,99 | 461,02 ± 48,03 | 651,91 ± 48,03 | 443,57 ± 28,63 |
| At4g29140 | MATE efflux family protein | *ACTIVATED DISEASE SUSCEPTIBILITY 1 (ADS1)* | 335,3 ± 26,8 | 211,02 ± 73,36 | 292,79 ± 73,36 | 93,92 ± 6,39 |
| At5g13100 | Unknown protein | | 574,11 ± 28,15 | 599,88 ± 34,61 | 668,9 ± 34,61 | 512,57 ± 22,18 |
| At1g07280 | Tetratricopeptide repeat (TPR)-like superfamily protein | | 928,78 ± 23,33 | 395,92 ± 44,62 | 1105,19 ± 44,62 | 268,69 ± 33,33 |
| At1g02520 | PGP11, P-glycoprotein 11 | *ATP-BINDING CASSETTE B11 (ABCB11)* | 83,34 ± 31,26 | 445,33 ± 99,37 | 407,39 ± 99,37 | 487,52 ± 31,34 |
| At3g27750 | | *EMBRYO DEFECTIVE 3123 (EMB3123)* | 530,65 ± 35,27 | 460,08 ± 64,74 | 535,21 ± 64,74 | 288,45 ± 18,2 |
| At3g16190 | Isochorismatase family protein | | 335,18 ± 18,99 | 344,81 ± 23,02 | 424,02 ± 23,02 | 326,13 ± 24,95 |
| At1g46554 | other RNA | | 103,4 ± 10,97 | 71,53 ± 11,62 | 134,19 ± 11,62 | 57,28 ± 7,64 |
| At1g74070 | Cyclophilin-like peptidyl-prolyl cis-trans isomerase family protein | | 376,06 ± 33,64 | 294,79 ± 137,59 | 492,41 ± 137,59 | 192,31 ± 20,61 |
| At2g35960 | NHL12, NDR1/HIN1-like 12 | *NDR1/HIN1-LIKE 12 (NHL12)* | 115,99 ± 10,5 | 106,45 ± 77,36 | 277,52 ± 77,36 | 88,6 ± 9,21 |
| At4g01050 | TROL, thylakoid rhodanese-like | *THYLAKOID RHODANESE-LIKE (TROL)* | 6704,4 ± 494,36 | 6263,96 ± 389,91 | 8246,66 ± 389,91 | 5290,93 ± 645,18 |
| At5g59060 | BEST Arabidopsis thaliana protein match is: RNA-directed DNA polymerase (reverse transcriptase)-related family protein | | 0,49 ± 0,86 | 0,69 ± 1,61 | 5,28 ± 1,61 | 5,55E-17 ± 7,55E-33 |

**Supplementary Table S1. UV-B responsive genes regulated by TGA2/5/6 factors.** List of genes differentially regulated by the interaction between treatment and genotype.

| | | | Normalized expression | | | |
|---|---|---|---|---|---|---|
| | | | *Wild type* | | *tga256* | |
| Locus | Gene model Description | Gene Symbol | Control | UV-B | Control | UV-B |
| | ***Cluster 2*** | | | | | |
| At1g30120 | PDH-E1 BETA, pyruvate dehydrogenase E1 beta | *PYRUVATE DEHYDROGENASE E1 BETA (PDH-E1 BETA)* | 1919,26 ± 48,22 | 1662,85 ± 175,77 | 2034,12 ± 175,77 | 1425,74 ± 12,3 |
| At4g37540 | LBD39, LOB domain-containing protein 39 | *LOB DOMAIN-CONTAINING PROTEIN 39 (LBD39)* | 615,14 ± 43,45 | 251,02 ± 106,24 | 540,12 ± 106,24 | 99,05 ± 6,93 |
| At3g60390 | HAT3, homeobox-leucine zipper protein 3 | *HOMEOBOX-LEUCINE ZIPPER PROTEIN 3 (HAT3)* | 177,55 ± 23,37 | 67,73 ± 14,92 | 195,67 ± 14,92 | 36,5 ± 2,51 |
| At3g47390 | PHS1, cytidine/deoxycytidylate deaminase family protein | *PHOTOSENSITIVE 1 (PHS1)* | 314,41 ± 8,57 | 298,19 ± 9,51 | 376,34 ± 9,51 | 294,78 ± 11,05 |
| At3g03990 | alpha/beta-Hydrolases superfamily protein | | 470,25 ± 42,86 | 653,05 ± 36,52 | 729,89 ± 36,52 | 650,67 ± 59,39 |
| At1g66350 | RGL, RGL1, RGA-like 1 | *RGA-LIKE 1 (RGL1)* | 164,49 ± 24,24 | 78,02 ± 17,2 | 230,81 ± 17,2 | 56,67 ± 14,99 |
| At1g19960 | BEST Arabidopsis thaliana protein match is: transmembrane receptors (TAIR:AT2G32140.1) | | 143,86 ± 108,45 | 82,14 ± 34,71 | 300,11 ± 34,71 | 26,39 ± 5,52 |
| At1g52120 | Mannose-binding lectin superfamily protein | | 5,83 ± 2,21 | 1,99 ± 20,16 | 153,85 ± 20,16 | 47,33 ± 5,39 |
| At4g26555 | FKBP-like peptidyl-prolyl cis-trans isomerase family protein | | 437,4 ± 32,5 | 338,19 ± 34,9 | 651,02 ± 34,9 | 340,73 ± 33,87 |
| At3g08010 | ATAB2, RNA binding | *(ATAB2)* | 938,16 ± 121,42 | 703,01 ± 72,21 | 1221,5 ± 72,21 | 579,89 ± 91,89 |
| At1g55510 | BCDH BETA1, branched-chain alpha-keto acid decarboxylase E1 beta subunit | *BRANCHED-CHAIN ALPHA-KETO ACID DECARBOXYLASE E1 BETA SUBUNIT (BCDH BETA1)* | 230,89 ± 15,42 | 210,77 ± 36,85 | 294,1 ± 36,85 | 187,62 ± 15,68 |
| At5g04880 | pseudogene of ABC transporter family protein | | 122,71 ± 7,04 | 80,12 ± 14,65 | 168,88 ± 14,65 | 71,39 ± 11,73 |
| At4g37200 | HCF164, Thioredoxin superfamily protein | *HIGH CHLOROPHYLL FLUORESCENCE 164 (HCF164)* | 549,84 ± 52,28 | 560,95 ± 76,27 | 696,81 ± 76,27 | 364,94 ± 31,05 |
| At1g03340 | Unknown protein | | 34,51 ± 9,62 | 33,28 ± 18,53 | 93,16 ± 18,53 | 32,38 ± 6,53 |
| At5g60700 | glycosyltransferase family protein 2 | | 425,02 ± 32,74 | 397,41 ± 13,73 | 525,24 ± 13,73 | 406,73 ± 6,97 |
| At4g22545 | pseudogene of unknown protein | | 5,5 ± 3,88 | 2,79 ± 3,8 | 19,59 ± 3,8 | 2,5 ± 2,25 |
| At3g16140 | PSAH-1, photosystem I subunit H-1 | *PHOTOSYSTEM I SUBUNIT H-1 (PSAH-1)* | 9993,32 ± 507,07 | 7626,18 ± 1504,38 | 14152,71 ± 1504,38 | 6542,29 ± 1259,45 |

**Supplementary Table S1. UV-B responsive genes regulated by TGA2/5/6 factors.** List of genes differentially regulated by the interaction between treatment and genotype.

| Locus | Gene model Description | Gene Symbol | Normalized expression | | | |
|---|---|---|---|---|---|---|
| | | | *Wild type* | | *tga256* | |
| | | | Control | UV-B | Control | UV-B |
| ***Cluster 2*** | | | | | | |
| At4g00150 | ATHAM3, HAM3, GRAS family transcription factor | *HAIRY MERISTEM 3 (HAM3)* | 407 ± 22,56 | 190,53 ± 14,87 | 528,41 ± 14,87 | 147,42 ± 25,56 |
| At1g01600 | CYP86A4, cytochrome P450, family 86, subfamily A, polypeptide 4 | *CYTOCHROME P45, SUBFAMILY A, POLYPEPTIDE 4 (CYP86A4)* | 63,42 ± 8,56 | 53,03 ± 17,08 | 125,74 ± 17,08 | 45,31 ± 6,98 |
| At2g31450 | ATNTH1, DNA glycosylase superfamily protein | *(ATNTH1)* | 134,76 ± 9,61 | 134,17 ± 22,25 | 157,43 ± 22,25 | 97,8 ± 4,4 |
| At3g15355 | PFU1, UBC25, ubiquitin-conjugating enzyme 25 | *UBIQUITIN-CONJUGATING ENZYME 25 (UBC25)* | 270,19 ± 12,99 | 373,66 ± 30,29 | 413,73 ± 30,29 | 452,41 ± 13,48 |
| At2g43540 | Unknown protein | | 190,94 ± 22,47 | 227,48 ± 13,85 | 344,27 ± 13,85 | 287,57 ± 16,91 |
| At1g75340 | Zinc finger C-x8-C-x5-C-x3-H type family protein | | 249,89 ± 15,62 | 274,6 ± 13,5 | 325,19 ± 13,5 | 278,66 ± 17,47 |
| At3g56630 | CYP94D2, cytochrome P450, family 94, subfamily D, polypeptide 2 | *CYTOCHROME P45, FAMILY 94, SUBFAMILY D, POLYPEPTIDE 2 (CYP94D2)* | 243,48 ± 6,89 | 219,86 ± 39,21 | 325,63 ± 39,21 | 199,03 ± 7,28 |
| At3g07522 | Unknown protein | | 0,26 ± 0,45 | 5,55E-17 ± 0,77 | 2,5 ± 0,77 | 5,55E-17 ± 7,55E-33 |
| At5g27650 | Tudor/PWWP/MBT superfamily protein | | 496,78 ± 15,67 | 421,35 ± 18,09 | 586,19 ± 18,09 | 381,91 ± 30,43 |
| At1g03140 | splicing factor Prp18 family protein | | 810,27 ± 33,72 | 743,1 ± 65,16 | 1114,41 ± 65,16 | 792,19 ± 27,78 |
| At4g04950 | thioredoxin family protein | *MONOTHIOL GLUTAREDOXIN 17 (GRXS17)* | 588,44 ± 6,86 | 528,47 ± 16,56 | 564,69 ± 16,56 | 439,1 ± 18,19 |
| At3g45660 | Major facilitator superfamily protein | | 9,39 ± 3,49 | 14,84 ± 3,92 | 36,15 ± 3,92 | 16,49 ± 1,97 |
| At2g43550 | Scorpion toxin-like knottin superfamily protein | | 328,07 ± 35,97 | 275 ± 105,08 | 430,87 ± 105,08 | 182,91 ± 24,32 |
| At5g42530 | Unknown protein | | 1629,01 ± 1145,59 | 6156,15 ± 3318,58 | 20691,07 ± 3318,58 | 13115,52 ± 954,17 |
| At2g32100 | ATOFP16, OFP16, ovate family protein 16 | *OVATE FAMILY PROTEIN 16 (OFP16)* | 122,38 ± 23,94 | 20,05 ± 100,01 | 383,57 ± 100,01 | 27,76 ± 14,51 |
| At1g60000 | RNA-binding (RRM/RBD/RNP motifs) family protein | | 863,77 ± 165,33 | 324,35 ± 213,71 | 1101,09 ± 213,71 | 199,6 ± 35,36 |
| At1g12050 | fumarylacetoacetase, putative | | 531,96 ± 46,91 | 794,23 ± 30,31 | 784,51 ± 30,31 | 786,44 ± 74,57 |
| At3g22210 | Unknown protein | | 616,97 ± 72,95 | 595,81 ± 54,33 | 719,47 ± 54,33 | 420,59 ± 47,28 |
| At5g55790 | Unknown protein | | 144,33 ± 5,45 | 144,3 ± 3,75 | 143,8 ± 3,75 | 111,61 ± 4,05 |

**Supplementary Table S1. UV-B responsive genes regulated by TGA2/5/6 factors.** List of genes differentially regulated by the interaction between treatment and genotype.

| Locus | Gene model Description | Gene Symbol | Normalized expression | | | |
|---|---|---|---|---|---|---|
| | | | *Wild type* | | *tga256* | |
| | | | Control | UV-B | Control | UV-B |
| | ***Cluster 2*** | | | | | |
| At5g62890 | Xanthine/uracil permease family protein | | 1841,81 ± 202,66 | 1444,84 ± 131,36 | 2148,18 ± 131,36 | 1180,93 ± 59,93 |
| At4g14390 | Ankyrin repeat family protein | | 15,98 ± 2,44 | 29,41 ± 11,86 | 75,84 ± 11,86 | 56,47 ± 10,4 |
| At4g11900 | S-locus lectin protein kinase family protein | | 99,98 ± 22,1 | 177,54 ± 25,67 | 329,15 ± 25,67 | 313,44 ± 15,71 |
| At1g18490 | Protein of unknown function (DUF1637) | | 148,62 ± 17,47 | 104,14 ± 5,33 | 172,11 ± 5,33 | 83,93 ± 6,02 |
| At2g26870 | NPC2, non-specific phospholipase C2 | *NON-SPECIFIC PHOSPHOLIPASE C2 (NPC2)* | 74,52 ± 5,78 | 70,43 ± 15,87 | 141,55 ± 15,87 | 68,71 ± 3,06 |
| At5g43470 | HRT, RCY1, RPP8, Disease resistance protein (CC-NBS-LRR class) family | *RECOGNITION OF PERONOSPORA PARASITICA 8 (RPP8)* | 1034,5 ± 35,42 | 1494,61 ± 56,25 | 1609,97 ± 56,25 | 1594,5 ± 147,88 |
| At3g57500 | Unknown protein | | 17,51 ± 2,55 | 18,98 ± 7,58 | 38,78 ± 7,58 | 19,68 ± 3,18 |
| At3g27950 | GDSL-like Lipase/Acylhydrolase superfamily protein | | 7,06 ± 3,13 | 5,36 ± 6,7 | 60,78 ± 6,7 | 23 ± 11,86 |
| At5g01650 | Tautomerase/MIF superfamily protein | | 437,36 ± 10,39 | 550,92 ± 40,45 | 605,86 ± 40,45 | 564,19 ± 45,76 |
| At3g09160 | RNA-binding (RRM/RBD/RNP motifs) family protein | | 21,32 ± 10,06 | 20,81 ± 4,19 | 45,69 ± 4,19 | 17,81 ± 3,04 |
| At5g07840 | Ankyrin repeat family protein | *PHYTOCHROME INTERACTING ANKYRIN-REPEAT PROTEIN 1 (PIA1)* | 290,07 ± 27,37 | 221,33 ± 16,54 | 384,04 ± 16,54 | 185,68 ± 33,73 |
| At5g58260 | oxidoreductases, acting on NADH or NADPH, quinone or similar compound as acceptor | *NADH DEHYDROGENASE-LIKE COMPLEX N (NdhN)* | 356,99 ± 91,82 | 388,19 ± 29,41 | 686,8 ± 29,41 | 398,1 ± 53,92 |
| At1g74910 | ADP-glucose pyrophosphorylase family protein | | 1743,47 ± 84 | 1125,03 ± 89,12 | 1838,87 ± 89,12 | 885,82 ± 12,34 |
| At2g31190 | RUS2, WXR1, Protein of unknown function, DUF647 | *ROOT UV-B SENSITIVE 2 (RUS2)* | 174,21 ± 12,29 | 134,71 ± 4,85 | 184,67 ± 4,85 | 95,32 ± 2,05 |
| At2g02910 | Protein of unknown function (DUF616) | | 149,58 ± 1,72 | 135,92 ± 6,43 | 173,8 ± 6,43 | 92,23 ± 13,33 |
| At1g18265 | Protein of unknown function, DUF593 | | 16,34 ± 6,52 | 13,64 ± 12,88 | 47,31 ± 12,88 | 10,38 ± 2,49 |
| At3g10850 | GLX2-2, GLY2, Metallo-hydrolase/oxidoreductase superfamily protein | *(GLY2)* | 1545,98 ± 58,11 | 921,86 ± 75,62 | 1716,68 ± 75,62 | 799,53 ± 21,07 |
| AtCg00310 | TRNG.2, tRNA-Gly | *(TRNG.2)* | 5,55E-17 ± 7,55E-33 | 5,55E-17 ± 0,76 | 1,8 ± 0,76 | 5,55E-17 ± 7,55E-33 |
| At3g16130 | ATROPGEF13, PIRF2, ROPGEF13, RHO guanyl-nucleotide exchange factor 13 | *ROPGEF13* | 3287,47 ± 396,69 | 2998,01 ± 725,6 | 5142,24 ± 725,6 | 2672,02 ± 432,03 |

**Supplementary Table S1. UV-B responsive genes regulated by TGA2/5/6 factors.** List of genes differentially regulated by the interaction between treatment and genotype.

| Locus | Gene model Description | Gene Symbol | Normalized expression | | | |
|---|---|---|---|---|---|---|
| | | | *Wild type* | | *tga256* | |
| | | | Control | UV-B | Control | UV-B |
| | ***Cluster 2*** | | | | | |
| At2g40750 | ATWRKY54, WRKY54, WRKY DNA-binding protein 54 | *WRKY54* | 27,2 ± 27,26 | 173,58 ± 15,19 | 290,51 ± 15,19 | 257,66 ± 31,38 |
| At3g06080 | TBL10, Plant protein of unknown function (DUF828) | | 865,89 ± 19,95 | 224,3 ± 11,52 | 891,51 ± 11,52 | 163,44 ± 13,68 |
| At5g66005 | Expressed protein | | 81,27 ± 4,81 | 73,38 ± 12,62 | 118,35 ± 12,62 | 65,86 ± 6,37 |
| At2g43280 | Far-red impaired responsive (FAR1) family protein | | 85,2 ± 10,85 | 67,14 ± 2,61 | 84,5 ± 2,61 | 39,85 ± 6,42 |
| At4g27610 | Unknown protein | | 281,24 ± 19,99 | 170,62 ± 24,55 | 305,93 ± 24,55 | 130,62 ± 6,63 |
| At2g36220 | Unknown protein | | 194,18 ± 37,14 | 421,38 ± 74,03 | 472,74 ± 74,03 | 557,19 ± 40,5 |
| At5g54530 | Protein of unknown function, DUF538 | | 58,41 ± 7,98 | 18,32 ± 3,46 | 93,64 ± 3,46 | 17,03 ± 1,92 |
| At4g14400 | ACD6, ankyrin repeat family protein | *ACCELERATED CELL DEATH 6 (ACD6)* | 522,25 ± 155,84 | 1280,83 ± 1023,35 | 3601,62 ± 1023,35 | 2194,96 ± 232,3 |
| At3g01740 | Mitochondrial ribosomal protein L37 | | 506,65 ± 29,36 | 407,13 ± 36,74 | 576,03 ± 36,74 | 342,67 ± 9,31 |
| At3g55790 | Unknown protein | | 5,55E-17 ± 7,55E-33 | 5,55E-17 ± 1,4 | 3,12 ± 1,4 | 5,55E-17 ± 7,55E-33 |
| At1g44050 | Cysteine/Histidine-rich C1 domain family protein | | 28,12 ± 10,58 | 1,61 ± 25,55 | 117,47 ± 25,55 | 16,45 ± 5 |
| At5g44520 | NagB/RpiA/CoA transferase-like superfamily protein | | 276,07 ± 11 | 221,64 ± 13,78 | 287,66 ± 13,78 | 157,2 ± 11,29 |
| At3g20060 | UBC19, ubiquitin-conjugating enzyme19 | *UBC19* | 771,99 ± 35,34 | 819,79 ± 60,8 | 981,57 ± 60,8 | 807,68 ± 15,81 |
| At1g78460 | SOUL heme-binding family protein | | 1193,58 ± 252,87 | 810,95 ± 162,5 | 1588,07 ± 162,5 | 346,99 ± 52,1 |
| At5g19950 | Domain of unknown function (DUF1767) | | 358,6 ± 28,01 | 301,46 ± 54,98 | 412,78 ± 54,98 | 220,51 ± 18,29 |
| At3g59010 | PME61, pectin methylesterase 61 | *PECTIN METHYLESTERASE 61 (PME61)* | 54,48 ± 13,69 | 27,61 ± 20,76 | 145,1 ± 20,76 | 41,42 ± 8,71 |
| At5g13570 | ATDCP2, DCP2, TDT, decapping 2 | *DECAPPING 2 (DCP2)* | 314,63 ± 38,87 | 260,1 ± 13,81 | 349,49 ± 13,81 | 197,37 ± 17,54 |
| At3g02430 | Protein of unknown function (DUF679) | | 0,49 ± 0,42 | 5,55E-17 ± 1,11 | 3,52 ± 1,11 | 5,55E-17 ± 7,55E-33 |
| At5g64330 | JK218, NPH3, RPT3, Phototropic-responsive NPH3 family protein | *nPH3* | 836,21 ± 67,38 | 361,12 ± 59,62 | 831,56 ± 59,62 | 222,39 ± 36,67 |
| At4g18160 | ATKCO6, ATTPK3, KCO6, TPK3, Ca2+ activated outward rectifying K+ channel 6 | *CA2+ ACTIVATED OUTWARD RECTIFYING K+ CHANNEL 6 (KCO6)* | 138,46 ± 6,7 | 128,3 ± 24,85 | 245,85 ± 24,85 | 144,37 ± 20,32 |
| At3g26300 | CYP71B34, cytochrome P450, family 71, subfamily B, polypeptide 34 | *CYP71B34* | 533,63 ± 24,04 | 408,95 ± 39,07 | 402,85 ± 39,07 | 226,1 ± 6,95 |
| At4g10910 | Unknown protein | | 16,03 ± 7,18 | 3,47 ± 17,68 | 53,94 ± 17,68 | 2,2 ± 1,94 |

**Supplementary Table S1. UV-B responsive genes regulated by TGA2/5/6 factors.** List of genes differentially regulated by the interaction between treatment and genotype.

| | | | Normalized expression | | | |
|---|---|---|---|---|---|---|
| | | | *Wild type* | | *tga256* | |
| Locus | Gene model Description | Gene Symbol | Control | UV-B | Control | UV-B |
| ***Cluster 2*** | | | | | | |
| At2g24710 | ATGLR2.3, GLR2.3, glutamate receptor 2.3 | *GLUTAMATE RECEPTOR 2.3 (GLR2.3)* | 2,54 ± 1,12 | 5,55E-17 ± 2,24 | 17,24 ± 2,24 | 5,73 ± 3,07 |
| At3g53700 | MEE40, Pentatricopeptide repeat (PPR) superfamily protein | *MATERNAL EFFECT EMBRYO ARREST 4 (MEE4)* | 344,44 ± 18,6 | 205,38 ± 26,88 | 329,17 ± 26,88 | 150,47 ± 9,54 |
| At3g61600 | ATPOB1, POB1, POZ/BTB containin G-protein 1 | *POZ/BTB CONTAININ G-PROTEIN 1 (POB1)* | 1471,52 ± 45,24 | 1221,89 ± 153,98 | 2006,25 ± 153,98 | 1312,41 ± 82,34 |
| At1g79790 | Haloacid dehalogenase-like hydrolase (HAD) superfamily protein | *FLAVIN MONONUCLEOTIDE HYDROLASE 1 (FHY1)* | 511,06 ± 38,29 | 443,2 ± 12,36 | 685,16 ± 12,36 | 393,79 ± 73,02 |
| At1g05030 | Major facilitator superfamily protein | | 244,45 ± 37,96 | 172,34 ± 26,99 | 405,11 ± 26,99 | 189,48 ± 24,76 |
| At3g61198 | other RNA | | 32,17 ± 10,37 | 36,47 ± 11,58 | 115,8 ± 11,58 | 46,82 ± 9,53 |
| At4g12800 | PSAL, photosystem I subunit l | *PHOTOSYSTEM I SUBUNIT L (PSAL)* | 33302,21 ± 4828,96 | 25083,55 ± 5837,66 | 50084,2 ± 5837,66 | 21162,37 ± 2884,27 |
| At4g23490 | Protein of unknown function (DUF604) | | 185,46 ± 13,68 | 131,29 ± 12,74 | 181,37 ± 12,74 | 79,29 ± 10,22 |
| At1g22430 | GroES-like zinc-binding dehydrogenase family protein | | 489,69 ± 23,58 | 539,24 ± 71,05 | 686,25 ± 71,05 | 461,49 ± 36,69 |
| At5g50060 | Plant invertase/pectin methylesterase inhibitor superfamily protein | | 1,14 ± 0,43 | 5,55E-17 ± 0,74 | 3,15 ± 0,74 | 5,55E-17 ± 7,55E-33 |
| At3g51150 | ATP binding microtubule motor family protein | | 265,67 ± 8,76 | 199,84 ± 49,7 | 416,62 ± 49,7 | 188,8 ± 9,18 |
| At1g67440 | emb1688, Minichromosome maintenance (MCM2/3/5) family protein | *EMBRYO DEFECTIVE 1688 (emb1688)* | 235,66 ± 4,89 | 172,17 ± 19,89 | 266,92 ± 19,89 | 143,51 ± 10,39 |
| At3g15900 | Unknown protein | | 163,72 ± 14,13 | 88,61 ± 24,31 | 273,01 ± 24,31 | 84,31 ± 2,81 |
| At4g25910 | ATCNFU3, NFU3, NFU domain protein 3 | *NFU DOMAIN PROTEIN 3 (NFU3)* | 292,34 ± 21,23 | 115,63 ± 37,81 | 498,34 ± 37,81 | 118,3 ± 11,7 |
| At4g26530 | Aldolase superfamily protein | *FRUCTOSE-BISPHOSPHATE ALDOLASE 5 (FBA5)* | 1346,83 ± 636,06 | 1345,06 ± 2170,49 | 5881,12 ± 2170,49 | 1171,95 ± 396,57 |
| At5g55700 | BAM4, BMY6, beta-amylase 4 | *BETA-AMYLASE 4 (BAM4)* | 352,01 ± 3,01 | 554,64 ± 65,98 | 544,16 ± 65,98 | 569,2 ± 21,65 |
| At3g55080 | SET domain-containing protein | | 241,64 ± 12,41 | 195,63 ± 20,68 | 266,2 ± 20,68 | 169,45 ± 7,33 |
| At5g14970 | Unknown protein | | 962,66 ± 54,17 | 925,29 ± 77,61 | 1198,86 ± 77,61 | 729,42 ± 101,39 |
| At5g07842 | CPuORF16, conserved peptide upstream open reading frame 16 | *CPuORF16* | 291,98 ± 26,84 | 222,03 ± 16 | 384,73 ± 16 | 185,68 ± 33,73 |

**Supplementary Table S1. UV-B responsive genes regulated by TGA2/5/6 factors.** List of genes differentially regulated by the interaction between treatment and genotype.

| Locus | Gene model Description | Gene Symbol | Normalized expression | | | |
|---|---|---|---|---|---|---|
| | | | ***Wild type*** | | ***tga256*** | |
| **Locus** | **Gene model Description** | **Gene Symbol** | **Control** | **UV-B** | **Control** | **UV-B** |
| ***Cluster 2*** | | | | | | |
| At3g11930 | Adenine nucleotide alpha hydrolases-like superfamily protein | | 1643,42 ± 187,63 | 857,36 ± 216,96 | 2473,42 ± 216,96 | 585,48 ± 181,44 |
| At1g06680 | OE23, OEE2, PSBP-1, PSII-P, photosystem II subunit P-1 | *PHOTOSYSTEM II SUBUNIT P-1 (PSBP-1)* | 40041,44 ± 8315,02 | 27583,14 ± 9363,86 | 54618,9 ± 9363,86 | 20138,95 ± 2187,25 |
| At3g27925 | Deg1, DEGP1, DegP protease 1 | *DEGP PROTEASE 1 (DEGP1)* | 1307,21 ± 186,61 | 1187,77 ± 233,22 | 2029,81 ± 233,22 | 1276,36 ± 82,74 |
| At5g66570 | MSP-1, OE33, OEE1, OEE33, PSBO-1, PSBO1, PS II oxygen-evolving complex 1 | *PS II OXYGEN-EVOLVING COMPLEX 1 (PSBO1)* | 66132,31 ± 9080,54 | 46451,45 ± 12261,18 | 87613,26 ± 12261,18 | 35014,04 ± 5140,9 |
| At2g12190 | Cytochrome P450 superfamily protein | | 16,45 ± 2,08 | 16,38 ± 3,79 | 31,96 ± 3,79 | 12,64 ± 3,46 |
| At3g46490 | 2-oxoglutarate (2OG) and Fe(II)-dependent oxygenase superfamily protein | | 112,73 ± 14,25 | 60,24 ± 14,16 | 155,48 ± 14,16 | 33,19 ± 6,99 |
| At2g19860 | ATHXK2, HXK2, hexokinase 2 | *HEXOKINASE 2 (HXK2)* | 389,58 ± 25,59 | 344,13 ± 11,93 | 342 ± 11,93 | 225,38 ± 13,61 |
| At5g55720 | Pectin lyase-like superfamily protein | | 2,54 ± 2,2 | 1,63 ± 6,75 | 23,36 ± 6,75 | 3,5 ± 2,91 |
| At4g08093 | pseudogene of unknown protein | | 1,38 ± 0,09 | 0,34 ± 0,64 | 4,92 ± 0,64 | 5,55E-17 ± 7,55E-33 |
| At1g06980 | Unknown protein | | 21,61 ± 2,96 | 10,08 ± 1,97 | 26,59 ± 1,97 | 5,2 ± 2,18 |
| At1g31335 | Unknown protein | | 94,12 ± 11,15 | 39,87 ± 12,48 | 110,98 ± 12,48 | 19,34 ± 6,8 |
| At4g38460 | GGR, geranylgeranyl reductase | *GERANYLGERANYL REDUCTASE (GGR)* | 1038,42 ± 68,16 | 651,18 ± 77,87 | 1361,39 ± 77,87 | 663,11 ± 20,27 |
| At2g03210 | ATFUT2, FUT2, fucosyltransferase 2 | *FUCOSYLTRANSFERASE 2 (FUT2)* | 0,25 ± 0,44 | 2,74 ± 0,74 | 3,15 ± 0,74 | 0,42 ± 0,68 |
| At5g16950 | Unknown protein | | 58,65 ± 11,15 | 65,06 ± 3,21 | 56,06 ± 3,21 | 32,18 ± 6,66 |
| At3g19080 | SWIB complex BAF60b domain-containing protein | | 101,64 ± 18,88 | 110,08 ± 7,54 | 126,76 ± 7,54 | 78,01 ± 4,04 |
| ***Cluster 3*** | | | | | | |
| At3g02750 | Protein phosphatase 2C family protein | | 881,63 ± 16,54 | 1312,04 ± 71,12 | 933,64 ± 71,12 | 1038,42 ± 20,13 |
| At1g21110 | O-methyltransferase family protein | *INDOLE GLUCOSINOLATE METHYLTRANSFERASE 3 (IGMT3)* | 116,91 ± 21,53 | 1393,75 ± 105,5 | 318,32 ± 105,5 | 1060,46 ± 108,47 |
| At5g41750 | Disease resistance protein (TIR-NBS-LRR class) family | | 39,24 ± 16,73 | 973,19 ± 25,53 | 210,3 ± 25,53 | 1860,53 ± 66,85 |
| At3g26850 | histone-lysine N-methyltransferases | | 153,91 ± 16,74 | 458,45 ± 16,67 | 110,57 ± 16,67 | 703,51 ± 183,82 |
| At4g20860 | FAD-binding Berberine family protein | | 339,62 ± 56,04 | 3143,17 ± 122,85 | 512,85 ± 122,85 | 1901,69 ± 203 |

**Supplementary Table S1. UV-B responsive genes regulated by TGA2/5/6 factors.** List of genes differentially regulated by the interaction between treatment and genotype.

| Locus | Gene model Description | Gene Symbol | Normalized expression | | | |
|---|---|---|---|---|---|---|
| | | | *Wild type* | | *tga256* | |
| | | | Control | UV-B | Control | UV-B |
| ***Cluster 3*** | | | | | | |
| At5g54855 | Pollen Ole e 1 allergen and extensin family protein | | 195,24 ± 10,68 | 286,84 ± 14,07 | 227,27 ± 14,07 | 220,24 ± 7,46 |
| At1g73500 | ATMKK9, MKK9, MAP kinase kinase 9 | *MAP KINASE KINASE 9 (MKK9)* | 155,98 ± 20,17 | 676,18 ± 15,47 | 272,69 ± 15,47 | 491,29 ± 61,12 |
| At1g12640 | MBOAT (membrane bound O-acyl transferase) family protein | *LYSOPHOSPHOLIPID ACYLTRANSFERASE 1 (LPLAT1)* | 468,13 ± 34,02 | 712,12 ± 26,49 | 627,96 ± 26,49 | 736,47 ± 26,78 |
| At5g24460 | Unknown protein | | 421,67 ± 24,29 | 933,1 ± 35,74 | 684,83 ± 35,74 | 985,03 ± 78,4 |
| At3g11840 | PUB24, plant U-box 24 | *PLANT U-BOX 24 (PUB24)* | 46,28 ± 8,17 | 624,58 ± 15,81 | 112,83 ± 15,81 | 750,7 ± 64,52 |
| At2g11520 | CRCK3, calmodulin-binding receptor-like cytoplasmic kinase 3 | *CALMODULIN-BINDING RECEPTOR-LIKE CYTOPLASMIC KINASE 3 (CRCK3)* | 374,36 ± 15,3 | 1219,2 ± 18,43 | 327,23 ± 18,43 | 1376,2 ± 80,27 |
| At1g27900 | RNA helicase family protein | | 307,2 ± 6,47 | 389,53 ± 16,72 | 342,18 ± 16,72 | 364,75 ± 8,81 |
| At3g47780 | ATATH6, ATH6, ABC2 homolog 6 | *ATP-BINDING CASSETTE A7 (ABCA7)* | 139,93 ± 9,75 | 1182,67 ± 45,16 | 231,02 ± 45,16 | 965,15 ± 105,01 |
| At2g34080 | Cysteine proteinases superfamily protein | | 74,08 ± 15,55 | 518,64 ± 3,57 | 13,29 ± 3,57 | 556,79 ± 195,12 |
| At1g03730 | Unknown protein | | 131,04 ± 15,24 | 275,77 ± 17,66 | 96,95 ± 17,66 | 376,51 ± 1,79 |
| At4g27290 | S-locus lectin protein kinase family protein | | 43,44 ± 4,46 | 124,65 ± 1,73 | 22,7 ± 1,73 | 192,73 ± 26,47 |
| At5g48540 | receptor-like protein kinase-related family protein | | 58,49 ± 35,88 | 1285,43 ± 95,57 | 172,66 ± 95,57 | 767,62 ± 40,88 |
| At5g58840 | Subtilase family protein | | 2,57 ± 1,06 | 54,55 ± 1,42 | 7,01 ± 1,42 | 26,67 ± 9,13 |
| At1g28240 | Protein of unknown function (DUF616) | | 556,94 ± 31,19 | 760,76 ± 23,34 | 529,95 ± 23,34 | 954,13 ± 34,5 |
| At1g79500 | AtkdsA1, Aldolase-type TIM barrel | *(AtkdsA1)* | 640,18 ± 34,61 | 1316,37 ± 28,1 | 807,77 ± 28,1 | 1195,38 ± 91,66 |
| At1g73880 | UGT89B1, UDP-glucosyl transferase 89B1 | *UGT89B1* | 120,56 ± 14,2 | 210,71 ± 13,3 | 156,21 ± 13,3 | 164,51 ± 25,86 |
| At1g11330 | S-locus lectin protein kinase family protein | | 314,51 ± 26,84 | 2008,34 ± 16,99 | 463,91 ± 16,99 | 1845,47 ± 154,89 |
| At3g08870 | Concanavalin A-like lectin protein kinase family protein | | 15 ± 5,71 | 115,32 ± 19,98 | 57,2 ± 19,98 | 102,09 ± 23,2 |
| At1g27730 | STZ, ZAT10, salt tolerance zinc finger | *SALT TOLERANCE ZINC FINGER (STZ)* | 48,07 ± 16,12 | 3204,31 ± 40,2 | 206,31 ± 40,2 | 3653,58 ± 383,52 |
| At2g46100 | Nuclear transport factor 2 family protein | | 356,82 ± 9,78 | 629,15 ± 53,65 | 479,86 ± 53,65 | 558,81 ± 77,71 |
| At1g30190 | Unknown protein | | 2,14 ± 2,27 | 42,81 ± 1,09 | 1,11 ± 1,09 | 99,9 ± 19,7 |

**Supplementary Table S1. UV-B responsive genes regulated by TGA2/5/6 factors.** List of genes differentially regulated by the interaction between treatment and genotype.

| | | | Normalized expression | | | |
|---|---|---|---|---|---|---|
| | | | *Wild type* | | *tga256* | |
| Locus | Gene model Description | Gene Symbol | Control | UV-B | Control | UV-B |
| | ***Cluster 3*** | | | | | |
| At1g02930 | ATGST1, ATGSTF3, ATGSTF6, ERD11, GST1, GSTF6, glutathione S-transferase 6 | *GSTF6* | 877,62 ± 215,09 | 38169,79 ± 1471,96 | 4306,32 ± 1471,96 | 45803,22 ± 5462,87 |
| At4g25380 | SAP10, stress-associated protein 10 | *STRESS-ASSOCIATED PROTEIN 1 (SAP1)* | 0,49 ± 0,86 | 44,17 ± 1,18 | 1,38 ± 1,18 | 16,3 ± 7,41 |
| At1g66570 | ATSUC7, SUC7, sucrose-proton symporter 7 | *SUCROSE-PROTON SYMPORTER 7 (SUC7)* | 0,96 ± 0,83 | 203,77 ± 7,55E-33 | 5,55E-17 ± 7,55E-33 | 69,27 ± 14,31 |
| At2g31820 | Ankyrin repeat family protein | | 395,81 ± 26,52 | 828,68 ± 4,11 | 444,08 ± 4,11 | 650,36 ± 29,24 |
| At3g22060 | Receptor-like protein kinase-related family protein | | 118,33 ± 33,42 | 2299,23 ± 156,93 | 372,95 ± 156,93 | 1792,63 ± 123,65 |
| At1g26410 | FAD-binding Berberine family protein | | 5,12 ± 4,5 | 279,24 ± 30,6 | 80 ± 30,6 | 382,16 ± 32,94 |
| At5g22520 | Unknown protein | | 1,71 ± 1,58 | 413,08 ± 2,56 | 2,95 ± 2,56 | 590,49 ± 32,45 |
| At3g28160 | transposable element gene | | 194,56 ± 24,85 | 1006,86 ± 23,01 | 88,22 ± 23,01 | 847,17 ± 13,74 |
| At5g02270 | ATNAP9, NAP9, non-intrinsic ABC protein 9 | *ATP-BINDING CASSETTE I2 (ABCI2)* | 386,19 ± 48,9 | 2568,3 ± 34,11 | 217,8 ± 34,11 | 2423,76 ± 83,48 |
| At1g20310 | Unknown protein | | 0,98 ± 1,23 | 278,14 ± 0,7 | 1,43 ± 0,7 | 202,02 ± 17,8 |
| At3g50790 | esterase/lipase/thioesterase family protein | | 389,96 ± 22,61 | 631,39 ± 42,21 | 474,47 ± 42,21 | 504,76 ± 41,4 |
| At3g55950 | ATCRR3, CCR3, CRINKLY4 related 3 | *CRINKLY4 RELATED 3 (CCR3)* | 87,84 ± 4,33 | 693,89 ± 12,25 | 89,34 ± 12,25 | 1069,82 ± 92,08 |
| At4g27820 | BGLU9, beta glucosidase 9 | *BETA GLUCOSIDASE 9 (BGLU9)* | 140,27 ± 6,28 | 461,96 ± 52,21 | 237,35 ± 52,21 | 381,81 ± 9,74 |
| At1g09340 | CRB, CSP41B, chloroplast RNA binding | *CHLOROPLAST RNA BINDING (CRB)* | 5901,04 ± 314,93 | 9591,53 ± 836,71 | 7421,98 ± 836,71 | 7936,08 ± 842,51 |
| At4g38980 | Unknown protein | | 94,33 ± 1,84 | 165,87 ± 1,07 | 111,56 ± 1,07 | 131,48 ± 17,78 |
| At2g32020 | Acyl-CoA N-acyltransferases (NAT) | | 9,73 ± 5,76 | 706,41 ± 7,22 | 59,76 ± 7,22 | 878,55 ± 48,05 |
| At5g14730 | Unknown protein | | 76,79 ± 18,32 | 612,36 ± 2,82 | 78,57 ± 2,82 | 398,1 ± 51,92 |
| At5g22545 | Unknown protein | | 0,49 ± 0,86 | 80,74 ± 0,63 | 0,36 ± 0,63 | 143,8 ± 22,31 |
| At5g22670 | F-box/RNI-like/FBD-like domains-containing protein | | 5,55E-17 ± 7,55E-33 | 7,49 ± 7,55E-33 | 5,55E-17 ± 7,55E-33 | 3,09 ± 1,49 |
| At1g02920 | ATGST11, ATGSTF7, ATGSTF8, glutathione S-transferase 7 | *GSTF7* | 657,73 ± 150,4 | 16779,88 ± 1382,93 | 3664,59 ± 1382,93 | 20837,17 ± 2651,07 |
| At4g00850 | GIF3, GRF1-interacting factor 3 | *GIF3)* | 52,68 ± 12,34 | 127,2 ± 1,34 | 41,74 ± 1,34 | 237,64 ± 21,01 |
| At3g56800 | acam-3, CAM3, calmodulin 3 | *CAM3* | 1944,3 ± 120,71 | 2541,23 ± 111,65 | 1679,63 ± 111,65 | 2865,35 ± 200,21 |

**Supplementary Table S1. UV-B responsive genes regulated by TGA2/5/6 factors.** List of genes differentially regulated by the interaction between treatment and genotype.

| | | | Normalized expression | | | |
|---|---|---|---|---|---|---|
| | | | *Wild type* | | *tga256* | |
| Locus | Gene model Description | Gene Symbol | Control | UV-B | Control | UV-B |
| ***Cluster 3*** | | | | | | |
| At3g05350 | Metallopeptidase M24 family protein | | 935,65 ± 10,43 | 3029,72 ± 105,89 | 1146,59 ± 105,89 | 2822,78 ± 103,06 |
| At4g02220 | zinc finger (MYND type) family protein / programmed cell death 2 C-terminal domain-containing protein | | 285,25 ± 7,18 | 423,01 ± 19,16 | 243,1 ± 19,16 | 481,4 ± 24,99 |
| At5g15400 | U-box domain-containing protein | | 999,5 ± 58,1 | 1214,83 ± 59,34 | 947,47 ± 59,34 | 1425,66 ± 73,32 |
| At4g18195 | ATPUP8, PUP8, purine permease 8 | *PURINE PERMEASE 8 (PUP8)* | 5,08 ± 4,26 | 115,46 ± 2,45 | 7,66 ± 2,45 | 76,5 ± 12,39 |
| At3g50950 | ZAR1, HOPZ-ACTIVATED RESISTANCE 1 | *HOPZ-ACTIVATED RESISTANCE 1 (ZAR1)* | 926,33 ± 29,35 | 3665,66 ± 253,1 | 2089,14 ± 253,1 | 4829,32 ± 76,34 |
| At1g76520 | Auxin efflux carrier family protein | | 440,25 ± 138,72 | 1827,16 ± 127,11 | 483,43 ± 127,11 | 771,83 ± 129,79 |
| At4g36530 | alpha/beta-Hydrolases superfamily protein | | 438,93 ± 13,18 | 1415,18 ± 45,28 | 688,51 ± 45,28 | 1290,24 ± 219,76 |
| At4g14910 | HISN5B, HISTIDINE BIOSYNTHESIS 5B | *HISTIDINE BIOSYNTHESIS 5B (HISN5B)* | 369,18 ± 20,48 | 559,86 ± 12,28 | 446,53 ± 12,28 | 467,83 ± 21,55 |
| At1g71520 | Integrase-type DNA-binding superfamily protein | | 2,34 ± 1,68 | 277,48 ± 2,5 | 1,46 ± 2,5 | 140,62 ± 30,42 |
| At1g65560 | Zinc-binding dehydrogenase family protein | | 83,02 ± 12,62 | 376,46 ± 4,83 | 58,97 ± 4,83 | 176,21 ± 12,03 |
| At1g79680 | ATWAKL10, WAKL10, WALL ASSOCIATED KINASE (WAK)-LIKE 10 | *WALL ASSOCIATED KINASE (WAK)-LIKE 1 (WAKL1)* | 7,23 ± 4,22 | 852,59 ± 9,89 | 45,37 ± 9,89 | 960,67 ± 89,7 |
| At4g15420 | Ubiquitin fusion degradation UFD1 family protein | | 224,14 ± 12,84 | 900,91 ± 16,23 | 316,71 ± 16,23 | 870,31 ± 28,39 |
| At1g20510 | OPCL1, OPC-8:0 CoA ligase1 | *OPC-8: COA LIGASE1 (OPCL1)* | 360,83 ± 61,68 | 2554,21 ± 22,16 | 620,73 ± 22,16 | 2588,16 ± 354,42 |
| At4g36430 | Peroxidase superfamily protein | | 213,62 ± 39,44 | 1599,57 ± 123,63 | 517,08 ± 123,63 | 1679,42 ± 205,74 |
| At1g01480 | ACS2, AT-ACC2, 1-amino-cyclopropane-1-carboxylate synthase 2 | *1-AMINO-CYCLOPROPANE-1-CARBOXYLATE SYNTHASE 2 (ACS2)* | 18,16 ± 5,6 | 1505,17 ± 2,98 | 11 ± 2,98 | 2314,15 ± 372,53 |
| At3g48195 | Phox (PX) domain-containing protein | | 373,63 ± 16,54 | 542,44 ± 15,1 | 468,53 ± 15,1 | 562,95 ± 37,21 |
| At2g43500 | Plant regulator RWP-RK family protein | | 104,83 ± 17,56 | 229,42 ± 17,56 | 165,08 ± 17,56 | 159,09 ± 15,74 |
| At2g47890 | B-box type zinc finger protein with CCT domain | | 430,06 ± 32,41 | 1901,48 ± 3,55 | 200,47 ± 3,55 | 1519,65 ± 235,75 |
| At5g37720 | ALY4, ALWAYS EARLY 4 | *ALWAYS EARLY 4 (ALY4)* | 609,37 ± 12,39 | 707 ± 22,26 | 595,72 ± 22,26 | 796,75 ± 41,88 |

**Supplementary Table S1. UV-B responsive genes regulated by TGA2/5/6 factors.** List of genes differentially regulated by the interaction between treatment and genotype.

| Locus | Gene model Description | Gene Symbol | Normalized expression | | | |
|---|---|---|---|---|---|---|
| | | | *Wild type* | | *tga256* | |
| | | | Control | UV-B | Control | UV-B |
| ***Cluster 3*** | | | | | | |
| At4g34380 | Transducin/WD40 repeat-like superfamily protein | | 10,13 ± 3,64 | 40,63 ± 0,11 | 1,06 ± 0,11 | 75,65 ± 11,09 |
| At4g02410 | Concanavalin A-like lectin protein kinase family protein | | 151,74 ± 26,2 | 1585,75 ± 50,42 | 265,81 ± 50,42 | 1331,88 ± 30,99 |
| At3g11270 | MEE34, Mov34/MPN/PAD-1 family protein | *(MEE34)* | 281,85 ± 10,21 | 578,38 ± 9,39 | 240,79 ± 9,39 | 613,12 ± 7,62 |
| At3g21400 | Unknown protein | | 126,45 ± 10,68 | 304,15 ± 14,29 | 122,86 ± 14,29 | 209,2 ± 9,19 |
| At3g17420 | GPK1, glyoxysomal protein kinase 1 | *GLYOXYSOMAL PROTEIN KINASE 1 (GPK1)* | 296,28 ± 5,79 | 489,4 ± 20,53 | 213,44 ± 20,53 | 650,69 ± 72,99 |
| At1g26420 | FAD-binding Berberine family protein | | 24,4 ± 14,72 | 1185,84 ± 45,3 | 132,65 ± 45,3 | 1341,02 ± 190,35 |
| At2g42350 | RING/U-box superfamily protein | | 63,32 ± 7,18 | 116,86 ± 8,94 | 49,23 ± 8,94 | 148,94 ± 5,12 |
| At4g23180 | CRK10, RLK4, cysteine-rich RLK (RECEPTOR-like protein kinase) 10 | *CYSTEINE-RICH RLK (RECEPTOR-LIKE PROTEIN KINASE) 1 (CRK1)* | 318,05 ± 39,67 | 2440,9 ± 94,6 | 716,01 ± 94,6 | 2786,2 ± 120,6 |
| At5g47170 | Unknown protein | | 5,55E-17 ± 7,55E-33 | 4,06 ± 0,76 | 1,8 ± 0,76 | 0,87 ± 1,42 |
| At5g65240 | Leucine-rich repeat protein kinase family protein | | 173,18 ± 5,88 | 353,85 ± 6,27 | 166,03 ± 6,27 | 274,38 ± 19,62 |
| At1g76040 | CPK29, calcium-dependent protein kinase 29 | *CALCIUM-DEPENDENT PROTEIN KINASE 29 (CPK29)* | 169,42 ± 29,48 | 952,01 ± 27,88 | 167,39 ± 27,88 | 1484,46 ± 66,37 |
| At4g14450 | Unknown protein | | 1,47 ± 1,59 | 82,53 ± 2,42 | 3,96 ± 2,42 | 141,43 ± 4,51 |
| At2g04160 | AIR3, Subtilisin-like serine endopeptidase family protein | *AUXIN-INDUCED IN ROOT CULTURES 3 (AIR3)* | 661,6 ± 58,01 | 1159,26 ± 91,9 | 757,5 ± 91,9 | 773,73 ± 83,89 |
| At4g11521 | Receptor-like protein kinase-related family protein | | 18,05 ± 1,36 | 672,03 ± 6,85 | 29,67 ± 6,85 | 594,76 ± 21,88 |
| At5g38990 | Malectin/receptor-like protein kinase family protein | | 369,92 ± 68,82 | 1214,01 ± 84,4 | 915,8 ± 84,4 | 1495,62 ± 46,64 |
| At5g27420 | ATL31, CNI1, carbon/nitrogen insensitive 1 | *CARBON/NITROGEN INSENSITIVE 1 (CNI1)* | 79,83 ± 17,7 | 1866,55 ± 63,21 | 314,64 ± 63,21 | 2319,8 ± 400,01 |
| At1g30640 | Protein kinase family protein | | 157,12 ± 7,28 | 242,97 ± 23,65 | 154,97 ± 23,65 | 360,79 ± 37,03 |
| At5g40000 | P-loop containing nucleoside triphosphate hydrolases superfamily protein | | 0,95 ± 0,83 | 178,92 ± 2,09 | 11,01 ± 2,09 | 144,81 ± 12,04 |
| At3g41768 | rRNA | | 5032,18 ± 1078,18 | 175967,9 ± 8490,97 | 50909,78 ± 8490,97 | 341448,5 ± 56462,06 |
| At2g44600 | Unknown protein | | 63,2 ± 10,93 | 117,41 ± 6,73 | 55,45 ± 6,73 | 198,49 ± 34,81 |

**Supplementary Table S1. UV-B responsive genes regulated by TGA2/5/6 factors.** List of genes differentially regulated by the interaction between treatment and genotype.

| | | | Normalized expression | | | |
|---|---|---|---|---|---|---|
| | | | *Wild type* | | *tga256* | |
| Locus | Gene model Description | Gene Symbol | Control | UV-B | Control | UV-B |
| ***Cluster 3*** | | | | | | |
| At5g44990 | Glutathione S-transferase family protein | | 1,91 ± 3,3 | 286,94 ± 8,04 | 19,13 ± 8,04 | 248,33 ± 33,52 |
| At3g23230 | Integrase-type DNA-binding superfamily protein | *TRANSCRIPTIONAL REGULATOR OF DEFENSE RESPONSE 1 (TDR1)* | 1,78 ± 1,99 | 247,66 ± 1,25 | 0,73 ± 1,25 | 105,75 ± 12,15 |
| At3g06890 | Unknown protein | | 16,89 ± 1,99 | 42 ± 4,99 | 13,22 ± 4,99 | 80,24 ± 5,33 |
| At5g59820 | RHL41, ZAT12, C2H2-type zinc finger family protein | *RESPONSIVE TO HIGH LIGHT 41 (RHL41)* | 31,44 ± 7,03 | 774,79 ± 38,37 | 91 ± 38,37 | 750,32 ± 111,41 |
| At1g63440 | HMA5, heavy metal atpase 5 | *HEAVY METAL ATPASE 5 (HMA5)* | 149,13 ± 7,5 | 453,43 ± 37 | 209,99 ± 37 | 387,51 ± 42,36 |
| At5g35690 | Unknown protein | | 565,73 ± 43,86 | 1127,48 ± 44,16 | 557,9 ± 44,16 | 787,71 ± 47,48 |
| At1g62200 | Major facilitator superfamily protein | *PEPTIDE TRANSPORTER 6 (PTR6)* | 347,91 ± 13,62 | 666,41 ± 21,82 | 446,19 ± 21,82 | 585,81 ± 55,91 |
| At1g57560 | AtMYB50, MYB50, myb domain protein 50 | *MYB DOMAIN PROTEIN 5 (MYB5)* | 39,28 ± 5,19 | 91,68 ± 5,17 | 17,86 ± 5,17 | 150,75 ± 15,48 |
| At4g35000 | APX3, ascorbate peroxidase 3 | *ASCORBATE PEROXIDASE 3 (APX3)* | 1366,91 ± 112,92 | 2318,64 ± 225,57 | 2005,15 ± 225,57 | 2257,31 ± 231,81 |
| At3g18830 | ATPLT5, ATPMT5, PMT5, polyol/monosaccharide transporter 5 | *POLYOL/MONOSACCHARIDE TRANSPORTER 5 (PMT5)* | 571,74 ± 46,68 | 4925,19 ± 193,67 | 1149,16 ± 193,67 | 4588,43 ± 386,59 |
| At3g07890 | Ypt/Rab-GAP domain of gyp1p superfamily protein | | 403,85 ± 10,7 | 625,59 ± 8,54 | 459,33 ± 8,54 | 541,58 ± 35,76 |
| At5g42830 | HXXXD-type acyl-transferase family protein | | 45,75 ± 15,04 | 1617,85 ± 181,02 | 452,63 ± 181,02 | 3208,82 ± 940,63 |
| At1g48980 | 2-oxoglutarate (2OG) and Fe(II)-dependent oxygenase superfamily protein | | 5,55E-17 ± 7,55E-33 | 24,29 ± 1,24 | 0,72 ± 1,24 | 41,83 ± 3,67 |
| At4g09100 | RING/U-box superfamily protein | | 4,42 ± 0,17 | 15,72 ± 1,39 | 2,86 ± 1,39 | 27,15 ± 6,39 |
| At2g37430 | C2H2 and C2HC zinc fingers superfamily protein | | 5,82 ± 4,23 | 278,21 ± 2,2 | 2,21 ± 2,2 | 132,45 ± 12,08 |
| At2g15780 | Cupredoxin superfamily protein | | 5,55E-17 ± 7,55E-33 | 31,61 ± 7,55E-33 | 5,55E-17 ± 7,55E-33 | 10,9 ± 3,57 |
| At5g56960 | basic helix-loop-helix (bHLH) DNA-binding family protein | | 1,93 ± 1,81 | 269,83 ± 3,93 | 3,18 ± 3,93 | 194,12 ± 11,43 |
| At2g38870 | Serine protease inhibitor, potato inhibitor I-type family protein | | 224,49 ± 39,89 | 1210,98 ± 200,33 | 759,53 ± 200,33 | 1145,74 ± 69,09 |
| At3g13380 | BRL3, BRI1-like 3 | *BRI1-LIKE 3 (BRL3)* | 284,31 ± 20,71 | 808,58 ± 38 | 229,58 ± 38 | 1255,28 ± 82,11 |

**Supplementary Table S1. UV-B responsive genes regulated by TGA2/5/6 factors.** List of genes differentially regulated by the interaction between treatment and genotype.

| Locus | Gene model Description | Gene Symbol | Normalized expression | | | |
|---|---|---|---|---|---|---|
| | | | ***Wild type*** | | ***tga256*** | |
| | | | **Control** | **UV-B** | **Control** | **UV-B** |
| | ***Cluster 3*** | | | | | |
| At5g14980 | alpha/beta-Hydrolases superfamily protein | | 1 ± 0,87 | 20,58 ± 7,55E-33 | 5,55E-17 ± 7,55E-33 | 10 ± 1,48 |
| At1g09950 | RAS1, RESPONSE TO ABA AND SALT 1 | *RESPONSE TO ABA AND SALT 1 (RAS1)* | 1,6 ± 0,35 | 61,99 ± 2,2 | 2,21 ± 2,2 | 27,62 ± 6,54 |
| | ***Cluster 4*** | | | | | |
| At2g16586 | Unknown protein | | 0,53 ± 0,93 | 259 ± 4,73 | 3,31 ± 4,73 | 891,49 ± 158,97 |
| At1g74710 | ATICS1, EDS16, ICS1, SID2, ADC synthase superfamily protein | *ENHANCED DISEASE SUSCEPTIBILITY TO ERYSIPHE ORONTII 16 (EDS16)* | 159,99 ± 45,54 | 702,5 ± 43,45 | 282,66 ± 43,45 | 5355,5 ± 1389,29 |
| At1g32330 | ATHSFA1D, HSFA1D, heat shock transcription factor A1D | *HEAT SHOCK TRANSCRIPTION FACTOR A1D (HSFA1D)* | 223,21 ± 8,86 | 312,03 ± 20,65 | 187,37 ± 20,65 | 353,4 ± 9,94 |
| At1g71110 | Unknown protein | | 211,35 ± 14,2 | 262,03 ± 26,35 | 170,64 ± 26,35 | 331,97 ± 6,04 |
| At1g05990 | RHS1, EF hand calcium-binding protein family | | 9,05 ± 5,02 | 23,2 ± 3,51 | 4,13 ± 3,51 | 81,68 ± 2,29 |
| At4g10620 | P-loop containing nucleoside triphosphate hydrolases superfamily protein | | 218,76 ± 4,35 | 284,98 ± 12,76 | 133,74 ± 12,76 | 241,76 ± 15,38 |
| At3g22310 | ATRH9, PMH1, putative mitochondrial RNA helicase 1 | *PUTATIVE MITOCHONDRIAL RNA HELICASE 1 (PMH1)* | 787,58 ± 82,47 | 1925,07 ± 14,59 | 471,33 ± 14,59 | 1809,07 ± 309,47 |
| At3g24900 | AtRLP39, RLP39, receptor like protein 39 | *(RLP39)* | 8,46 ± 9,01 | 37,9 ± 3,67 | 12,17 ± 3,67 | 133,38 ± 14,35 |
| At2g40390 | Unknown protein | | 47,98 ± 2,65 | 52,58 ± 9,55 | 20,44 ± 9,55 | 59,8 ± 8,63 |
| At5g52730 | Copper transport protein family | | 0,53 ± 0,93 | 12,75 ± 0,54 | 0,32 ± 0,54 | 129,27 ± 58,72 |
| At2g21900 | WRKY DNA-binding protein 59 | *WRKY59* | 5,55E-17 ± 7,55E-33 | 2,07 ± 0,77 | 15,52 ± 0,77 | 61,13 ± 6,57 |
| At5g39770 | Restriction endonuclease, type II-like superfamily protein | | 0,49 ± 0,86 | 4,01 ± 0,55 | 2,83 ± 0,55 | 20,44 ± 3,9 |
| At3g17340 | ARM repeat superfamily protein | | 461,83 ± 30,88 | 476,23 ± 37,06 | 401,01 ± 37,06 | 617,04 ± 90,31 |
| At1g33720 | CYP76C6, cytochrome P450, family 76, subfamily C, polypeptide 6 | *CYP76C6* | 26,95 ± 11 | 79,32 ± 16,72 | 57,49 ± 16,72 | 549,69 ± 30,76 |
| At1g61990 | Mitochondrial transcription termination factor | | 244,83 ± 17,07 | 234,89 ± 15,34 | 169,84 ± 15,34 | 221,21 ± 4,63 |
| At1g32690 | Unknown protein | | 56,31 ± 12,46 | 33,7 ± 10,82 | 46,99 ± 10,82 | 77,49 ± 4,2 |
| At5g24190 | Lipase class 3-related protein | | 0,49 ± 0,86 | 1,38 ± 7,55E-33 | 5,55E-17 ± 7,55E-33 | 7,8 ± 1,67 |

**Supplementary Table S1. UV-B responsive genes regulated by TGA2/5/6 factors.** List of genes differentially regulated by the interaction between treatment and genotype.

| | | | Normalized expression | | | |
|---|---|---|---|---|---|---|
| | | | *Wild type* | | *tga256* | |
| Locus | Gene model Description | Gene Symbol | Control | UV-B | Control | UV-B |
| ***Cluster 4*** | | | | | | |
| At3g22400 | LOX5, PLAT/LH2 domain-containing lipoxygenase family protein | *(LOX5)* | 163,77 ± 18,4 | 283,54 ± 29,7 | 174,3 ± 29,7 | 810,19 ± 104,22 |
| At1g49590 | C2H2 and C2HC zinc fingers superfamily protein | | 294,55 ± 11,49 | 328,57 ± 8,87 | 243,26 ± 8,87 | 338,7 ± 11,74 |
| At1g70690 | HWI1, PDLP5, Receptor-like protein kinase-related family protein | *HOPW1-1-INDUCED GENE1 (HWI1)* | 114,59 ± 13,23 | 86,83 ± 13,91 | 157,38 ± 13,91 | 265,2 ± 56,09 |
| At5g01970 | Unknown protein | | 201,5 ± 18,79 | 174,4 ± 24,83 | 186,11 ± 24,83 | 284,66 ± 17,12 |
| At2g04450 | ATNUDT6, ATNUDX6, NUDT6, NUDX6, nudix hydrolase homolog 6 | *NUDIX HYDROLASE HOMOLOG 6 (NUDT6)* | 59,61 ± 41,87 | 205,76 ± 11,46 | 84,78 ± 11,46 | 1468,88 ± 57,77 |
| At4g09030 | AGP10, ATAGP10, arabinogalactan protein 10 | *ARABINOGALACTAN PROTEIN 1 (AGP1)* | 362,64 ± 47,27 | 851,63 ± 78,78 | 321,72 ± 78,78 | 2077,75 ± 723,84 |
| At5g38250 | Protein kinase family protein | | 3,33 ± 4,15 | 39,95 ± 6,29 | 9,35 ± 6,29 | 165,54 ± 6,67 |
| At1g66250 | O-Glycosyl hydrolases family 17 protein | | 194,5 ± 10,12 | 235,73 ± 13,16 | 175,34 ± 13,16 | 462,58 ± 69,75 |
| At5g17270 | Protein prenylyltransferase superfamily protein | | 295,26 ± 11,53 | 287,72 ± 2,63 | 209,1 ± 2,63 | 265,59 ± 9,2 |
| At3g13850 | LBD22, LOB domain-containing protein 22 | *(LBD22)* | 3,56 ± 3,08 | 2,71 ± 2,29 | 2,58 ± 2,29 | 17,03 ± 1,92 |
| At2g04430 | atnudt5, NUDT5, nudix hydrolase homolog 5 | *(NUDT5)* | 65,62 ± 12,08 | 90,87 ± 9,87 | 75,17 ± 9,87 | 220,64 ± 4,72 |
| At2g14620 | XTH10, xyloglucan endotransglucosylase/hydrolase 10 | *(XTH1)* | 23,06 ± 12,93 | 29,65 ± 5,78 | 19,67 ± 5,78 | 152,36 ± 66,17 |
| At1g14060 | GCK domain-containing protein | | 218,96 ± 23,56 | 593,5 ± 10,54 | 108,96 ± 10,54 | 446,64 ± 40,87 |
| At4g36800 | RCE1, RUB1 conjugating enzyme 1 | *RUB1 CONJUGATING ENZYME 1 (RCE1)* | 1619,21 ± 37,7 | 1660,28 ± 51,5 | 1625,11 ± 51,5 | 1923,81 ± 59,85 |
| At4g34390 | XLG2, extra-large GTP-binding protein 2 | *(XLG2)* | 399,13 ± 47,97 | 1182,6 ± 32,1 | 439,3 ± 32,1 | 2194,07 ± 236,25 |
| At4g13890 | EDA36, EDA37, SHM5, Pyridoxal phosphate (PLP)-dependent transferases superfamily protein | *EMBRYO SAC DEVELOPMENT ARREST 37 (EDA36)* | 20,46 ± 5,22 | 9,24 ± 10,52 | 39,63 ± 10,52 | 68,15 ± 22,47 |
| At3g24535 | Unknown protein | | 15,42 ± 2,99 | 11,85 ± 3,13 | 16,82 ± 3,13 | 25,59 ± 4,12 |
| At3g29170 | Eukaryotic protein of unknown function (DUF872) | | 116,19 ± 17,92 | 134,01 ± 6,81 | 95,66 ± 6,81 | 162,17 ± 4,91 |
| At5g52700 | Copper transport protein family | | 5,55E-17 ± 7,55E-33 | 5,55E-17 ± 1,09 | 0,65 ± 1,09 | 8,23 ± 2,16 |

**Supplementary Table S1. UV-B responsive genes regulated by TGA2/5/6 factors.** List of genes differentially regulated by the interaction between treatment and genotype.

| | | | Normalized expression | | | |
|---|---|---|---|---|---|---|
| | | | *Wild type* | | *tga256* | |
| Locus | Gene model Description | Gene Symbol | Control | UV-B | Control | UV-B |
| ***Cluster 4*** | | | | | | |
| At2g30750 | CYP71A12, cytochrome P450, family 71, subfamily A, polypeptide 12 | *CYP71A12* | 44,48 ± 18,22 | 1214,84 ± 262,01 | 662,86 ± 262,01 | 2575,63 ± 326,76 |
| At2g14120 | DRP3B, dynamin related protein | *DYNAMIN RELATED PROTEIN (DRP3B)* | 876,46 ± 13,25 | 906,78 ± 39,51 | 786,17 ± 39,51 | 990,65 ± 39,9 |
| At5g41550 | Disease resistance protein (TIR-NBS-LRR class) family | | 8,9 ± 2,12 | 41,99 ± 2,93 | 17,88 ± 2,93 | 113,14 ± 20,88 |
| At1g64880 | Ribosomal protein S5 family protein | | 989,75 ± 77,38 | 941,64 ± 10,97 | 547,09 ± 10,97 | 679,23 ± 26,24 |
| At2g07806 | Unknown protein | | 0,49 ± 0,42 | 0,65 ± 1,71 | 2,49 ± 1,71 | 14,05 ± 5,52 |
| At5g39960 | GTP binding;GTP binding | | 185,37 ± 23,52 | 209,4 ± 11,15 | 99,29 ± 11,15 | 173,24 ± 8,8 |
| At3g54960 | ATPDI1, ATPDIL1-3, PDI1, PDIL1-3, PDI-like 1-3 | *PDI-LIKE 1-3 (PDIL1-3)* | 874,81 ± 48,5 | 926,38 ± 110,01 | 794,37 ± 110,01 | 1290,68 ± 51,38 |
| At3g10090 | Nucleic acid-binding, OB-fold-like protein | | 287,82 ± 14,61 | 266,77 ± 17,16 | 203,18 ± 17,16 | 256,17 ± 15,79 |
| At1g21370 | Unknown protein | | 243,09 ± 17,01 | 410,49 ± 13,56 | 257,64 ± 13,56 | 690,72 ± 57,58 |
| At4g15020 | hAT transposon superfamily | | 329,08 ± 36,19 | 377,04 ± 18,94 | 288,71 ± 18,94 | 475,36 ± 21,79 |
| At5g61900 | BON, BON1, CPN1, Calcium-dependent phospholipid-binding Copine family protein | *BONZAI 1 (BON1)* | 382,23 ± 41,7 | 1097,79 ± 36,95 | 492,91 ± 36,95 | 2078,88 ± 41,79 |
| At2g17790 | VPS35A, ZIP3, VPS35 homolog A | *VPS35 HOMOLOG A (VPS35A)* | 622,3 ± 32,72 | 712,25 ± 24,68 | 585,48 ± 24,68 | 938,65 ± 61,83 |
| At3g29160 | AKIN11, ATKIN11, KIN11, SNRK1.2, SNF1 kinase homolog 11 | *SNF1 KINASE HOMOLOG 11 (KIN11)* | 533,5 ± 7,81 | 605,26 ± 11,44 | 518,1 ± 11,44 | 776,8 ± 17,26 |
| At1g66173 | other RNA | | 8,27 ± 3,17 | 12,05 ± 4,16 | 3,98 ± 4,16 | 26,47 ± 2,11 |
| At1g68990 | MGP3, male gametophyte defective 3 | *MALE GAMETOPHYTE DEFECTIVE 3 (MGP3)* | 492,55 ± 24,25 | 833,2 ± 26,56 | 324,95 ± 26,56 | 739,72 ± 79,08 |
| At3g14475 | snoRNA | | 17,22 ± 6,75 | 19,53 ± 2,27 | 28,03 ± 2,27 | 62,06 ± 9,02 |
| At2g36300 | Integral membrane Yip1 family protein | | 280,22 ± 13,26 | 431,77 ± 17,52 | 302,93 ± 17,52 | 685,96 ± 35,07 |
| At1g58225 | Unknown protein | | 5,97 ± 7,4 | 34,01 ± 8,72 | 21,41 ± 8,72 | 192,88 ± 30,72 |
| At3g48100 | ARR5, ATRR2, IBC6, RR5, response regulator 5 | *RR5* | 128,33 ± 21,73 | 126,94 ± 10,81 | 70,2 ± 10,81 | 172,07 ± 26,96 |
| At1g33730 | CYP76C5, cytochrome P450, family 76, subfamily C, polypeptide 5 | *CYP76C5* | 0,53 ± 0,93 | 8,63 ± 1,24 | 0,72 ± 1,24 | 136,86 ± 3,13 |
| At3g01316 | snoRNA | | 0,24 ± 0,41 | 0,31 ± 7,55E-33 | 5,55E-17 ± 7,55E-33 | 2,18 ± 0,56 |
| At1g67060 | Unknown protein | | 164,49 ± 5,19 | 305,6 ± 9,46 | 201,72 ± 9,46 | 523,39 ± 45,01 |
| At5g40720 | Domain of unknown function (DUF23) | | 156,78 ± 3,12 | 112,24 ± 39,5 | 216,75 ± 39,5 | 275,4 ± 16,56 |

**Supplementary Table S1. UV-B responsive genes regulated by TGA2/5/6 factors.** List of genes differentially regulated by the interaction between treatment and genotype.

| | | | Normalized expression | | | |
|---|---|---|---|---|---|---|
| | | | *Wild type* | | *tga256* | |
| Locus | Gene model Description | Gene Symbol | Control | UV-B | Control | UV-B |
| ***Cluster 4*** | | | | | | |
| At4g30990 | ARM repeat superfamily protein | | 1186,2 ± 122,46 | 1095,6 ± 12,55 | 630,69 ± 12,55 | 890,54 ± 73,95 |
| At3g57950 | Unknown protein | | 24,93 ± 5,24 | 25,79 ± 7,61 | 19,81 ± 7,61 | 80,07 ± 12,6 |
| At3g60470 | Plant protein of unknown function (DUF247) | | 0,53 ± 0,93 | 40,23 ± 6,16 | 14,01 ± 6,16 | 152,65 ± 42,72 |
| At4g35190 | Putative lysine decarboxylase family protein | *LONELY GUY 5 (LOG5)* | 9,81 ± 3,19 | 129,1 ± 9,95 | 14,97 ± 9,95 | 338,68 ± 68,53 |
| At4g04500 | CRK37, cysteine-rich RLK (RECEPTOR-like protein kinase) 37 | *CYSTEINE-RICH RLK (RECEPTOR-LIKE PROTEIN KINASE) 37 (CRK37)* | 6,2 ± 10,65 | 54,02 ± 11,36 | 23,18 ± 11,36 | 720,3 ± 357,2 |
| At1g55610 | BRL1, BRI1 like | *BRI1 LIKE (BRL1)* | 179,23 ± 15,44 | 145,88 ± 5,52 | 150,46 ± 5,52 | 228,06 ± 37,56 |
| At3g13100 | ATMRP7, MRP7, MRP7, multidrug resistance-associated protein 7 | *ATP-BINDING CASSETTE C7 (ABCC7)* | 64,58 ± 28,21 | 507,84 ± 16,17 | 58,13 ± 16,17 | 1505,49 ± 398,42 |
| At2g47810 | NF-YB5, nuclear factor Y, subunit B5 | *NUCLEAR FACTOR Y, SUBUNIT B5 (NF-YB5)* | 5,4 ± 1,52 | 2,77 ± 0,55 | 1,76 ± 0,55 | 10,83 ± 1,86 |
| At1g22885 | Unknown protein | | 130,2 ± 7,88 | 190,1 ± 3,1 | 126,77 ± 3,1 | 295,97 ± 26,41 |
| At3g46520 | ACT12, actin-12 | *ACTIN-12 (ACT12)* | 0,53 ± 0,93 | 10,9 ± 7,55E-33 | 5,55E-17 ± 7,55E-33 | 57,21 ± 31,67 |
| At3g20330 | PYRB, PYRIMIDINE B | *PYRIMIDINE B (PYRB)* | 695,84 ± 76,21 | 1078,53 ± 36,62 | 419,33 ± 36,62 | 1002,18 ± 78,24 |
| At1g74740 | ATCPK30, CDPK1A, CPK30, calcium-dependent protein kinase 30 | *CALCIUM-DEPENDENT PROTEIN KINASE 3 (CPK3)* | 317,64 ± 31,11 | 348,97 ± 9,13 | 256,58 ± 9,13 | 384,52 ± 24,26 |
| At1g17615 | Disease resistance protein (TIR-NBS class) | | 5,55E-17 ± 7,55E-33 | 5,22 ± 7,55E-33 | 5,55E-17 ± 7,55E-33 | 21,95 ± 7,01 |
| At4g00700 | C2 calcium/lipid-binding plant phosphoribosyltransferase family protein | | 221,36 ± 20,39 | 140,04 ± 34,93 | 193,28 ± 34,93 | 960,38 ± 235,21 |
| At1g77570 | Winged helix-turn-helix transcription repressor DNA-binding | | 19,04 ± 2,51 | 34,11 ± 1,99 | 7,82 ± 1,99 | 47,76 ± 9,61 |
| At1g70750 | Protein of unknown function, DUF593 | | 263,37 ± 2,28 | 290,78 ± 11,23 | 187,65 ± 11,23 | 313,29 ± 15,99 |
| At3g09700 | Chaperone DnaJ-domain superfamily protein | | 79,45 ± 5,77 | 77,44 ± 3,12 | 51,24 ± 3,12 | 76,28 ± 5,56 |
| At5g51630 | Disease resistance protein (TIR-NBS-LRR class) family | | 237,13 ± 18,7 | 429,37 ± 14,68 | 242,51 ± 14,68 | 675,89 ± 51,99 |
| At4g23160 | CRK8, cysteine-rich RLK (RECEPTOR-like protein kinase) 8 | *CYSTEINE-RICH RLK (RECEPTOR-LIKE PROTEIN KINASE) 8 (CRK8)* | 6,15 ± 4,05 | 67,1 ± 4,38 | 11,51 ± 4,38 | 326,08 ± 69,31 |

**Supplementary Table S1. UV-B responsive genes regulated by TGA2/5/6 factors.** List of genes differentially regulated by the interaction between treatment and genotype.

| Locus | Gene model Description | Gene Symbol | Normalized expression | | | |
|---|---|---|---|---|---|---|
| | | | *Wild type* | | *tga256* | |
| | | | Control | UV-B | Control | UV-B |
| | ***Cluster 4*** | | | | | |
| At1g70490 | ARFA1D, ATARFA1D, Ras-related small GTP-binding family protein | *(ARFA1D)* | 1945,71 ± 39,52 | 2300,33 ± 77,19 | 2117,13 ± 77,19 | 2892,36 ± 90,41 |
| At4g39030 | EDS5, SID1, MATE efflux family protein | *ENHANCED DISEASE SUSCEPTIBILITY 5 (EDS5)* | 200,39 ± 14,27 | 883,63 ± 33,76 | 183,56 ± 33,76 | 3176,33 ± 737,92 |
| At1g33390 | ATFAS4, FAS4, RNA helicase family protein | *FASCIATED STEM 4 (FAS4)* | 364,23 ± 23,23 | 445,06 ± 31,66 | 265,16 ± 31,66 | 515,67 ± 53,98 |
| At4g20370 | TSF, PEBP (phosphatidylethanolamine-binding protein) family protein | *TWIN SISTER OF FT (TSF)* | 0,53 ± 0,93 | 5,55E-17 ± 7,55E-33 | 5,55E-17 ± 7,55E-33 | 6,09 ± 2,37 |
| At1g61490 | S-locus lectin protein kinase family protein | | 55,92 ± 19,68 | 69,28 ± 19,6 | 62,38 ± 19,6 | 245,87 ± 76,84 |
| At5g65860 | ankyrin repeat family protein | | 223,57 ± 3,12 | 215,93 ± 1,27 | 139,56 ± 1,27 | 187,06 ± 14,17 |
| At1g73050 | Glucose-methanol-choline (GMC) oxidoreductase family protein | | 5,55E-17 ± 7,55E-33 | 5,55E-17 ± 7,55E-33 | 5,55E-17 ± 7,55E-33 | 3,08 ± 0,85 |
| At5g52720 | Copper transport protein family | | 4,02 ± 5,59 | 19,98 ± 11,89 | 14,62 ± 11,89 | 142,09 ± 37,52 |
| At5g01900 | ATWRKY62, WRKY62, WRKY DNA-binding protein 62 | *WRKY62* | 2,05 ± 2,47 | 6,48 ± 4,32 | 19,53 ± 4,32 | 78,72 ± 13,04 |
| At1g54170 | CID3, CTC-interacting domain 3 | *CTC-INTERACTING DOMAIN 3 (CID3)* | 459,28 ± 52,45 | 763,96 ± 29,79 | 523,13 ± 29,79 | 1132,04 ± 44,4 |
| At1g25580 | ANAC008, SOG1, NAC (No Apical Meristem) domain transcriptional regulator superfamily protein | *SUPPRESSOR OF GAMMA RADIATION 1 (SOG1)* | 218,04 ± 4,33 | 215,65 ± 32,32 | 214,44 ± 32,32 | 316,18 ± 25,73 |
| At4g26455 | WIP1, WPP domain interacting protein 1 | *WPP DOMAIN INTERACTING PROTEIN 1 (WIP1)* | 309 ± 10,79 | 403,56 ± 6,72 | 253,88 ± 6,72 | 446,81 ± 25,87 |
| At5g41730 | Protein kinase family protein | | 8,63 ± 3,4 | 13,87 ± 1,07 | 3,52 ± 1,07 | 24,15 ± 6,67 |
| At3g62420 | ATBZIP53, BZIP53, basic region/leucine zipper motif 53 | *BASIC REGION/LEUCINE ZIPPER MOTIF 53 (BZIP53)* | 891,11 ± 80,18 | 1031,45 ± 46,1 | 665,73 ± 46,1 | 1017,49 ± 31,87 |
| At1g77730 | Pleckstrin homology (PH) domain superfamily protein | | 5,13 ± 0,87 | 11,07 ± 1,09 | 1,05 ± 1,09 | 15,59 ± 3,27 |
| At4g29510 | ATPRMT11, ATPRMT1B, PRMT11, PRMT1B, arginine methyltransferase 11 | *ARGININE METHYLTRANSFERA SE 11 (PRMT11)* | 660,29 ± 34,78 | 982,12 ± 24,82 | 379,92 ± 24,82 | 824,83 ± 31,31 |
| At4g23200 | CRK12, cysteine-rich RLK (RECEPTOR-like protein kinase) 12 | *(CRK12)* | 37 ± 18,25 | 156,56 ± 7,73 | 11,3 ± 7,73 | 387,56 ± 32,44 |
| At2g20150 | Unknown protein | | 0,53 ± 0,93 | 6,95 ± 1,24 | 0,72 ± 1,24 | 21,05 ± 5,53 |

**Supplementary Table S1. UV-B responsive genes regulated by TGA2/5/6 factors.** List of genes differentially regulated by the interaction between treatment and genotype.

| | | | Normalized expression | | | |
|---|---|---|---|---|---|---|
| | | | *Wild type* | | *tga256* | |
| **Locus** | **Gene model Description** | **Gene Symbol** | **Control** | **UV-B** | **Control** | **UV-B** |
| ***Cluster 4*** | | | | | | |
| At3g60960 | Tetratricopeptide repeat (TPR)-like superfamily protein | | 120,4 ± 5,1 | 123,17 ± 8,65 | 70,82 ± 8,65 | 111,21 ± 7,47 |
| At2g33080 | AtRLP28, RLP28, receptor like protein 28 | *RECEPTOR LIKE PROTEIN 28 (RLP28)* | 1,09 ± 1,88 | 31,41 ± 4,72 | 10,41 ± 4,72 | 194,09 ± 55,77 |
| At4g26990 | Unknown protein | | 88,84 ± 8,36 | 179,98 ± 3,57 | 67,73 ± 3,57 | 439,4 ± 125,4 |
| At3g13130 | Unknown protein | | 1,52 ± 1,56 | 3,78 ± 4,76 | 4,3 ± 4,76 | 21,82 ± 3,05 |
| At2g40360 | Transducin/WD40 repeat-like superfamily protein | | 1783,59 ± 156,48 | 2104,75 ± 21,92 | 1040,49 ± 21,92 | 1643,6 ± 72,6 |
| At5g16480 | Phosphotyrosine protein phosphatases superfamily protein | *PLANT AND FUNGI ATYPICAL DUAL-SPECIﬁCITY PHOSPHATASE 5 (PFA-DSP5)* | 154,38 ± 30,08 | 181,24 ± 24,37 | 136,19 ± 24,37 | 385,93 ± 39,67 |
| At2g40740 | ATWRKY55, WRKY55, WRKY DNA-binding protein 55 | *WRKY55* | 2,78 ± 3,33 | 10,37 ± 3,28 | 6,5 ± 3,28 | 53,06 ± 14,04 |
| At3g01760 | Transmembrane amino acid transporter family protein | | 14,78 ± 1,28 | 4,51 ± 1,28 | 1,48 ± 1,28 | 18,72 ± 1,43 |
| At2g29110 | ATGLR2.8, GLR2.8, GLR2.8, glutamate receptor 2.8 | *GLUTAMATE RECEPTOR 2.8 (GLR2.8)* | 3,02 ± 1,57 | 19,11 ± 4,66 | 13,26 ± 4,66 | 71,13 ± 5,97 |
| At5g01700 | Protein phosphatase 2C family protein | | 41,3 ± 7,95 | 57,06 ± 1,35 | 22,93 ± 1,35 | 66,79 ± 5,54 |
| At1g12420 | ACR8, ACT domain repeat 8 | *ACT DOMAIN REPEAT 8 (ACR8)* | 120,69 ± 11,46 | 196,3 ± 22,82 | 127,09 ± 22,82 | 382,98 ± 52,99 |
| At5g63225 | Carbohydrate-binding X8 domain superfamily protein | | 5,55E-17 ± 7,55E-33 | 2,64 ± 1,87 | 1,09 ± 1,87 | 14,44 ± 4 |
| At3g18620 | DHHC-type zinc finger family protein | | 118,1 ± 4,57 | 138,46 ± 14,2 | 85,23 ± 14,2 | 159,29 ± 7,28 |
| At3g48630 | Unknown protein | | 2,83 ± 3,86 | 118,37 ± 3,51 | 8,71 ± 3,51 | 296,42 ± 37,39 |
| At3g04460 | APM4, ATPEX12, PEX12, peroxin-12 | *PEROXIN-12 (PEX12)* | 182,38 ± 5,19 | 269,71 ± 10,53 | 194,6 ± 10,53 | 409,4 ± 10,53 |
| At4g00840 | DHHC-type zinc finger family protein | | 197,02 ± 5,68 | 190,74 ± 30,58 | 211,67 ± 30,58 | 293,5 ± 10,43 |
| At1g01680 | ATPUB54, PUB54, plant U-box 54 | *PLANT U-BOX 54 (PUB54)* | 6,71 ± 11,63 | 44,43 ± 15,24 | 17,66 ± 15,24 | 485,82 ± 154,06 |
| At3g03670 | Peroxidase superfamily protein | | 1,26 ± 1,17 | 3,03 ± 6,52 | 44,43 ± 6,52 | 100,04 ± 17,45 |
| At5g55500 | ATXYLT, XYLT, beta-1,2-xylosyltransferase | *BETA-1,2-XYLOSYLTRANSFERASE (XYLT)* | 394,65 ± 29,12 | 358,01 ± 29,36 | 410,92 ± 29,36 | 493,64 ± 11,43 |
| At3g17690 | ATCNGC19, CNGC19, cyclic nucleotide gated channel 19 | *(CNGC19)* | 8,69 ± 8,84 | 225,95 ± 5,21 | 16,68 ± 5,21 | 566,43 ± 45,69 |

**Supplementary Table S1. UV-B responsive genes regulated by TGA2/5/6 factors.** List of genes differentially regulated by the interaction between treatment and genotype.

| | | | Normalized expression | | | |
|---|---|---|---|---|---|---|
| | | | *Wild type* | | *tga256* | |
| Locus | Gene model Description | Gene Symbol | Control | UV-B | Control | UV-B |
| ***Cluster 4*** | | | | | | |
| At2g26610 | Transducin family protein / WD-40 repeat family protein | | 5,55E-17 ± 7,55E-33 | 1,68 ± 1,09 | 1,11 ± 1,09 | 7,42 ± 1,4 |
| At5g39710 | EMB2745, Tetratricopeptide repeat (TPR)-like superfamily protein | *EMBRYO DEFECTIVE 2745 (EMB2745)* | 131,08 ± 8,46 | 158,5 ± 2,48 | 83,2 ± 2,48 | 133,32 ± 5,76 |
| At5g25540 | CID6, CTC-interacting domain 6 | *CTC-INTERACTING DOMAIN 6 (CID6)* | 528,43 ± 27,84 | 607,09 ± 44,33 | 538,09 ± 44,33 | 826,02 ± 54,58 |
| At4g11340 | Disease resistance protein (TIR-NBS-LRR class) family | | 5,55E-17 ± 7,55E-33 | 2,72 ± 0,54 | 0,32 ± 0,54 | 8,78 ± 2,95 |
| At1g21525 | pseudogene of unknown protein | | 8,5 ± 13,34 | 19,74 ± 4,97 | 8,34 ± 4,97 | 170,18 ± 36,56 |
| AtMg00120 | ORF143, hypothetical protein | *(ORF143)* | 0,26 ± 0,45 | 5,55E-17 ± 1,28 | 1,48 ± 1,28 | 5,26 ± 1,32 |
| At3g14590 | NTMC2T6.2, NTMC2TYPE6.2, Calcium-dependent lipid-binding (CaLB domain) family protein | *(NTMC2T6.2)* | 131,55 ± 17,2 | 157,3 ± 10,18 | 90,45 ± 10,18 | 188,71 ± 15,5 |
| At3g51320 | Pentatricopeptide repeat (PPR) superfamily protein | | 126,96 ± 2,98 | 122,11 ± 2,87 | 76,23 ± 2,87 | 107,17 ± 7,21 |
| At4g01560 | MEE49, Ribosomal RNA processing Brix domain protein | *MATERNAL EFFECT EMBRYO ARREST 49 (MEE49)* | 560,79 ± 23,16 | 953,8 ± 18,16 | 338,74 ± 18,16 | 764,4 ± 47,88 |
| At4g21840 | ATMSRB8, MSRB8, methionine sulfoxide reductase B8 | *METHIONINE SULFOXIDE REDUCTASE B8 (MSRB8)* | 12,77 ± 18,13 | 61,18 ± 14,91 | 30,08 ± 14,91 | 374,61 ± 76,37 |
| ***Cluster 5*** | | | | | | |
| At3g27960 | Tetratricopeptide repeat (TPR)-like superfamily protein | | 303,23 ± 36,64 | 103,45 ± 33,01 | 161,23 ± 33,01 | 93,01 ± 7,31 |
| At3g13420 | Unknown protein | | 50,27 ± 6,06 | 27,86 ± 5,37 | 32,25 ± 5,37 | 37,83 ± 1,27 |
| At3g09250 | Nuclear transport factor 2 (NTF2) family protein | | 1360,35 ± 32,99 | 868,7 ± 43,81 | 584,71 ± 43,81 | 501,65 ± 51,78 |
| At5g01330 | PDC3, pyruvate decarboxylase-3 | *PYRUVATE DECARBOXYLASE-3 (PDC3)* | 63,05 ± 8,38 | 5,78 ± 6,97 | 32,86 ± 6,97 | 13,72 ± 9,28 |
| At4g11780 | Unknown protein | *TON1 RECRUITING MOTIF 1 (TRM1)* | 40,13 ± 3,57 | 4,65 ± 2,18 | 18,25 ± 2,18 | 5,86 ± 4,93 |
| At2g22180 | hydroxyproline-rich glycoprotein family protein | | 0,9 ± 0,4 | 5,55E-17 ± 7,55E-33 | 5,55E-17 ± 7,55E-33 | 5,55E-17 ± 7,55E-33 |
| At1g62500 | Bifunctional inhibitor/lipid-transfer protein/seed storage 2S albumin superfamily protein | | 749,99 ± 78,48 | 231,67 ± 58,83 | 399,31 ± 58,83 | 213,77 ± 17,38 |

**Supplementary Table S1. UV-B responsive genes regulated by TGA2/5/6 factors.** List of genes differentially regulated by the interaction between treatment and genotype.

| | | | Normalized expression | | | |
|---|---|---|---|---|---|---|
| | | | *Wild type* | | *tga256* | |
| Locus | Gene model Description | Gene Symbol | Control | UV-B | Control | UV-B |
| ***Cluster 5*** | | | | | | |
| At5g23990 | ATFRO5, FRO5, ferric reduction oxidase 5 | *FERRIC REDUCTION OXIDASE 5 (FRO5)* | 125,13 ± 36,91 | 22,01 ± 7,44 | 29,72 ± 7,44 | 6,92 ± 2,2 |
| At1g53935 | Unknown protein | | 2,55 ± 1,22 | 5,55E-17 ± 7,55E-33 | 5,55E-17 ± 7,55E-33 | 5,55E-17 ± 7,55E-33 |
| At3g21340 | Leucine-rich repeat protein kinase family protein | | 31,04 ± 4,08 | 10,55 ± 0,66 | 6,02 ± 0,66 | 0,8 ± 1,29 |
| At2g37840 | Protein kinase superfamily protein | | 418,48 ± 4,35 | 286,94 ± 11,64 | 476,07 ± 11,64 | 399,27 ± 18,36 |
| At3g27884 | other RNA | | 36,9 ± 5,77 | 4,82 ± 2,48 | 9,82 ± 2,48 | 6,57 ± 1,38 |
| At3g24310 | ATMYB71, MYB305, myb domain protein 305 | *MYB35* | 11,08 ± 3,12 | 2,75 ± 1,86 | 1,08 ± 1,86 | 2,2 ± 2,59 |
| At3g32030 | Terpenoid cyclases/Protein prenyltransferases superfamily protein | | 41,48 ± 13,1 | 5,5 ± 1,17 | 1,38 ± 1,17 | 4,34 ± 1,14 |
| At4g04745 | Unknown protein | | 64,42 ± 15,24 | 4,35 ± 3,44 | 24,92 ± 3,44 | 2,07 ± 3,37 |
| At2g31970 | ATRAD50, RAD50, DNA repair-recombination protein (RAD50) | *(RAD5)* | 453,02 ± 5,38 | 381,07 ± 16,99 | 361,77 ± 16,99 | 375,47 ± 25,6 |
| At4g12330 | CYP706A7, cytochrome P450, family 706, subfamily A, polypeptide 7 | *CYP76A7* | 90,01 ± 26,56 | 32,77 ± 2,98 | 7,99 ± 2,98 | 9,2 ± 2,78 |
| At5g60660 | PIP2;4, PIP2F, plasma membrane intrinsic protein 2;4 | *PLASMA MEMBRANE INTRINSIC PROTEIN 2;4 (PIP2;4)* | 279,17 ± 6,32 | 58,81 ± 29,48 | 152,56 ± 29,48 | 70,72 ± 8,25 |
| At1g03010 | Phototropic-responsive NPH3 family protein | | 48,37 ± 13,85 | 9,92 ± 2,95 | 12,94 ± 2,95 | 12,14 ± 1,79 |
| At4g33880 | RSL2, ROOT HAIR DEFECTIVE 6-LIKE 2 | *ROOT HAIR DEFECTIVE 6-LIKE 2 (RSL2)* | 62,73 ± 13,37 | 2,62 ± 2,9 | 12,93 ± 2,9 | 1,27 ± 1,18 |
| At1g61950 | CPK19, calcium-dependent protein kinase 19 | *(CPK19)* | 23,17 ± 2,09 | 3,08 ± 3,3 | 4,8 ± 3,3 | 5,55E-17 ± 7,55E-33 |
| At1g12070 | Immunoglobulin E-set superfamily protein | | 7,24 ± 0,75 | 0,97 ± 0,83 | 3,56 ± 0,83 | 0,87 ± 0,71 |
| At5g65440 | Unknown protein | | 721,81 ± 33,71 | 434,23 ± 17,87 | 717,66 ± 17,87 | 527,93 ± 16,41 |
| At5g18460 | Protein of Unknown Function (DUF239) | | 394,4 ± 36,93 | 96,06 ± 17,41 | 298,33 ± 17,41 | 127,57 ± 3,57 |
| At1g74660 | MIF1, mini zinc finger 1 | *MINI ZINC FINGER 1 (MIF1)* | 59,68 ± 6,24 | 14,58 ± 12,47 | 46,67 ± 12,47 | 24,82 ± 2,22 |
| At4g20210 | Terpenoid cyclases/Protein prenyltransferases superfamily protein | | 15,08 ± 3,99 | 1,06 ± 2,2 | 4,87 ± 2,2 | 1,66 ± 2,7 |
| At4g26220 | S-adenosyl-L-methionine-dependent methyltransferases superfamily protein | | 137,34 ± 18,75 | 78,76 ± 10,38 | 39,98 ± 10,38 | 52,98 ± 13,11 |
| At5g57650 | eukaryotic translation initiation factor-related | | 1,14 ± 0,44 | 5,55E-17 ± 7,55E-33 | 5,55E-17 ± 7,55E-33 | 5,55E-17 ± 7,55E-33 |

**Supplementary Table S1. UV-B responsive genes regulated by TGA2/5/6 factors.** List of genes differentially regulated by the interaction between treatment and genotype.

| Locus | Gene model Description | Gene Symbol | Normalized expression | | | |
|---|---|---|---|---|---|---|
| | | | *Wild type* | | *tga256* | |
| | | | Control | UV-B | Control | UV-B |
| | ***Cluster 5*** | | | | | |
| At5g28913 | SADHU4-1, transposable element gene | *SADHU NON-CODING RETROTRANSPOSON 4-1 (SADHU4-1)* | 97,15 ± 10,92 | 32,27 ± 9,03 | 106,93 ± 9,03 | 62,12 ± 7,03 |
| At5g06200 | Uncharacterised protein family (UPF0497) | *CASPARIAN STRIP MEMBRANE DOMAIN PROTEIN 4 (CASP4)* | 42,09 ± 3,58 | 5,24 ± 3,51 | 9,24 ± 3,51 | 4,34 ± 3,75 |
| At5g65800 | ACS5, ATACS5, CIN5, ACC synthase 5 | *(ACS5)* | 25,73 ± 2,55 | 0,71 ± 0,44 | 15,56 ± 0,44 | 5,55E-17 ± 7,55E-33 |
| At3g01260 | Galactose mutarotase-like superfamily protein | | 174,06 ± 39,49 | 15,55 ± 11,42 | 45,42 ± 11,42 | 5,15 ± 5,07 |
| At1g72510 | Protein of unknown function (DUF1677) | | 654,3 ± 59,69 | 194,41 ± 47,06 | 459,62 ± 47,06 | 228 ± 13,16 |
| At5g56540 | AGP14, ATAGP14, arabinogalactan protein 14 | *AGP14)* | 204,84 ± 34,73 | 57,06 ± 2,41 | 84,71 ± 2,41 | 45,64 ± 7,68 |
| At2g17440 | PIRL5, plant intracellular ras group-related LRR 5 | *PLANT INTRACELLULAR RAS GROUP-RELATED LRR 5 (PIRL5)* | 614,1 ± 59 | 425,92 ± 24,65 | 549,1 ± 24,65 | 551,91 ± 53,93 |
| At1g77530 | O-methyltransferase family protein | | 68,98 ± 6,96 | 16,34 ± 1,69 | 7,49 ± 1,69 | 2,52 ± 2,36 |
| At5g24100 | Leucine-rich repeat protein kinase family protein | | 80,44 ± 7,02 | 4,79 ± 8,73 | 24,95 ± 8,73 | 4,77 ± 1,63 |
| At1g08590 | Leucine-rich receptor-like protein kinase family protein | | 388,6 ± 17,1 | 106,63 ± 25,09 | 236,27 ± 25,09 | 110,76 ± 8,27 |
| At5g67620 | Unknown protein | | 30,89 ± 3,52 | 6,59 ± 0,97 | 10,28 ± 0,97 | 4,34 ± 3,75 |
| At3g12977 | NAC (No Apical Meristem) domain transcriptional regulator superfamily protein | | 60,62 ± 7,43 | 9,33 ± 3,02 | 19,18 ± 3,02 | 5,08 ± 4,58 |
| At2g47530 | Pollen Ole e 1 allergen and extensin family protein | | 35,17 ± 2,18 | 10,03 ± 0,1 | 4,59 ± 0,1 | 3,03 ± 1,73 |
| At2g31310 | LBD14, LOB domain-containing protein 14 | *(LBD14)* | 7,93 ± 1,31 | 1,66 ± 1,24 | 0,72 ± 1,24 | 0,8 ± 1,29 |
| At2g42850 | CYP718, cytochrome P450, family 718 | *CYP718* | 64,15 ± 5,78 | 7,15 ± 3,26 | 15,39 ± 3,26 | 2,52 ± 2,36 |
| At1g05530 | UGT2, UGT75B2, UDP-glucosyl transferase 75B2 | *UGT75B2* | 33,91 ± 3,57 | 6,23 ± 2,84 | 13,51 ± 2,84 | 4,3 ± 2,41 |
| At5g65390 | AGP7, arabinogalactan protein 7 | *ARABINOGALACTAN PROTEIN 7 (AGP7)* | 746,55 ± 89,83 | 49 ± 27,82 | 504,14 ± 27,82 | 64,54 ± 16,4 |
| At3g56470 | F-box family protein | | 112,34 ± 3,73 | 64,51 ± 8,95 | 106,15 ± 8,95 | 90,51 ± 4,03 |
| At4g25250 | Plant invertase/pectin methylesterase inhibitor superfamily protein | | 49,04 ± 10,57 | 0,71 ± 2,79 | 5,73 ± 2,79 | 5,55E-17 ± 7,55E-33 |

**Supplementary Table S1. UV-B responsive genes regulated by TGA2/5/6 factors.** List of genes differentially regulated by the interaction between treatment and genotype.

| Locus | Gene model Description | Gene Symbol | Normalized expression | | | |
|---|---|---|---|---|---|---|
| | | | *Wild type* | | *tga256* | |
| | | | Control | UV-B | Control | UV-B |
| | ***Cluster 5*** | | | | | |
| At1g06710 | Tetratricopeptide repeat (TPR)-like superfamily protein | | 149,42 ± 12,88 | 114,09 ± 17,44 | 79,09 ± 17,44 | 103,35 ± 8,04 |
| At5g52790 | CBS domain-containing protein with a domain of unknown function (DUF21) | | 217,13 ± 45,02 | 51,13 ± 18,95 | 33,73 ± 18,95 | 46,96 ± 5,29 |
| At1g21740 | Protein of unknown function (DUF630 and DUF632) | | 175,48 ± 6,76 | 85,73 ± 4,73 | 144,38 ± 4,73 | 130,77 ± 24,85 |
| At4g20325 | Unknown | | 120,46 ± 13,98 | 88,27 ± 1,73 | 72,71 ± 1,73 | 84,66 ± 11,3 |
| At3g22240 | Unknown protein | | 469,92 ± 41,17 | 175,89 ± 60,85 | 279,31 ± 60,85 | 196,9 ± 19,77 |
| At3g07490 | AGD11, ARF-GAP domain 11 | *ARF-GAP DOMAIN 11 (AGD11)* | 20,73 ± 4,78 | 1,66 ± 1,86 | 1,08 ± 1,86 | 0,87 ± 1,42 |
| At1g74830 | Protein of unknown function, DUF593 | | 27,98 ± 9,56 | 3,7 ± 2,2 | 4,87 ± 2,2 | 4,4 ± 1,56 |
| At1g62440 | LRX2, leucine-rich repeat/extensin 2 | *LEUCINE-RICH REPEAT/EXTENSIN 2 (LRX2)* | 208,14 ± 23,05 | 34,32 ± 40,91 | 124,15 ± 40,91 | 54,23 ± 6,3 |
| At5g46890 | Bifunctional inhibitor/lipid-transfer protein/seed storage 2S albumin superfamily protein | | 457,01 ± 74,45 | 25,28 ± 41,89 | 132,56 ± 41,89 | 25,82 ± 7,66 |
| At4g37160 | sks15, SKU5 similar 15 | *SKU5 SIMILAR 15 (sks15)* | 160,15 ± 12,57 | 31,9 ± 6,61 | 76,28 ± 6,61 | 28,56 ± 6,59 |
| At1g79320 | AtMC6, MC6, metacaspase 6 | *METACASPASE 6 (MC6)* | 43,3 ± 2,87 | 10,3 ± 4,73 | 14,96 ± 4,73 | 9,94 ± 3,91 |
| At3g50710 | F-box/RNI-like/FBD-like domains-containing protein | | 6,33 ± 1,59 | 0,31 ± 7,55E-33 | 5,55E-17 ± 7,55E-33 | 0,42 ± 0,68 |
| At3g18170 | Glycosyltransferase family 61 protein | | 128,41 ± 13,73 | 20,19 ± 17,73 | 84,81 ± 17,73 | 29,52 ± 6,83 |
| At2g31085 | CLE6, CLAVATA3/ESR-RELATED 6 | *CLAVATA3/ESR-RELATED 6 (CLE6)* | 21,43 ± 4,43 | 3,92 ± 1,21 | 3,49 ± 1,21 | 5,55E-17 ± 7,55E-33 |
| At5g26080 | proline-rich family protein | | 38,47 ± 1,01 | 1,43 ± 6,56 | 12,75 ± 6,56 | 2,5 ± 2,25 |
| At3g59680 | Unknown protein | | 86,31 ± 7,23 | 20,85 ± 7,69 | 54,93 ± 7,69 | 25,27 ± 1,93 |
| At4g20230 | Terpenoid cyclases/Protein prenyltransferases superfamily protein | | 52 ± 14,76 | 5,91 ± 3,09 | 7,48 ± 3,09 | 5,55E-17 ± 7,55E-33 |
| At3g45700 | Major facilitator superfamily protein | | 93,07 ± 22,59 | 7,76 ± 11,8 | 28,75 ± 11,8 | 3,92 ± 1,07 |
| At4g12080 | AHL1, ATAHL1, AT-hook motif nuclear-localized protein 1 | *AT-HOOK MOTIF NUCLEAR-LOCALIZED PROTEIN 1 (AHL1)* | 284,36 ± 29,87 | 90,59 ± 26,9 | 254,86 ± 26,9 | 152,03 ± 16,33 |
| At4g11320 | Papain family cysteine protease | | 6658,86 ± 835,21 | 1688,22 ± 290,94 | 746,41 ± 290,94 | 637,71 ± 116,17 |
| At5g36270 | pseudogene of dehydroascorbate reductase | | 18,38 ± 4,08 | 2,5 ± 1,64 | 0,97 ± 1,64 | 0,83 ± 1,35 |
| At1g06120 | Fatty acid desaturase family protein | | 29,83 ± 5,8 | 2,61 ± 4,82 | 10,91 ± 4,82 | 2,16 ± 1,91 |

**Supplementary Table S1. UV-B responsive genes regulated by TGA2/5/6 factors.** List of genes differentially regulated by the interaction between treatment and genotype.

| | | | Normalized expression | | | |
|---|---|---|---|---|---|---|
| | | | *Wild type* | | *tga256* | |
| Locus | Gene model Description | Gene Symbol | Control | UV-B | Control | UV-B |
| ***Cluster 5*** | | | | | | |
| At1g61050 | alpha 1,4-glycosyltransferase family protein | | 70,17 ± 4,82 | 13,83 ± 7,96 | 19,03 ± 7,96 | 3,47 ± 1,14 |
| At2g20520 | FLA6, FASCICLIN-like arabinogalactan 6 | *FASCICLIN-LIKE ARABINOGALACTAN 6 (FLA6)* | 87,46 ± 20,84 | 3,05 ± 4,36 | 48,28 ± 4,36 | 6,5 ± 1,66 |
| At1g10130 | ATECA3, ECA3, endoplasmic reticulum-type calcium-transporting ATPase 3 | *ENDOPLASMIC RETICULUM-TYPE CALCIUM-TRANSPORTING ATPASE 3 (ECA3)* | 795,7 ± 17,56 | 566,45 ± 20,83 | 915,03 ± 20,83 | 765,64 ± 14,43 |
| At5g65170 | VQ motif-containing protein | | 82,01 ± 13,95 | 57,21 ± 7,3 | 30,31 ± 7,3 | 51,12 ± 7,92 |
| At3g29780 | RALFL27, ralf-like 27 | *RALF-LIKE 27 (RALFL27)* | 54,72 ± 5,46 | 6,85 ± 3,8 | 7,95 ± 3,8 | 1,31 ± 1,2 |
| At4g29180 | RHS16, root hair specific 16 | *ROOT HAIR SPECIFIC 16 (RHS16)* | 154,86 ± 12,58 | 25,59 ± 5,77 | 23,59 ± 5,77 | 20,91 ± 3,15 |
| At5g65470 | O-fucosyltransferase family protein | | 599,2 ± 25,02 | 388,22 ± 36,99 | 465,71 ± 36,99 | 405,72 ± 33,33 |
| At1g11670 | MATE efflux family protein | | 391,83 ± 15,5 | 243,84 ± 12,91 | 248,3 ± 12,91 | 234,07 ± 30,05 |
| At2g31540 | GDSL-like Lipase/Acylhydrolase superfamily protein | | 13,02 ± 6,05 | 0,63 ± 7,55E-33 | 5,55E-17 ± 7,55E-33 | 0,83 ± 1,35 |
| At1g73020 | Unknown protein | | 195,64 ± 33,27 | 87,17 ± 18,22 | 168,17 ± 18,22 | 161,85 ± 12,67 |
| At5g50790 | Nodulin MtN3 family protein | *(SWEET1)* | 54,94 ± 11,77 | 14,78 ± 2,5 | 1,46 ± 2,5 | 2,15 ± 3,54 |
| At5g62310 | IRE, AGC (cAMP-dependent, cGMP-dependent and protein kinase C) kinase family protein | *INCOMPLETE ROOT HAIR ELONGATION (IRE)* | 51,21 ± 4,95 | 12,75 ± 3,35 | 24,12 ± 3,35 | 19,2 ± 4,76 |
| At5g54260 | ATMRE11, MRE11, DNA repair and meiosis protein (Mre11) | *MEIOTIC RECOMBINATION 11 (MRE11)* | 146,58 ± 16,93 | 67,03 ± 11,74 | 112,75 ± 11,74 | 95,88 ± 9,62 |
| At3g50640 | Unknown protein | | 93,17 ± 19,02 | 14,54 ± 2,57 | 36,63 ± 2,57 | 7,47 ± 2,24 |
| At3g32040 | Terpenoid synthases superfamily protein | | 30,95 ± 7,92 | 4,71 ± 1,11 | 3,52 ± 1,11 | 1,72 ± 1,41 |
| At1g55290 | 2-oxoglutarate (2OG) and Fe(II)-dependent oxygenase superfamily protein | | 21,47 ± 3,81 | 2,69 ± 1,84 | 3,59 ± 1,84 | 2,52 ± 2,36 |
| At5g24313 | Unknown protein | | 35,07 ± 5,74 | 2,48 ± 4,67 | 10,13 ± 4,67 | 5,55E-17 ± 7,55E-33 |
| At2g28990 | Leucine-rich repeat protein kinase protein | | 59,25 ± 4,59 | 20,77 ± 7,07 | 20,39 ± 7,07 | 11,72 ± 1,49 |
| At1g66950 | ATPDR11, PDR11, pleiotropic drug resistance 11 | *ATP-BINDING CASSETTE G39 (ABCG39)* | 17,34 ± 3,7 | 4,07 ± 4,21 | 15,33 ± 4,21 | 13,47 ± 0,99 |

**Supplementary Table S1. UV-B responsive genes regulated by TGA2/5/6 factors.** List of genes differentially regulated by the interaction between treatment and genotype.

| Locus | Gene model Description | Gene Symbol | Normalized expression | | | |
|---|---|---|---|---|---|---|
| | | | *Wild type* | | *tga256* | |
| | | | Control | UV-B | Control | UV-B |
| | ***Cluster 5*** | | | | | |
| At1g14220 | Ribonuclease T2 family protein | | 33,6 ± 6,15 | 2,34 ± 2,76 | 2,58 ± 2,76 | 5,55E-17 ± 7,55E-33 |
| At4g22230 | Arabidopsis defensin-like protein | | 23,05 ± 3,37 | 4,44 ± 3,73 | 6,19 ± 3,73 | 0,87 ± 1,42 |
| At2g01520 | MLP328, MLP-like protein 328 | *MLP-LIKE PROTEIN 328 (MLP328)* | 1971,1 ± 295,21 | 434,64 ± 21,02 | 19,24 ± 21,02 | 19,91 ± 15,15 |
| At3g30875 | pseudogene, putative multidrug resistance protein | | 96,28 ± 17,82 | 16,66 ± 3,95 | 27 ± 3,95 | 7,37 ± 1,36 |
| At1g64210 | Leucine-rich repeat protein kinase family protein | | 34,86 ± 2,58 | 17,48 ± 5,25 | 7,13 ± 5,25 | 8,27 ± 1,69 |
| At5g25810 | tny, Integrase-type DNA-binding superfamily protein | *TINY (tny)* | 113,16 ± 4,35 | 14,34 ± 2,64 | 65,18 ± 2,64 | 9,97 ± 2,69 |
| At4g22666 | Bifunctional inhibitor/lipid-transfer protein/seed storage 2S albumin superfamily protein | | 172,8 ± 21,15 | 10,07 ± 4,04 | 95,61 ± 4,04 | 10,24 ± 5,83 |
| At3g25790 | myb-like transcription factor family protein | | 77,67 ± 4,77 | 17,86 ± 4,73 | 11,12 ± 4,73 | 3,33 ± 2,86 |
| At3g09240 | Protein kinase protein with tetratricopeptide repeat domain | | 41,13 ± 11,05 | 6,41 ± 1,94 | 12,98 ± 1,94 | 6,95 ± 1,02 |
| At1g23160 | Auxin-responsive GH3 family protein | | 14,36 ± 3,12 | 3,41 ± 2,2 | 2,21 ± 2,2 | 0,83 ± 1,35 |
| At4g20240 | CYP71A27, cytochrome P450, family 71, subfamily A, polypeptide 27 | *CYP71A27* | 119,87 ± 26,5 | 24,3 ± 3,74 | 33,84 ± 3,74 | 7,5 ± 4,09 |
| At2g14830 | Regulator of Vps4 activity in the MVB pathway protein | | 76,6 ± 5,22 | 44,78 ± 10,28 | 89,95 ± 10,28 | 76,09 ± 0,79 |
| At1g69810 | ATWRKY36, WRKY36, WRKY DNA-binding protein 36 | *WRKY36* | 176,86 ± 19,73 | 84,1 ± 13,44 | 148,65 ± 13,44 | 126,37 ± 4,84 |
| At1g50630 | Protein of unknown function (DUF3537) | | 219,34 ± 5,83 | 113,07 ± 27,11 | 134,09 ± 27,11 | 118,85 ± 2,89 |
| At5g57760 | Unknown protein | | 208,44 ± 37,02 | 14,03 ± 9,11 | 95,83 ± 9,11 | 11,33 ± 0,33 |
| At1g49240 | ACT8, actin 8 | *ACTIN 8 (ACT8)* | 7051,71 ± 116,05 | 2736,79 ± 768,55 | 7977,95 ± 768,55 | 4171,74 ± 238,79 |
| At1g31772 | Defensin-like (DEFL) family protein | | 1,83 ± 0,8 | 5,55E-17 ± 7,55E-33 | 5,55E-17 ± 7,55E-33 | 5,55E-17 ± 7,55E-33 |
| At2g38320 | TBL34, TRICHOME BIREFRINGENCE-LIKE 34 | *TRICHOME BIREFRINGENCE-LIKE 34 (TBL34)* | 64,72 ± 6,14 | 24,84 ± 9,05 | 74,71 ± 9,05 | 55,35 ± 8,29 |
| At5g03530 | ATRAB, ATRAB ALPHA, ATRAB18B, ATRABC2A, RABC2A, RAB GTPase homolog C2A | *RAB GTPASE HOMOLOG C2A (RABC2A)* | 203,09 ± 1,39 | 117,76 ± 4,66 | 130,46 ± 4,66 | 115,38 ± 8,56 |
| At2g45890 | ATROPGEF4, RHS11, ROPGEF4, RHO guanyl-nucleotide exchange factor 4 | *RHO GUANYL-NUCLEOTIDE EXCHANGE FACTOR 4 (ROPGEF4)* | 83,56 ± 3,22 | 12,21 ± 5,45 | 40,36 ± 5,45 | 18,59 ± 4,93 |

Normalized expression

**Supplementary Table S1. UV-B responsive genes regulated by TGA2/5/6 factors.** List of genes differentially regulated by the interaction between treatment and genotype.

| Locus | Gene model Description | Gene Symbol | *Wild type* Control | *Wild type* UV-B | *tga256* Control | *tga256* UV-B |
|---|---|---|---|---|---|---|
| | ***Cluster 5*** | | | | | |
| At4g14130 | XTH15, XTR7, xyloglucan endotransglucosylase/hydrolase 15 | *XYLOGLUCAN ENDOTRANSGLUCOSYLASE/HYDROLASE 15 (XTH15)* | 1243,13 ± 369,7 | 86,35 ± 23,28 | 249,44 ± 23,28 | 62,64 ± 12,11 |
| At3g50190 | Plant protein of unknown function (DUF247) | | 31,35 ± 3,05 | 16,21 ± 3,43 | 24,27 ± 3,43 | 27,07 ± 4,44 |
| At1g02575 | Unknown protein | | 8,93 ± 1,7 | 0,36 ± 1,09 | 0,65 ± 1,09 | 5,55E-17 ± 7,55E-33 |
| At1g64480 | CBL8, calcineurin B-like protein 8 | *CALCINEURIN B-LIKE PROTEIN 8 (CBL8)* | 13,9 ± 7,22 | 0,36 ± 1,09 | 0,65 ± 1,09 | 5,55E-17 ± 7,55E-33 |
| At5g26740 | Protein of unknown function (DUF300) | | 569,98 ± 40,41 | 351,77 ± 6,08 | 510,59 ± 6,08 | 380,76 ± 3,94 |
| At1g51870 | protein kinase family protein | | 27,25 ± 11,83 | 2,39 ± 1,05 | 4,58 ± 1,05 | 0,4 ± 0,65 |
| At1g70860 | Polyketide cyclase/dehydrase and lipid transport superfamily protein | | 12,9 ± 1,05 | 3,4 ± 0,77 | 2,5 ± 0,77 | 0,8 ± 1,29 |
| At3g61270 | Arabidopsis thaliana protein of unknown function (DUF821) | | 123,15 ± 16,33 | 25,69 ± 8,57 | 45,49 ± 8,57 | 20,75 ± 4,22 |
| At4g00651 | Unknown | | 3,03 ± 0,61 | 0,71 ± 7,55E-33 | 5,55E-17 ± 7,55E-33 | 5,55E-17 ± 7,55E-33 |
| At5g58360 | ATOFP3, OFP3, ovate family protein 3 | *OVATE FAMILY PROTEIN 3 (OFP3)* | 20,18 ± 1,66 | 1,07 ± 1,87 | 1,09 ± 1,87 | 5,55E-17 ± 7,55E-33 |
| At3g10710 | RHS12, root hair specific 12 | *ROOT HAIR SPECIFIC 12 (RHS12)* | 155,22 ± 10,57 | 14,76 ± 7,88 | 55,94 ± 7,88 | 14,89 ± 4,22 |
| At5g52990 | SNARE-like superfamily protein | | 100,66 ± 8,99 | 32,37 ± 14,72 | 65,79 ± 14,72 | 41,32 ± 4,02 |
| At1g58037 | Cysteine/Histidine-rich C1 domain family protein | | 14,58 ± 6,11 | 1,43 ± 0,68 | 1,43 ± 0,68 | 5,55E-17 ± 7,55E-33 |
| At3g20880 | WIP4, WIP domain protein 4 | *WIP DOMAIN PROTEIN 4 (WIP4)* | 2,09 ± 0,82 | 5,55E-17 ± 7,55E-33 | 5,55E-17 ± 7,55E-33 | 5,55E-17 ± 7,55E-33 |
| At3g24020 | Disease resistance-responsive (dirigent-like protein) family protein | | 229,36 ± 19,46 | 70,89 ± 18,39 | 166,04 ± 18,39 | 94,14 ± 16,31 |
| At2g45830 | DTA2, downstream target of AGL15 2 | *DOWNSTREAM TARGET OF AGL15 2 (DTA2)* | 155,35 ± 5,25 | 55,68 ± 2,93 | 85,3 ± 2,93 | 68,85 ± 18,94 |
| At1g11655 | Unknown protein | | 17,74 ± 3,6 | 5,55E-17 ± 2,96 | 6,59 ± 2,96 | 5,55E-17 ± 7,55E-33 |
| At5g42690 | Protein of unknown function, DUF547 | | 83,08 ± 2,37 | 29,66 ± 8,13 | 42,51 ± 8,13 | 20,44 ± 0,61 |
| At1g23030 | ARM repeat superfamily protein | | 535,01 ± 41,64 | 214,36 ± 24,79 | 336,57 ± 24,79 | 221,36 ± 24,09 |
| At2g16980 | Major facilitator superfamily protein | | 101,62 ± 8,29 | 12,07 ± 4,37 | 56,39 ± 4,37 | 10,54 ± 4,66 |
| At3g23690 | basic helix-loop-helix (bHLH) DNA-binding superfamily protein | | 570,61 ± 32,04 | 327,7 ± 9,59 | 541,7 ± 9,59 | 381,91 ± 21,68 |

**Supplementary Table S1. UV-B responsive genes regulated by TGA2/5/6 factors.** List of genes differentially regulated by the interaction between treatment and genotype.

| Locus | Gene model Description | Gene Symbol | Normalized expression | | | |
|---|---|---|---|---|---|---|
| | | | *Wild type* | | *tga256* | |
| | | | Control | UV-B | Control | UV-B |
| ***Cluster 5*** | | | | | | |
| At5g54790 | Unknown protein | | 21,07 ± 3,76 | 1,72 ± 3,3 | 2,94 ± 3,3 | 1,27 ± 1,18 |
| At1g69560 | ATMYB105, LOF2, MYB105, myb domain protein 105 | *MYB DOMAIN PROTEIN 15 (MYB15)* | 1,6 ± 0,35 | 5,55E-17 ± 7,55E-33 | 5,55E-17 ± 7,55E-33 | 0,4 ± 0,65 |
| At2g35585 | Unknown protein | | 31,92 ± 7,39 | 3,09 ± 3,79 | 14,67 ± 3,79 | 3,96 ± 3,33 |
| At2g18980 | Peroxidase superfamily protein | | 387,87 ± 31,97 | 73,27 ± 1,81 | 173,65 ± 1,81 | 48,16 ± 3,81 |
| At5g21130 | Late embryogenesis abundant (LEA) hydroxyproline-rich glycoprotein family | | 10,96 ± 3,48 | 0,34 ± 2,16 | 2,1 ± 2,16 | 5,55E-17 ± 7,55E-33 |
| At4g14650 | Unknown protein | | 44,14 ± 7,62 | 11,64 ± 5,35 | 11,83 ± 5,35 | 6,53 ± 1,02 |
| At1g51260 | LPAT3, lysophosphatidyl acyltransferase 3 | *LYSOPHOSPHATIDYL ACYLTRANSFERASE 3 (LPAT3)* | 8,22 ± 2,68 | 0,63 ± 7,55E-33 | 5,55E-17 ± 7,55E-33 | 1,6 ± 2,58 |
| At3g13840 | GRAS family transcription factor | | 27,8 ± 8,14 | 4,52 ± 0,91 | 4,62 ± 0,91 | 5,55E-17 ± 7,55E-33 |
| At5g66280 | GMD1, GDP-D-mannose 4,6-dehydratase 1 | *GDP-D-MANNOSE 4,6-DEHYDRATASE 1 (GMD1)* | 384,67 ± 22,91 | 111,87 ± 32,83 | 302,69 ± 32,83 | 162,02 ± 40,29 |
| At4g35160 | O-methyltransferase family protein | | 207,64 ± 29,77 | 83,53 ± 0,72 | 21,91 ± 0,72 | 24,81 ± 8,56 |
| At1g63450 | RHS8, root hair specific 8 | *ROOT HAIR SPECIFIC 8 (RHS8)* | 56,73 ± 7,74 | 7,57 ± 4,26 | 12,15 ± 4,26 | 7,72 ± 3,8 |
| At2g43050 | ATPMEPCRD, Plant invertase/pectin methylesterase inhibitor superfamily | *(ATPMEPCRD)* | 67,9 ± 11,17 | 16,4 ± 0,58 | 28,57 ± 0,58 | 7,77 ± 2,62 |
| At5g22500 | FAR1, fatty acid reductase 1 | *FATTY ACID REDUCTASE 1 (FAR1)* | 611,43 ± 57,42 | 249,49 ± 62,47 | 235,07 ± 62,47 | 218,41 ± 17,83 |
| At1g33800 | Protein of unknown function (DUF579) | | 267,74 ± 26,81 | 143,16 ± 11,63 | 162,17 ± 11,63 | 165,96 ± 18,59 |
| At1g30750 | Unknown protein | | 141,08 ± 7,41 | 39,96 ± 9,67 | 84,31 ± 9,67 | 35,47 ± 5,97 |
| At4g08950 | EXO, Phosphate-responsive 1 family protein | *EXORDIUM (EXO)* | 865,82 ± 244,1 | 222,15 ± 34,38 | 443,15 ± 34,38 | 387,78 ± 77,41 |
| At3g19320 | Leucine-rich repeat (LRR) family protein | | 26,45 ± 6,73 | 5,91 ± 0,66 | 4,22 ± 0,66 | 3,46 ± 3,61 |
| At1g08592 | Potential natural antisense gene, locus overlaps with AT1G08590 | | 343,89 ± 6,82 | 99,11 ± 21,55 | 220,64 ± 21,55 | 103,72 ± 5,1 |
| At3g60280 | UCC3, uclacyanin 3 | *UCLACYANIN 3 (UCC3)* | 55,02 ± 7,29 | 1,7 ± 2,79 | 19,64 ± 2,79 | 0,83 ± 1,35 |
| At4g31910 | HXXXD-type acyl-transferase family protein | | 378,16 ± 67,45 | 16,1 ± 31,1 | 143,28 ± 31,1 | 32 ± 7,38 |
| At1g18075 | MIR159, MIR159B, MIR159/MIR159B; miRNA | *MICRORNA159B (MIR159B)* | 41,15 ± 5,03 | 21,06 ± 0,96 | 30,7 ± 0,96 | 28,44 ± 5,3 |

**Supplementary Table S1. UV-B responsive genes regulated by TGA2/5/6 factors.** List of genes differentially regulated by the interaction between treatment and genotype.

| Locus | Gene model Description | Gene Symbol | Normalized expression | | | |
|---|---|---|---|---|---|---|
| | | | *Wild type* | | *tga256* | |
| | | | Control | UV-B | Control | UV-B |
| | ***Cluster 5*** | | | | | |
| At3g63470 | scpl40, serine carboxypeptidase-like 40 | *SERINE CARBOXYPEPTIDASE-LIKE 4 (scpl4)* | 110,08 ± 7,44 | 29,1 ± 14,85 | 50,5 ± 14,85 | 29,53 ± 1,89 |
| At5g57530 | AtXTH12, XTH12, xyloglucan endotransglucosylase/hydrolase 12 | *XYLOGLUCAN ENDOTRANSGLUCOSYLASE/HYDROLASE 12 (XTH12)* | 123,85 ± 7,95 | 12,55 ± 9,69 | 40,9 ± 9,69 | 11,24 ± 3,39 |
| At5g14180 | MPL1, Myzus persicae-induced lipase 1 | *MYZUS PERSICAE-INDUCED LIPASE 1 (MPL1)* | 64,2 ± 6,94 | 8,12 ± 10,14 | 9,5 ± 10,14 | 9,64 ± 13,9 |
| At4g39350 | ATCESA2, ATH-A, CESA2, cellulose synthase A2 | *CELLULOSE SYNTHASE A2 (CESA2)* | 2165,47 ± 160,18 | 1105,21 ± 144,13 | 1456,2 ± 144,13 | 970,57 ± 43,93 |
| At3g45110 | Unknown protein | | 0,9 ± 0,4 | 5,55E-17 ± 7,55E-33 | 5,55E-17 ± 7,55E-33 | 5,55E-17 ± 7,55E-33 |
| At5g56320 | ATEXP14, ATEXPA14, ATHEXP ALPHA 1.5, EXP14, EXPA14, expansin A14 | *EXPANSIN A14 (EXPA14)* | 218,19 ± 2,59 | 24,51 ± 8,48 | 109,98 ± 8,48 | 31,77 ± 1,7 |
| At3g59850 | Pectin lyase-like superfamily protein | | 22,66 ± 2,91 | 2,73 ± 4,3 | 9,36 ± 4,3 | 3,53 ± 1,99 |
| At4g30420 | nodulin MtN21 /EamA-like transporter family protein | | 22,71 ± 1,44 | 6,46 ± 2,54 | 8,75 ± 2,54 | 3,89 ± 2,2 |
| At5g15290 | Uncharacterised protein family (UPF0497) | *CASPARIAN STRIP MEMBRANE DOMAIN PROTEIN 5 (CASP5)* | 13,83 ± 5,31 | 0,71 ± 1,94 | 1,67 ± 1,94 | 5,55E-17 ± 7,55E-33 |
| At3g55230 | Disease resistance-responsive (dirigent-like protein) family protein | | 409,81 ± 28,59 | 81,92 ± 28,65 | 272,46 ± 28,65 | 112,55 ± 2,66 |
| At3g51770 | ATEOL1, ETO1, tetratricopeptide repeat (TPR)-containing protein | *ETHYLENE OVERPRODUCER 1 (ETO1)* | 532,02 ± 25,56 | 422,44 ± 30,1 | 537,19 ± 30,1 | 542,98 ± 28,88 |
| At5g42840 | Cysteine/Histidine-rich C1 domain family protein | | 38,42 ± 9,42 | 8,37 ± 2,95 | 7,86 ± 2,95 | 3,08 ± 0,85 |
| At3g52900 | Family of unknown function (DUF662) | | 105,15 ± 15,97 | 42,77 ± 2,58 | 82,79 ± 2,58 | 68,41 ± 10,59 |
| At2g04090 | MATE efflux family protein | | 27,2 ± 2,91 | 2,67 ± 1,41 | 8,19 ± 1,41 | 0,89 ± 0,73 |
| At1g54450 | Calcium-binding EF-hand family protein | | 73,96 ± 9,59 | 35,35 ± 2,23 | 50,96 ± 2,23 | 42,13 ± 9,33 |
| At1g05660 | Pectin lyase-like superfamily protein | | 61,32 ± 7,75 | 3,11 ± 3,95 | 12,63 ± 3,95 | 2,18 ± 1,33 |
| At2g04800 | Unknown protein | | 80,64 ± 13,21 | 22,9 ± 8,67 | 17,26 ± 8,67 | 9,9 ± 3,58 |
| At5g40730 | AGP24, ATAGP24, arabinogalactan protein 24 | *ARABINOGALACTAN PROTEIN 24 (AGP24)* | 573,05 ± 121,29 | 89,05 ± 20,96 | 293,99 ± 20,96 | 116,45 ± 27,28 |

**Supplementary Table S1. UV-B responsive genes regulated by TGA2/5/6 factors.** List of genes differentially regulated by the interaction between treatment and genotype.

| Locus | Gene model Description | Gene Symbol | Normalized expression | | | |
|---|---|---|---|---|---|---|
| | | | *Wild type* | | *tga256* | |
| | | | Control | UV-B | Control | UV-B |
| | ***Cluster 5*** | | | | | |
| At5g46370 | ATKCO2, ATTPK2, KCO2, Ca2+ activated outward rectifying K+ channel 2 | *CA2+ ACTIVATED OUTWARD RECTIFYING K+ CHANNEL 2 (KCO2)* | 9,08 ± 2,69 | 5,55E-17 ± 1,25 | 0,73 ± 1,25 | 0,87 ± 1,42 |
| At4g28530 | anac074, NAC074, NAC domain containing protein 74 | *NAC DOMAIN CONTAINING PROTEIN 74 (NAC74)* | 38,59 ± 6,6 | 6,1 ± 3,04 | 4,11 ± 3,04 | 3,52 ± 0,85 |
| At4g22460 | Bifunctional inhibitor/lipid-transfer protein/seed storage 2S albumin superfamily protein | | 54,31 ± 18,41 | 18,98 ± 4,07 | 4,79 ± 4,07 | 7,37 ± 2,1 |
| At5g13990 | ATEXO70C2, EXO70C2, exocyst subunit exo70 family protein C2 | *EXOCYST SUBUNIT EXO7 FAMILY PROTEIN C2 (EXO7C2)* | 106,62 ± 3,66 | 32,03 ± 13,36 | 39,45 ± 13,36 | 28,37 ± 11,99 |
| At5g35940 | Mannose-binding lectin superfamily protein | | 105,27 ± 28,69 | 16,11 ± 8,56 | 38,56 ± 8,56 | 18,8 ± 9,43 |
| At5g65090 | BST1, DER4, MRH3, DNAse I-like superfamily protein | *BRISTLED 1 (BST1)* | 45 ± 0,87 | 15,43 ± 0,97 | 18,01 ± 0,97 | 20,24 ± 7,54 |
| At1g05570 | ATGSL06, ATGSL6, CALS1, GSL06, GSL6, callose synthase 1 | *CALLOSE SYNTHASE 1 (CALS1)* | 1824,39 ± 144,82 | 871,94 ± 294,2 | 1770,25 ± 294,2 | 1542,04 ± 119,23 |
| At3g05150 | Major facilitator superfamily protein | | 69,94 ± 1,93 | 3,78 ± 7,45 | 41,22 ± 7,45 | 13,77 ± 5,38 |
| At2g43040 | NPG1, tetratricopeptide repeat (TPR)-containing protein | *NO POLLEN GERMINATION 1 (NPG1)* | 127,67 ± 11,76 | 48,38 ± 7,03 | 111,42 ± 7,03 | 69,95 ± 6,59 |
| At1g67510 | Leucine-rich repeat protein kinase family protein | | 228,74 ± 9,79 | 103,55 ± 24,33 | 232,69 ± 24,33 | 167,76 ± 14,09 |
| At1g70885 | pseudogene, similar to major latex-like protein | | 17,93 ± 4,53 | 1,69 ± 2,69 | 2,79 ± 2,69 | 5,55E-17 ± 7,55E-33 |
| At4g08620 | SULTR1;1, sulphate transporter 1;1 | *SULPHATE TRANSPORTER 1;1 (SULTR1;1)* | 36,46 ± 17,58 | 5,06 ± 1,42 | 1,35 ± 1,42 | 2,91 ± 2,82 |
| At2g18800 | ATXTH21, XTH21, xyloglucan endotransglucosylase/hydrolase 21 | *XYLOGLUCAN ENDOTRANSGLUCOSYLASE/HYDROLASE 21 (XTH21)* | 48,95 ± 13,79 | 5,61 ± 1,95 | 9,09 ± 1,95 | 6,12 ± 2,5 |
| At1g64990 | GTG1, GPCR-type G protein 1 | *GPCR-TYPE G PROTEIN 1 (GTG1)* | 450,14 ± 25,18 | 321,72 ± 25,01 | 380,32 ± 25,01 | 405,72 ± 35,55 |
| At1g60450 | AtGolS7, GolS7, galactinol synthase 7 | *GALACTINOL SYNTHASE 7 (GolS7)* | 2,32 ± 0,89 | 5,55E-17 ± 7,55E-33 | 5,55E-17 ± 7,55E-33 | 5,55E-17 ± 7,55E-33 |

**Supplementary Table S1. UV-B responsive genes regulated by TGA2/5/6 factors.** List of genes differentially regulated by the interaction between treatment and genotype.

| Locus | Gene model Description | Gene Symbol | Normalized expression | | | |
|---|---|---|---|---|---|---|
| | | | *Wild type* | | *tga256* | |
| | | | Control | UV-B | Control | UV-B |
| | ***Cluster 5*** | | | | | |
| At5g49270 | COBL9, DER9, MRH4, SHV2, COBRA-like extracellular glycosyl-phosphatidyl inositol-anchored protein family | *SHAVEN 2 (SHV2)* | 172,14 ± 27,1 | 11,29 ± 12,42 | 72,7 ± 12,42 | 13,68 ± 6,59 |
| At5g03310 | SAUR-like auxin-responsive protein family | | 17,71 ± 5,64 | 1,02 ± 1,47 | 1,71 ± 1,47 | 5,55E-17 ± 7,55E-33 |
| At4g36240 | GATA7, GATA transcription factor 7 | *GATA TRANSCRIPTION FACTOR 7 (GATA7)* | 67,56 ± 9,88 | 13,76 ± 1,83 | 29,74 ± 1,83 | 10,92 ± 2,07 |
| At4g34880 | Amidase family protein | | 30,32 ± 3,09 | 20,69 ± 2,61 | 4,84 ± 2,61 | 12,96 ± 3,54 |
| At3g07070 | Protein kinase superfamily protein | | 93,88 ± 3,36 | 9,35 ± 6,01 | 45,57 ± 6,01 | 4,78 ± 2,77 |
| At5g57770 | Plant protein of unknown function (DUF828) with plant pleckstrin homology-like region | | 221,9 ± 39,89 | 25,36 ± 7,64 | 96,19 ± 7,64 | 17,21 ± 5,55 |
| At3g59370 | Vacuolar calcium-binding protein-related | | 104,75 ± 27,43 | 6,49 ± 4,07 | 37,63 ± 4,07 | 3,5 ± 1,34 |
| At5g41080 | PLC-like phosphodiesterases superfamily protein | *GLYCEROPHOSPHODIESTER PHOSPHODIESTERASE 2 (GDPD2)* | 274,86 ± 80,73 | 111,53 ± 33,57 | 209,39 ± 33,57 | 193,63 ± 11,67 |
| At4g32790 | Exostosin family protein | | 149,27 ± 4,23 | 72,86 ± 15,57 | 89,84 ± 15,57 | 83,33 ± 14,79 |
| At2g24800 | Peroxidase superfamily protein | | 4,44 ± 0,63 | 5,55E-17 ± 1,28 | 1,48 ± 1,28 | 0,44 ± 0,72 |
