## Supplementary material for "Transcription factor TGA2 is essential for UV-B stress tolerance controlling oxidative stress in Arabidopsis": Table S2

**Table S2. Primers used for cloning, ChIP and RT-qPCR assay**

| Gene/allele | AGI | Use | Direction | Sequence |
|---|---|---|---|---|
| *GRXC9* | AT1G28480 | RT-qPCR | Forward | 5’- CACTCCAAGTCCAAGAAGCAG - 3’ |
| | | | Reverse | 5’- AGAGAGTTCGGATGGTGGTG - 3’ |
| *YLS8* | AT5G08290 | RT-qPCR | Forward | 5'- TTACTGTTTCGGTTGTTCTCCATTT -3’ |
| | | | Reverse | 5'- CACTGAATCATGTTCGAAGCAAGT -3’ |
| *PR-1* | AT2G14610 | RT-qPCR | Forward | 5'- ACACGTGCAATGGAGTTTGTGG -3' |
| | | | Reverse | 5'- TTGGCACATCCGAGTCTCACTG -3' |
| *CHS* | AT5G13930 | RT-qPCR | Forward | 5'- TTCCGCATCACCAACAGTGAAC -3' |
| | | | Reverse | 5'- CGCACATGCGCTTGAACTTCTC -3' |
| *GSTU7* | AT2G29420 | ChIP | Forward | 5' - TCTTCCGATGTGGGACAAAGTG - 3' |
| | | | Reverse | 5' - ACTCGCCCACATTCCCAAAAG - 3' |
| | | RT-qPCR | Forward | 5' - TGTGACGGCGATGAAAGTTGTG - 3' |
| | | | Reverse | 5' - AATCTCTCGTCGCTTCAACCACAG - 3' |
| | | Cloning | Forward | 5' - CACCATGGCGGAGAGATCAAATTCA - 3' |
| | | | Reverse | 5' - AGCAGATTTGATATTGAG - 3' |
| *GSTU8* | AT3G09270 | ChIP | Forward | 5' - ACAGGCCTTCAACCACTACC - 3' |
| | | | Reverse | 5' - TCCCTTTGTGTGTGTGTGTT - 3' |
| | | RT-qPCR | Forward | 5' - CACAAAGGGAAAGCCAAACCGG - 3' |
| | | | Reverse | 5' - CGTTTACGTGCTCTTCTTGGTTCA - 3' |
| *GSTU25* | AT1G17180 | ChIP | Forward | 5'-TTTTGGTAATGTATAACCCCTTGA -3' |
| | | | Reverse | 5'-TGATTAAACATTTTCTTTTCATTAGC -3' |
| | | RT-qPCR | Forward | 5'- TCAGCATTGAAGCCGAGTGTCC -3' |
| | | | Reverse | 5'- TAGCCACACTCTCTCTCTCCACAC -3' |
| *ACT2* | AT3G18780 | ChIP | Forward | 5'- CTTGCACCAAGCAGCATGAA - 3' |
| | | | Reverse | 5' - CCGATCCAGACACTGTACTTCCTT - 3' |
| *TGA2* | AT5G06950 | Cloning | Forward | 5'- caccATGGCTGATACCAGTCC - 3' |
| | | | Reverse | 5' - CTCTCTGGGTCGAGCAAG - 3' |
